# Cell-Type-Resolved Transcriptomics Defines Stable and Accessible Markers of the Cardiac Purkinje Fiber in Sheep and Human Translation

**DOI:** 10.64898/2026.08.21.746241

**Authors:** Sabine Charron-Guitoger, Nestor Palleres-Lupon, Marion Constantin, Jason D. Bayer, Philippe Pasdois, Fanny Vaillant, Richard D. Walton

**Affiliations:** IHU Liryc, University of Bordeaux, INSERM, CRCTB, U1045, Bordeaux, France; IHU Liryc, University of Bordeaux, CNRS, Bordeaux INP, IMB, UMR 5251, Bordeaux, France; IBGC, University of Bordeaux, CNRS, UMR 5095, Bordeaux, France

**Author notes:** **Corresponding author:** Richard D. Walton Avenue du Haut Lévêque Hopital Xavier Arnozan – IHU-Liryc 33604 Pessac France.

**Keywords:** Purkinje fibers, cardiac conduction system, transcriptomics, laser-capture microdissection, cell-type-specific markers

## Abstract

**Background:** The His-Purkinje network drives rapid ventricular activation and is a major substrate for ventricular arrhythmias, yet it is among the least molecularly characterized cardiac compartments. Markers validated in rodents transfer poorly across species, few are confirmed at the protein level in large mammals or humans, and most lack the stability and surface accessibility that demanding applications require.

**Methods:** We combined histology-guided laser-capture microdissection with low-input, cell-type-resolved RNA-sequencing to profile Purkinje fibers, left-ventricular cardiomyocytes and peri-Purkinje stroma from adult sheep. Differentially expressed genes were ranked by a transparent composite framework weighting expression specificity, cross-individual stability and predicted subcellular accessibility; leading candidates were validated by RT-qPCR and immunolabelling in sheep and by RT-qPCR in human myocardium.

**Results:** RNA-sequencing resolved a Purkinje transcriptome distinct from cardiomyocytes and stroma and defined 331 concordantly enriched genes, which the composite framework ranked into stable, specific candidates spanning intracellular and cell-surface compartments. By RT-qPCR, the canonical conduction markers connexin-40/*GJA5*, *HCN4*, *NEFM* and *MYL4* were strongly enriched in Purkinje fibers, whereas the rodent gold-standard contactin-2 was not, underscoring species divergence. Thirteen of sixteen prioritized candidates were confirmed by RT-qPCR, and immunolabelling localized *MYL4*, *CNN1*, *TAGLN* and *DKK3* to Purkinje fibers; contactin-5 emerged as a novel transcript-and protein-validated Purkinje marker. In human myocardium, a defined subset - *MYL4*, connexin-40/*GJA5*, contactin-5 and *TAGLN* - was conserved, while several markers proved species-restricted.

**Conclusions:** We provide the first genome-wide, cell-type-resolved molecular portrait of the Purkinje fiber in a large-animal model and a generalizable strategy that selects markers for specificity, stability and accessibility. The resulting resource - including the cross-species marker contactin-5 and compartment-matched candidates - supplies validated tools to identify, isolate and target Purkinje cells and demonstrates the necessity of cross-species validation.

## Introduction

The His-Purkinje network is the heart’s high-velocity electrical distribution system, converting the orderly atrioventricular impulse into the rapid, near-synchronous activation of the ventricles ^1^. Beyond this physiological role, the Purkinje fiber (PF) network has emerged as a pivotal contributor to disease. These specialized fast conducting cells generate much of the ectopic and re-entrant activity that triggers and sustains idiopathic and ischemic ventricular fibrillation and polymorphic ventricular tachycardia, and they are an increasingly important substrate and target for catheter ablation of otherwise refractory ventricular arrhythmias ^2–7^. A detailed molecular understanding of the PF, together with the means to identify, isolate and selectively target it is therefore of considerable basic and clinical importance.

Despite this prominence, the Purkinje network remains among the most poorly characterized compartments of the heart. PFs are sparse, structurally continuous with the working myocardium and difficult to delineate without specialized histology, which has long impeded their molecular dissection. The molecular and developmental definition of the cardiac conduction system has been established predominantly in mouse, with additional formative contributions from avian and canine studies ^8–14^. Human conduction-system anatomy and developmental patterning were described earlier by classical embryologic, molecular, and high-resolution anatomical studies, and have been further refined recently by single-cell and spatial transcriptomic atlases ^15,16^. Crucially, markers and reporter strategies validated in rodents do not transfer reliably across species or developmental stages, and very few have been confirmed at the protein level in large mammals or in human tissue. The field consequently lacks a robust, cross-species toolkit for recognizing the PF, and the molecular landscape of the network is largely uncharted in precisely the settings most relevant to translation.

At the root of this deficit lies a shortage of accessible and reliable molecular targets. A useful Purkinje marker must be not only specific to the fiber but stable, faithful to the tissue across ageing and disease states in which it would be applied, and, for the most demanding applications such as viable-cell isolation, molecular imaging or targeted delivery, which often relies on cell surface accessibility. The few established markers fall short on one or more of these axes: the gap-junction protein connexin-40 (encoded by gene GJA5) is shared with the atria, the canonical adhesion marker contactin-2 (*CNTN2*) is not uniformly conserved across species ^13^, and several otherwise informative markers are intracellular and therefore unsuited to surface-based targeting. This gap is most acute where it matters most for translation - in the large-mammalian models that approximate human cardiac anatomy and electrophysiology, and in human tissue itself. The sheep, whose large free-running Purkinje false tendons and human-like ventricular electrophysiology have made it a workhorse of translational arrhythmia and device research^17–19^, has to date no dedicated Purkinje transcriptome or validated, accessible marker set. Two developments now make it possible to address this gap directly. First, histology-guided laser-capture microdissection (LCM) permits the clean isolation of PFs alongside their immediate neighbors, yielding cell-type-resolved transcriptomes that distinguish genuine Purkinje enrichment from contamination by adjacent cardiomyocytes and stroma. Second, and equally important, such data can now be interrogated far beyond differential expression. The wealth of curated knowledge linking each gene to its signaling pathways, protein subcellular localization, structural context, regulatory burden and disease associations can be integrated *in silico* to prioritize candidates not merely by how strongly they are enriched, but by how stable and how accessible their products are likely to be. Exploiting this methodologically rich, complementary bioinformatic dimension - rather than treating the transcriptome as an end in itself - offers a route to markers selected for the very properties that determine their real-world utility, and represents an opportunity that conventional fold-change-based screens leave untapped.

We therefore set out to define a robust, cell-type-specific molecular signature of cardiac PFs in a large-animal (ovine) model by resolving the Purkinje transcriptome against working left-ventricular cardiomyocytes and the surrounding stroma, and to prioritize candidate markers using a framework that weights expression specificity together with cross-individual stability and predicted subcellular accessibility, before validating the leading candidates at the transcript and protein levels and assessing their conservation in human myocardium. We hypothesized that Purkinje fibers possess a distinct and reproducible transcriptional identity, separable from neighboring cardiomyocytes and stroma, that encodes candidate marker genes which are both specifically enriched and stably expressed across individuals; and that prioritizing candidates on specificity together with stability and accessibility - rather than on enrichment alone - would nominate markers that validate at the protein level and are conserved in the human conduction system.

## Materials and Methods

### Data availability

The RNA-sequencing data sets are in accordance with the MIAME guidelines and can be made accessible for GEO submission, where necessary. All other data, analytic methods and study materials are available from the corresponding author upon reasonable request, and a Major Resources Table is provided in the Data Supplement.

### Ethics statements

All procedures used in this study complied with European Parliament directive 2010/63/EU, which regulates the protection of animals, used for scientific purposes, and were approved by the local ethics committee of the University of Bordeaux (CEEA-050) and the French government (approval no. A33-318-3). Human hearts were obtained in collaboration with the University Hospital of Bordeaux in accordance with the Declaration of Helsinki. A dedicated research program approved by the French Biomedicine Agency (PFS14-004) procures heart organs from brain-dead patients whose hearts could not be used for transplantation, who had not expressed their opposition to organ donation during their lifetime, and whose families gave written informed consent. A brief description of each patient heart donor are listed in the Data Supplement (**Table S1**).

### Study design

The experimental workflow is summarized in **Figure 1** and a detailed methods section can be found in the supplemental materials.

**Figure 1.**
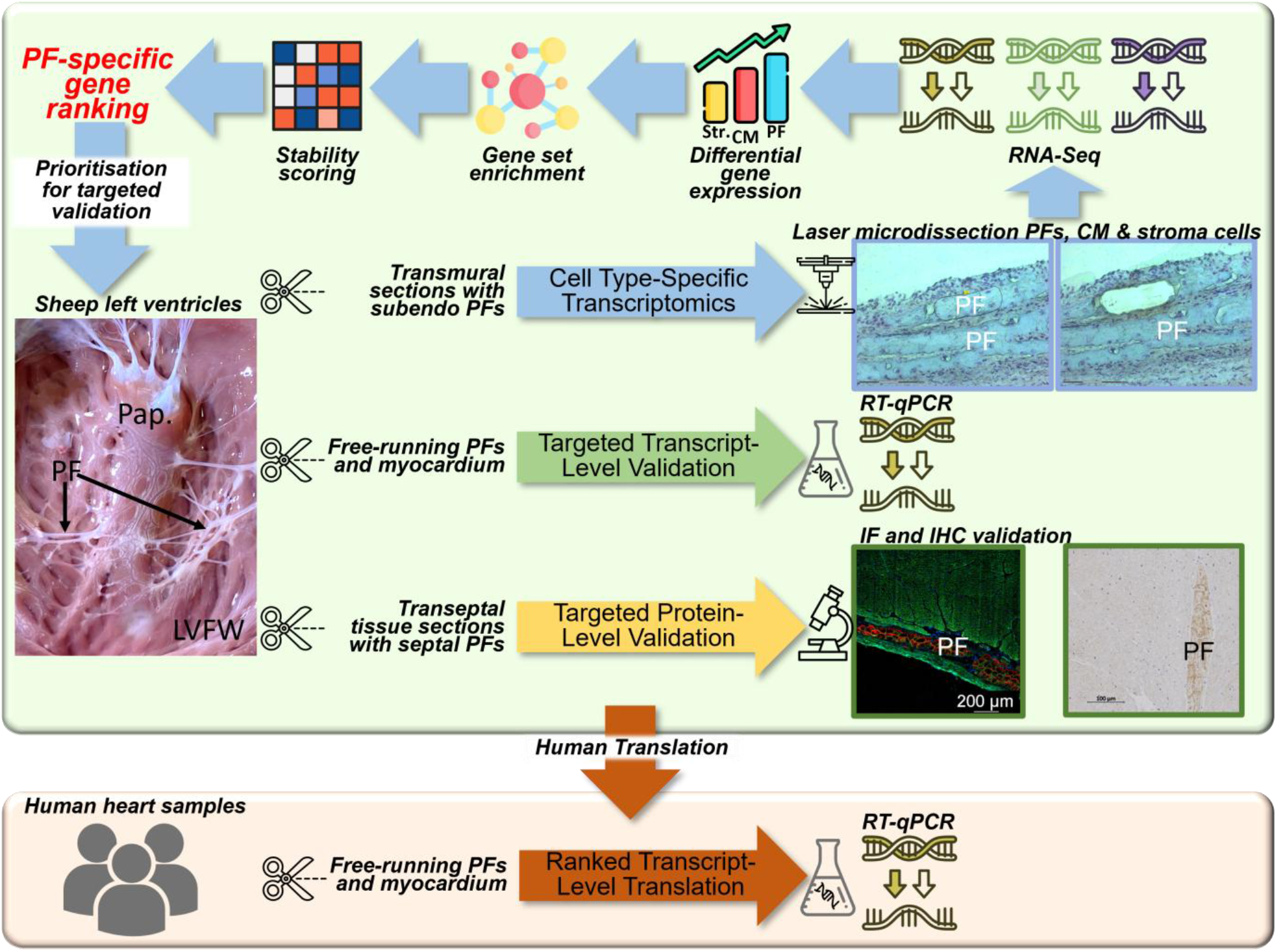
Study design and analytical workflow. Histology-guided LCM of three cell populations - free-running PFs, left-ventricular cardiomyocytes and the peri-Purkinje collagen stroma - from the ovine left ventricle, followed by low-input, cell-type-resolved RNA-sequencing, differential expression against both comparators, cross-species ortholog mapping, a multi-source composite scoring step, and targeted transcript-level (RT-qPCR) and protein-level (immunolabelling) validation, with a human RT-qPCR bridge.

### Animal handling

Hearts from adult ovine (N=10, Charmoise breed, 1-2 yrs old, 40-45 kg) were used owing to their increasing application in translational research in cardiology, and prominent PF morphology permitting clean, cell-type-resolved sampling. Anesthesia was induced with intravenous ketamine (15 mg/kg) and midazolam (1.5 mg/kg); animals were intubated, ventilated and given heparin (2.5 mg/kg), and anesthesia was maintained with ketamine (40 mg/kg/h) and midazolam (2 mg/kg/h). Animals were euthanized with intravenous sodium pentobarbital (35 mg/kg); hearts were rapidly excised, the aorta was cannulated, and the coronary circulation was flushed with cardioplegic solution, containing (mM): NaCl, 110; CaCl2, 1.2; KCl, 16; MgCL2, 16; NaHCO3, 10 and glucose, 9.01 at 4°C.

### Tissue collection

Septobasal regions of the ventricles from ovine hearts, owing to a rich density and conserved presence of PFs, were dissected from 6 ovine hearts for LCM and downstream transcriptomics, and 2 additional ovine hearts were dedicated to independent immunolabelling studies. Transeptal tissue blocks were fixed in paraformaldehyde (4%) and embedded in paraffin. For real-time quantitative polymerase chain reaction (RT-qPCR) studies, PF-containing false tendons (free-running portions of the PF network) and sub-epicardial portions of the adjacent left ventricular free wall were dissected from 7 ovine and 4 human hearts, and snap-frozen by immersion in liquid nitrogen-cooled isopentane. Frozen samples were stored at-80°C until use. Several false tendon branches dissected from ovine hearts were subject to fixation and paraffin-embedded for histology.

### RT-qPCR validation

Total RNA was extracted from Purkinje fibers and from left-ventricular myocardium using column-based kits (Supplemental Methods). Efficiency-validated primers designed from Ensembl sequences (**Table S2**) were used to reverse-transcribe and amplify each target by SYBR-based RT-qPCR on a Bio-Rad CFX96 instrument in technical triplicate. Relative quantification used the 2^(-delta-delta-Ct) method, normalized to the geometric mean of two reference genes (*HPRT1* and *B2M* in sheep; *B2M* and *GUSB* in human), and Purkinje fibers were compared with cardiomyocytes (n = 4 biological replicates per group) as described under Statistical analysis. For *CNTN5*, amplification in several ovine cardiomyocyte samples fell at or below the reliably measurable cycle window, so cardiomyocyte expression for that gene is reported as at or below the limit of detection. Extraction, reagent and thermal-cycling details are provided in the Supplemental Methods.

### Immunofluorescence and immunohistochemistry

Immunofluorescence and immunohistochemistry were performed on 6-micrometre paraffin sections (Leica RM2255) with heat-induced epitope retrieval (citrate buffer, pH 6.0, 98 degrees C, 20 min), peroxidase and serum blocking. A full listing of antibodies and detection reagents are found in the Data Supplement (**Table S3**). Briefly, for immunuofluorescence, primary antibodies were tested against cardiac troponin T (*cTNT*), connexin-40 (*GJA5*), *MYL4* and *CNTN5* and a DAPI nuclear counterstain. Sections were imaged on a Leica THUNDER Imager with a Leica DFC9000 GTC camera. For immunohistochemistry, primary antibodies against *MYL4*, *CNN1*, sm22-alpha (*TAGLN)* and *DKK3* were applied. Detection used a biotinylated secondary antibody with avidin-biotin-peroxidase detection and 3,3-prime-diaminobenzidine, and a haematoxylin counterstain; sections were imaged using an AxioScan 7 (Zeiss, Germany). Absence-of-primary-antibody negative controls were performed for all targets.

### Free-running PF histology

For histopathological characterization of the free-running PF, formaldehyde-fixed tissue was paraffin-embedded, sectioned at 6 µm on a Leica RM2255 microtome and stained with Masson’s trichrome and periodic acid-Schiff (see supplemental materials for detailed protocols) to resolve false tendon composition (**Figure 2A**).

**Figure 2.**
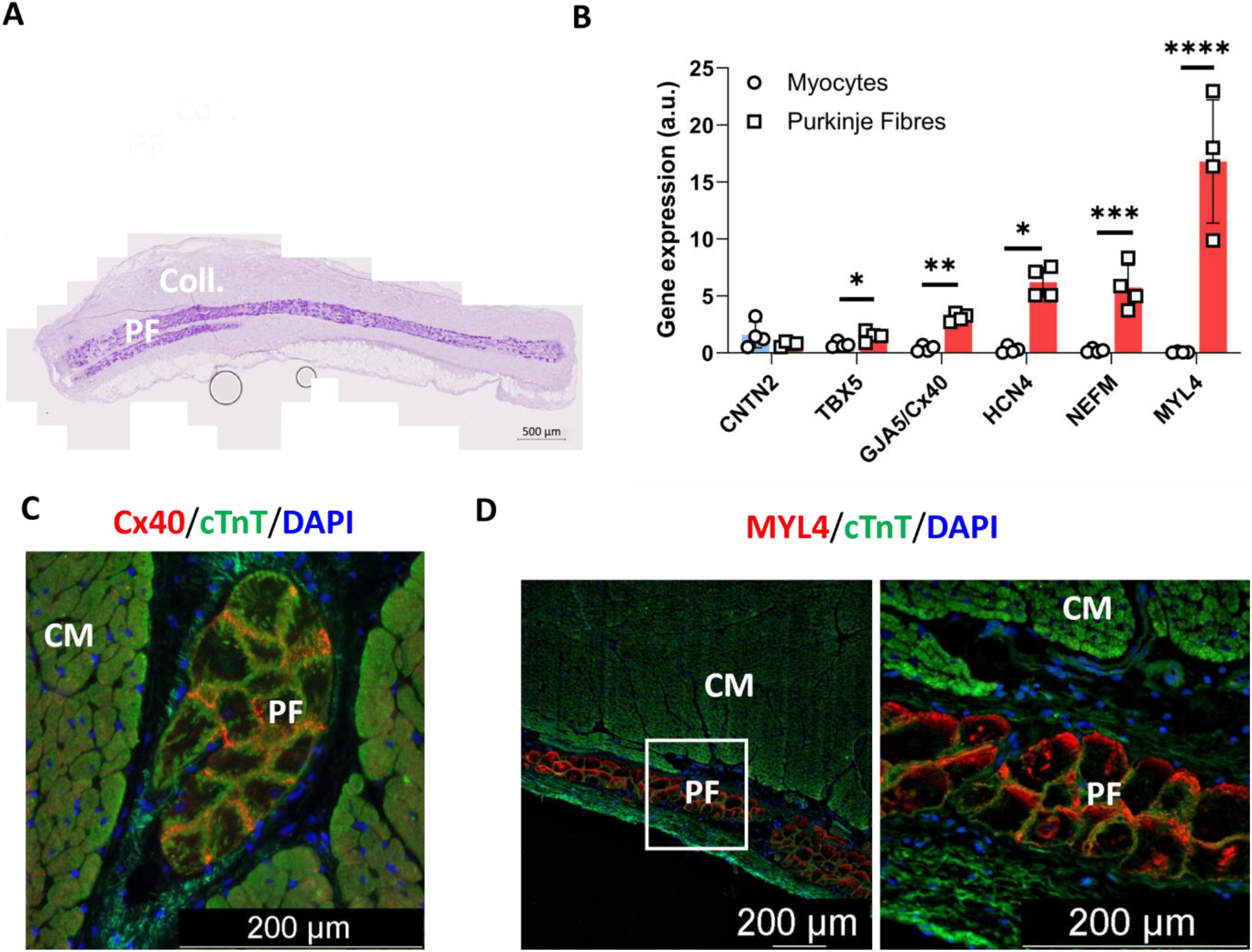
PFs recapitulate canonical conduction-system markers. (A) Representative histology (Masson’s trichrome and periodic acid-Schiff) of free-running Purkinje strands and the investing collagen sheath; scale bar 500 µm. (B) RT-qPCR of canonical markers in PFs versus cardiomyocytes: *GJA5* (connexin-40), *HCN4*, *NEFM* and *MYL4* are strongly enriched in PFs and *TBX5* is modestly enriched, whereas *CNTN2* is not enriched. Data are mean ± SD; comparison by Welch’s unequal-variance t test on log2 reference-normalized expression (reference genes *HPRT1* and *B2M* in sheep); n = 4 per group; *P < 0.05, **P < 0.01, ***P < 0.001 and **** P < 0.0001, and n.s. denotes not significant. (C, D) Immunofluorescence co-staining for cardiac troponin T (*cTnT*) with connexin-40 (C) and *MYL4* (D); PF are indicated by asterisks and the working myocardium by arrows; nuclei are counterstained with DAPI; scale bars 200 µm.

**Figure 3.**
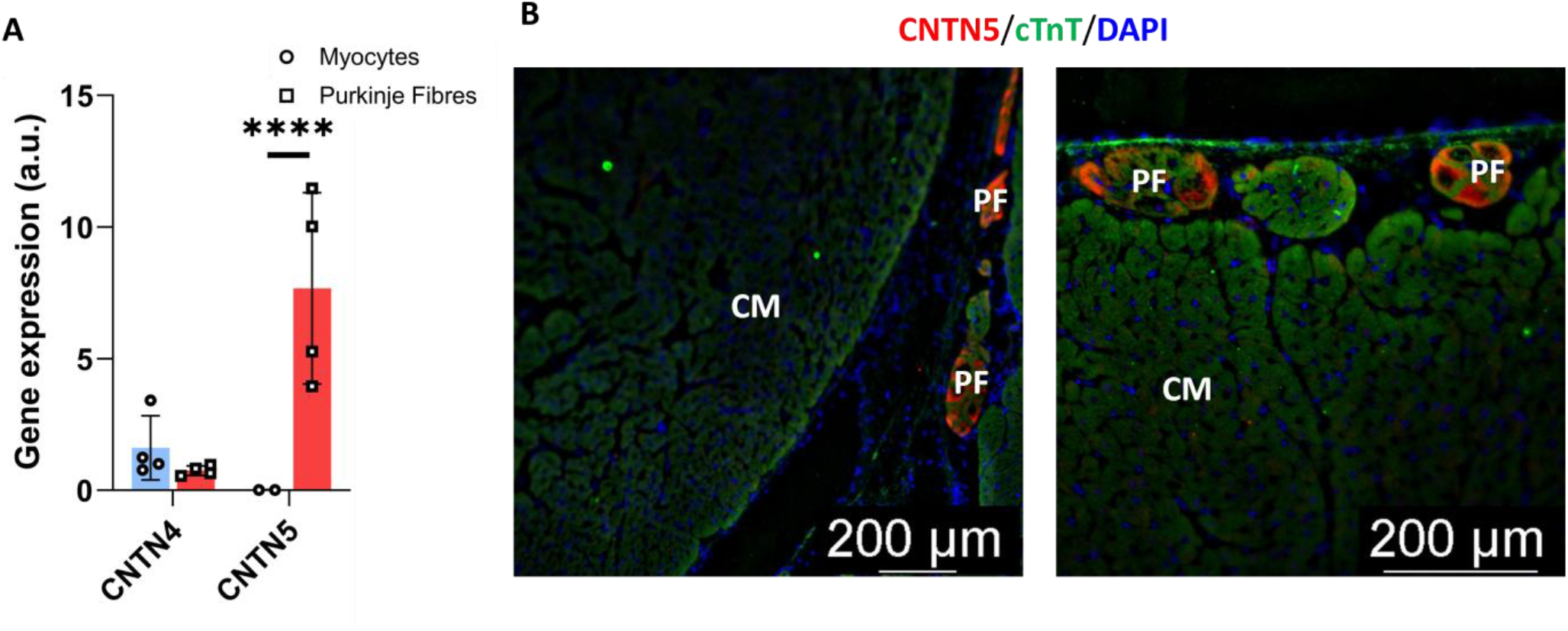
Contactin-5 is a transcript-and protein-validated Purkinje marker. (A) RT-qPCR of *CNTN4* and *CNTN5* (PFs versus cardiomyocytes): *CNTN5* is strongly enriched in PFs, with cardiomyocyte expression at or below the limit of detection, whereas *CNTN4* is not enriched. Data are mean +/-SD; statistics as in **Figure 2**; with n = 4 per group (for *CNTN5*, two cardiomyocyte replicates were quantifiable, the remainder being at or below the limit of detection). (B, C) *CNTN5* immunofluorescence (with cTnT and DAPI) is restricted to PFs (asterisks) and absent from the surrounding cardiomyocytes (arrows); scale bars 200 µm.

### Laser-capture microdissection

Laser-capture microdissection (LCM) and RNA-sequencing were performed at the PUMA core facility (Neurocentre Magendie, INSERM U1215, University of Bordeaux). From PF-rich transeptal sections of the ventricles, three cell populations - Purkinje fibers, left-ventricular cardiomyocytes and stroma of the peri-Purkinje collagen sheath - were identified by cresyl-violet histological staining and captured separately from each animal for downstream RNA-sequencing. OCT-embedded, snap-frozen tissue was cryosectioned at 20 µm (Purkinje fibers, stroma) or 30 µm (cardiomyocytes) onto polyethylene naphthalate (PEN) membrane slides, cresyl-violet stained and dehydrated, and target cells were isolated on a Zeiss P.A.L.M. MicroBeam system by laser cutting and pressure catapulting. Approximately 5-8 mm² of Purkinje fibers and peri-Purkinje collagen sheath, and approximately 30 mm² of cardiomyocytes, were captured per animal. Full staining, dehydration and microdissection settings are provided in the Supplemental Methods.

### RNA isolation, library preparation and sequencing

Cells were lysed immediately after microdissection and total RNA was extracted with a column-based kit including on-column DNase treatment (Supplemental Methods). RNA quality was assessed on an Agilent 2100 Bioanalyzer; RNA integrity numbers (RIN) were 7.9-8.2 for cardiomyocytes and 5.3-7.0 for Purkinje fibers, while stromal RNA was at the limit of detection and, being derived from the same sections as the Purkinje material, was processed alongside it. To accommodate the low-input, variable-integrity material, sequencing libraries were prepared with a low-input-tolerant, template-switching chemistry incorporating unique molecular identifiers (UMIs) for 5-prime molecule counting (Takara SMART-Seq mRNA LP with UMIs) under a poly(A) capture strategy; input RNA was 0.5 ng for Purkinje fibers and stroma and 10 ng for cardiomyocytes. Dual-indexed libraries were quality-controlled and quantified by qPCR, pooled in equivalent amounts for Purkinje fibers and stroma and at twice that depth for cardiomyocytes, and sequenced at the PGTB facility ^20^ on an Illumina NextSeq 2000 (P2 XLEAP-SBS, 300 cycles) with a PhiX spike-in. Paired-end sequencing (2 x 150 base pairs) yielded a mean of approximately 24.5 million read pairs per sample (range 15.2-39.2 million). Full reagent, catalogue and cycling details are provided in the Supplemental Methods and **Tables S4** and **S5**.

### Read processing, alignment, quality control and differential expression

Sequencing data were processed with the Cogent NGS Analysis Pipeline v2.0.1 (Takara Bio), which demultiplexed reads and performed trimming, alignment with STAR ^21^ to the *Ovis aries* reference genome (assembly ARS-UI_Ramb_v2.0; Ensembl release 113), and UMI parsing with gene-level counting. Mean mapping rates were 92.3% for cardiomyocytes, 90.2% for Purkinje fibers and 81.9% for stroma. Reads sharing a UMI were collapsed to per-gene molecule counts, the count matrix was imported into RStudio, and after quality control and removal of genes with zero counts across all samples, differential expression was assessed with DESeq2 ^22^ using the Wald test with Benjamini-Hochberg control of the false-discovery rate ^23^. Two contrasts were carried forward - Purkinje fibers versus cardiomyocytes and Purkinje fibers versus stroma - with Purkinje enrichment defined by an absolute log2 fold-change greater than 1.0 and an adjusted P value below 0.01. Pipeline components, tool versions and quality-control metrics are detailed in the Supplemental Methods (**Table S5**).

### Definition of the Purkinje-enriched panel

To isolate genes enriched in PFs irrespective of the comparator, the transcripts up-regulated in PFs in both contrasts (1,795 versus cardiomyocytes and 2,276 versus stroma) were intersected and filtered for robust expression (at least 10 reads in all samples of at least one group and detection in every Purkinje replicate), yielding the final 331-gene Purkinje-enriched panel (**Figure 4D**). Of these, 286 genes carried an annotated symbol and a mappable human ortholog and were scored, while 45 were retained with differential-expression statistics only (**Table S6**).

**Figure 4.**
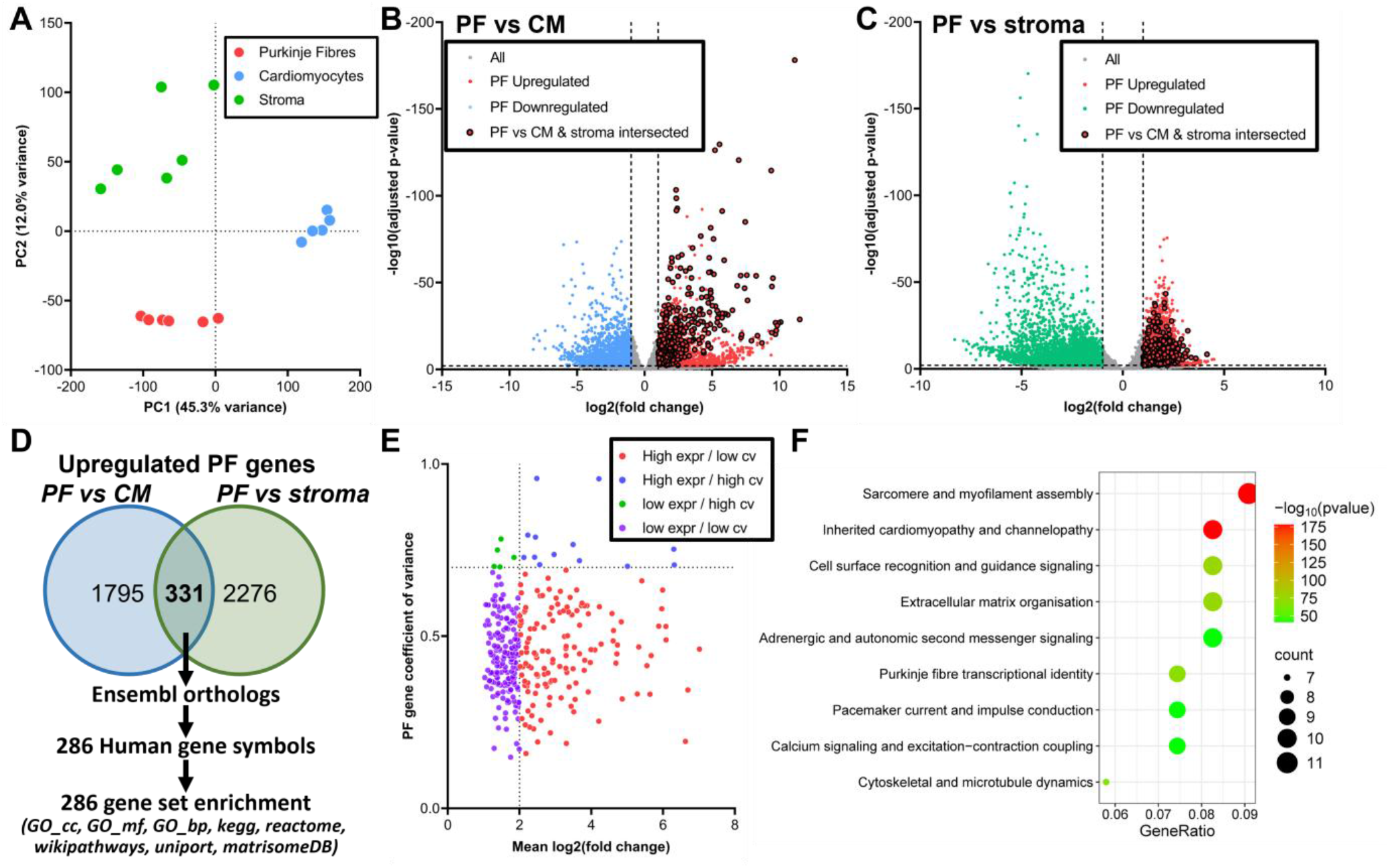
PFs possess a distinct transcriptome and a specificity-stability landscape. (A) Principal-component analysis of the three populations (PC1, 45.3%; PC2, 12.0%). (B, C) Volcano plots of differential expression for PFs versus cardiomyocytes (B) and versus stroma (C). (D) Intersection of the transcripts up-regulated in both contrasts, defining the 331-gene Purkinje-enriched panel. (E) Inter-animal coefficient of variation versus mean fold-change for the panel, partitioned into expression-and stability-defined quadrants. (F) Functional-enrichment categories of the panel. Enrichment thresholds were an absolute log2 fold-change greater than 1.0 and an adjusted P value below 0.01; n = 6 animals per population.

### Cross-species ortholog mapping

Sheep Ensembl gene identifiers were mapped to human orthologs with Ensembl BioMart ^24,25^ using the oaries_homolog_ensembl_gene attribute, querying releases 114 and 115 to maximize cross-species mapping, since sheep identifiers change across assembly releases for a substantial proportion of genes; locally cached genome-annotation files were used with programmatic enrichment. For each gene, the human symbol and Ensembl identifier, orthology type and confidence, percentage identity and the resolved Ensembl release were recorded.

### Functional annotation and prioritization framework

Genes were annotated for pathway, ontology, subcellular-localization and matrisome membership from established databases (Supplemental Methods), and prioritized by a transparent composite score combining five experimental sub-scores with five annotation-based stability pillars, each expressed as a percentile rank or normalized value across the 286 scored genes. The experimental sub-scores captured fold-change effect size, fold-change consistency across the two contrasts, statistical confidence, expression abundance and inter-animal consistency. The stability pillars quantified, from annotation-term membership, the latent propensity of each gene product to fluctuate - (A) upstream-signal convergence, (B) transcriptional fragility, (C) proteostatic vulnerability, (D) multi-pathway breadth and disease responsiveness, and (E) structural embedding - with pillars A to D treated as instability axes and inverted and pillar E applied directly. A composite stability score and a specificity score were combined with a compartment bonus reflecting predicted accessibility (cell-surface, intracellular or unknown) to give the final priority score, which ordered all 286 genes (**Figure 5C**). Database versions, the exact sub-score and pillar definitions, the 41-term pillar dictionary (**Table S7**), and all weights, thresholds and formulae are provided in the Supplemental Methods; the framework is implemented in an in-house Python suite.

**Figure 5.**
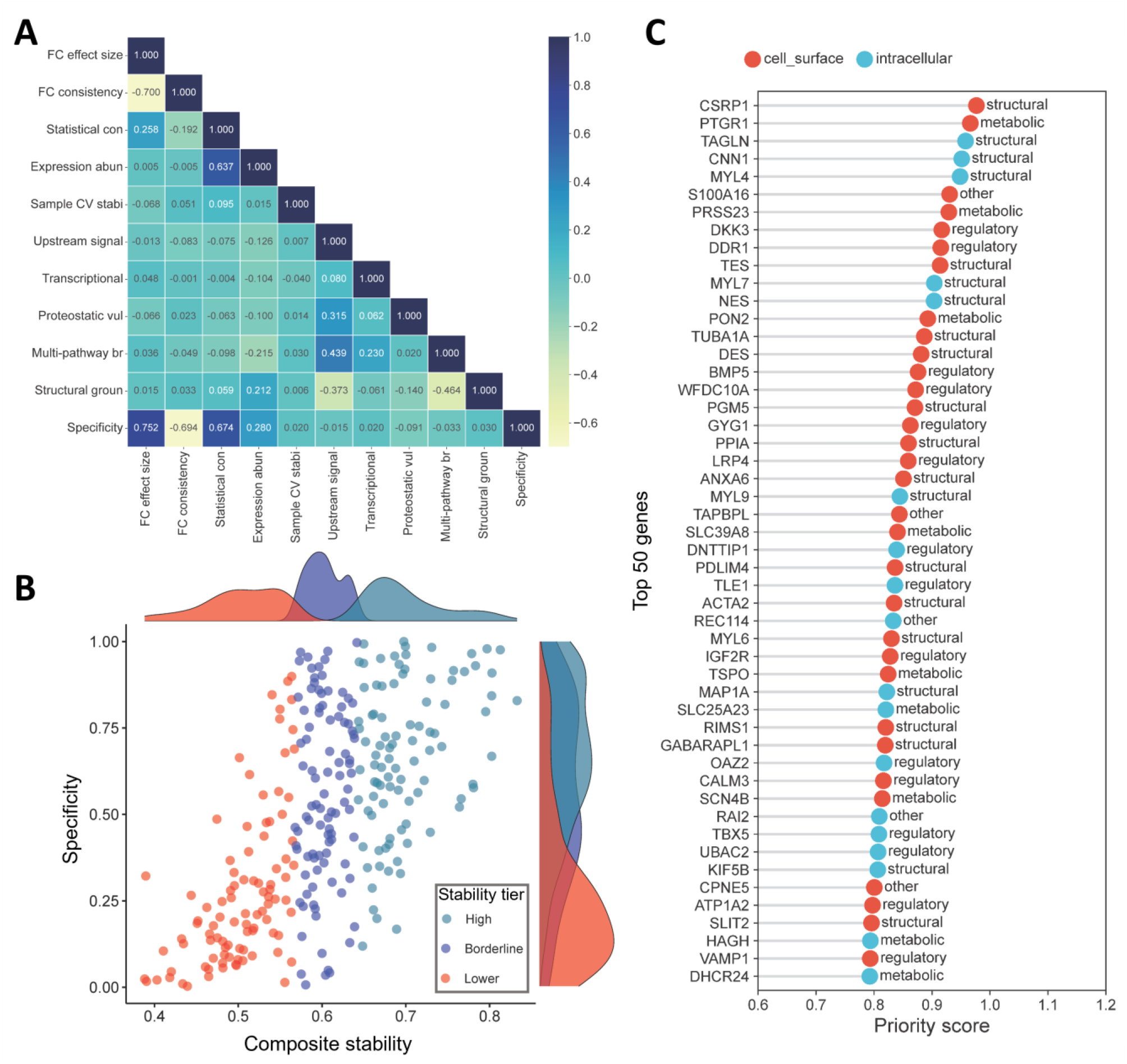
A multi-source composite framework prioritizes specific and stable candidates. (A) Pairwise correlation among the ten scoring components and the specificity score. (B) Composite stability versus specificity, with genes stratified by stability tier. (C) The fifty highest-priority candidates ranked by the final priority score (0.55 x stability percentile + 0.35 x specificity + 0.10 x compartment bonus), each annotated by predicted subcellular compartment and functional class. The priority score is a deterministic percentile-rank composite and is not an inferential test; the five stability pillars are defined in **Figure S2**.

## Statistical analysis

Differential expression was assessed with DESeq2 applied to STAR gene-level counts. Size factors were estimated by the median-of-ratios method, dispersions fitted under the negative-binomial model, and significance assessed with the Wald test; p-values were adjusted for multiple testing by the Benjamini– Hochberg procedure to control false-discovery rate, with genes at adjusted P < 0.01 and |log2 fold change| ≥ 1 considered differentially expressed. For RT-qPCR, relative expression was quantified by normalization to the geometric mean of two reference genes (*HPRT1* and *B2M* in sheep; *B2M* and *GUSB* in human), and the pre-specified primary between-group test was Welch’s unequal-variance t test applied to log2-transformed normalized expression - which is mathematically equivalent to a t test on the delta-Ct values - comparing PFs with cardiomyocytes (sheep, n = 4 per group; human, n = 4 per group), with two-sided P values. This test was applied identically to the sheep and human data for a single, consistent strategy, the choice reflecting the approximately log-normal distribution of quantitative PCR data and unequal between-group variances. As a non-parametric sensitivity analysis, two-tailed exact Mann-Whitney U tests were also computed and were concordant except for a small number of borderline genes identified in the text. Findings are described primarily in terms of fold-change, direction and reproducibility across individuals, with P values reported as supporting evidence given the modest sample sizes; the validated candidates were pre-specified, and P values are presented without formal correction for multiplicity. The prioritization score is a deterministic percentile-rank composite rather than a hypothesis test. Pathway-term frequencies across the panel were reported descriptively without inferential testing, and Fisher’s exact tests with Benjamini-Hochberg correction were applied only to pre-defined biological subgroups. Analyses used GraphPad Prism 9.5 (GraphPad Software), R (DESeq2, RNAseqQC and gage) and in-house programmatical tools written in Python.

## Results

### A laser-capture transcriptomic strategy for ovine PFs

To resolve the molecular identity of PFs against their immediate tissue neighbors, we combined histology-guided LCM with cell-type-resolved RNA-sequencing and a tiered validation pipeline (**Figure 1**). Subendocardial PFs of the basal septum, cardiomyocytes and the investing peri-Purkinje collagen stroma were micro-dissected from six adult sheep. Low-input RNA-sequencing of the three retained populations provided the discovery substrate, from which differentially expressed genes were carried through statistical filtering, network annotation and a composite scoring step to a ranked list of Purkinje-specific candidates, and finally to targeted transcript-and protein-level validation (**Figure 1**).

### Ovine Purkinje fibers recapitulate canonical conduction-system markers

We first confirmed that Purkinje cell-rich false tendons reproduced established conduction-system biology. Histological sections delineated the false tendon-residing PFs and their surrounding collagen sheath (**Figure 2A**). By RT-qPCR (PFs versus cardiomyocytes; Welch’s t test on reference-normalized expression), the canonical conduction markers *GJA5* (encoding connexin-40), *HCN4*, *NEFM* and *MYL4* were each strongly enriched in false tedons, with complete separation between groups and fold-changes ranging from roughly ten-fold for connexin-40 to more than two hundred-fold for *MYL4*, the most enriched canonical marker (all P < 0.012; **Figure 2B**); full per-gene sheep RT-qPCR statistics are provided in the Data Supplement (**Table S8**). The conduction transcription factor *TBX5* was modestly but significantly enriched (approximately two-fold; P = 0.02), whereas *CNTN2* was not enriched and not found to be significantly different to cardiomyocytes (P = 0.25; **Figure 2B**). At the protein level, immunofluorescence co-staining for cTNT localized connexin-40 (**Figure 2C**) and *MYL4* (**Figure 2D**) to the PFs, with negligible signal in the adjacent working myocardium. These data establish that the laser-capture strategy faithfully captures Purkinje identity, while already indicating that the rodent gold-standard marker *CNTN2* is not enriched in adult ovine ventricular PFs.

### Contactin-5 (*CNTN5*) is a transcript-and protein-validated Purkinje marker

Because contactins feature prominently in conduction-system biology, we examined two additional isoforms. *CNTN4* was not enriched in PFs and leaned toward the cardiomyocyte direction (P = 0.15). *CNTN5*, in contrast, was strongly enriched in PFs at the transcript level, with cardiomyocyte expression at or below the limit of detection in most samples (Welch’s t test, P < 0.001; **Figure 3A**). This transcript-level enrichment was corroborated at the protein level: *CNTN5* immunofluorescence was unambiguously restricted to PFs and absent from the surrounding cardiomyocytes (**Figure 3B** and **3C**), establishing *CNTN5* as a transcript-and protein-validated marker of Purkinje and proximal conduction tissue; full per-gene sheep RT-qPCR statistics are provided in the Data Supplement (**Table S8**).

### PFs possess a transcriptome distinct from cardiomyocytes and stroma

Unsupervised analysis of the cell-type-resolved transcriptomes separated the three populations along the first two principal components, which captured 45.3% and 12.0% of the variance; PFs, cardiomyocytes and stroma each formed a discrete cluster (**Figure 4A**). Differential-expression analysis (adjusted P < 0.01 and absolute log2 fold-change greater than 1) revealed extensive remodeling of the Purkinje transcriptome relative to each comparator (**Figure 4B**, PFs versus cardiomyocytes; **Figure 4C**, PFs versus stroma). Intersecting the transcripts up-regulated in both contrasts (1,795 versus cardiomyocytes and 2,276 versus stroma) defined a core panel of 331 genes concordantly enriched in PFs relative to both cardiomyocytes and stroma (**Figure 4D**); 286 of these mapped to annotated human orthologs and provided the basis for the subsequent cross-species analysis.

### A specificity-versus-stability landscape and the pathways of the Purkinje-enriched panel

Reasoning that a dependable marker must be not only specifically enriched but also reproducible across individuals, we plotted each panel gene by its inter-animal coefficient of variation against its mean fold-change, partitioning the panel into expression-and stability-defined quadrants (**Figure 4E**). A subset of genes combined high expression, high specificity and low variance - the profile sought for a robust marker - whereas others were specific but more variable. Functional-enrichment analysis of the panel returned categories congruent with Purkinje biology, including sarcomere and myofilament assembly, inherited cardiomyopathy and channelopathy, cell-surface recognition and guidance signaling, extracellular-matrix organization, adrenergic and autonomic second-messenger signaling, PF transcriptional identity, pacemaker current and impulse conduction, calcium signaling and excitation-contraction coupling, and cytoskeletal and microtubule dynamics (**Figure 4F**), with the panel’s subcellular-localization, molecular-function and disease-association breakdown provided in **Figure S1**.

### A composite framework prioritizes specific and stable candidate markers

To rank the panel objectively, we devised a transparent composite score that integrated the five experimental sub-scores with five annotation-based stability pillars reflecting the burden of conditional regulation and the degree of structural embedding of each gene product (**Figure S2**). Pairwise correlation confirmed that the components were largely non-redundant: specificity correlated positively with fold-change effect size (r = 0.75) and statistical confidence (r = 0.67) but inversely with fold-change consistency (r =-0.69), supporting the separation of specificity from stability (**Figure 5A**). Panel genes were positioned within a composite stability-versus-specificity space and stratified by stability tier (**Figure 5B**), and a final priority score - weighting stability, specificity and predicted subcellular accessibility - ordered the fifty highest-ranked candidates, each annotated by subcellular compartment and functional class (**Figure 5C**), with the per-pillar scores underlying the ranking shown for these fifty candidates in **Figure S3**. *CSRP1*, the gene with the lowest inter-animal variance in the panel, ranked first, followed by *PTGR1*, *TAGLN*, *CNN1* and *MYL4*. The ranking spanned both subcellular compartments: abundant intracellular and cytoskeletal candidates (*CSRP1*, *TAGLN*, *CNN1*, *MYL4*) were accompanied by cell-surface and membrane candidates of conduction or neuronal character, including the collagen-receptor kinase *DDR1*, the sodium-channel auxiliary subunit *SCN4B* and the membrane-associated copine *CPNE5*, providing candidate markers matched to distinct downstream applications. A full ranking of the 286 genes, stability tier, priority score and pathway membership can be found in the Data Supplement (**Table S9**).

### Targeted RT-qPCR validates prioritized Purkinje-enriched transcripts

Sixteen prioritized candidates were tested by RT-qPCR (Purkinje fibers versus cardiomyocytes; Welch’s t test on reference-normalized expression). Thirteen were significantly enriched in PFs, with complete separation between groups and fold-changes spanning roughly three-fold to more than two hundred-fold (all P < 0.005): *TAGLN*, *CNN1*, *COL19A1*, *RIMS1*, *CAPN3*, *SCN4B*, *PTGR1*, *TES*, *DKK3*, *ACTA2*, *MYL7*, *CSRP1* and *S100A16* (**Figure 6**); full per-gene sheep RT-qPCR statistics are provided in the Data Supplement (**Table S8**). *THSD4* was not differentially expressed (P = 0.72), whereas *GYS1* and *TUBA1A* were each significantly enriched in the cardiomyocyte rather than the Purkinje direction (P = 0.04 and P = 0.005, respectively) - results that confirm the directionality and specificity of the prioritized set while delineating its boundaries.

**Figure 6.**
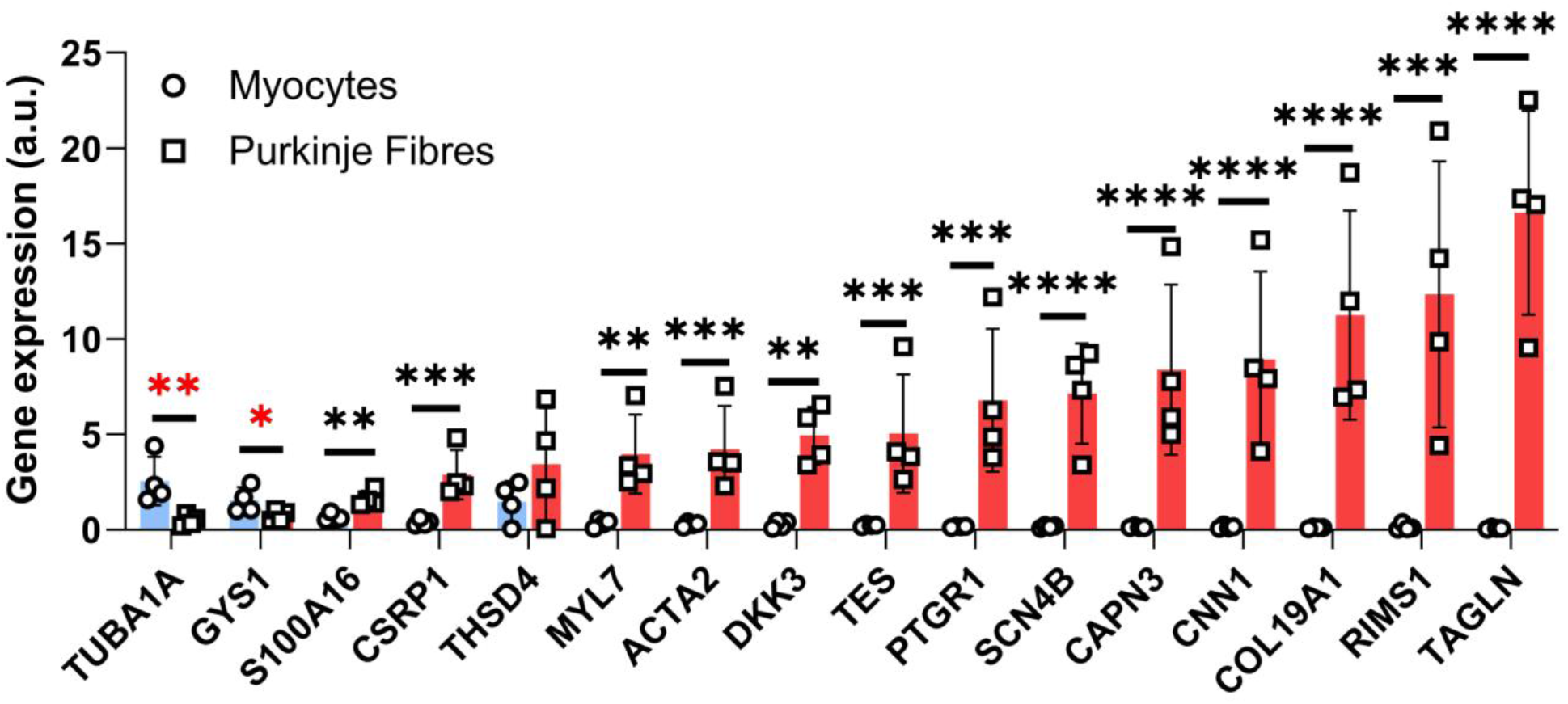
Targeted RT-qPCR validation of prioritized new markers. RT-qPCR of sixteen prioritized candidates in PFs versus cardiomyocytes. Thirteen are significantly enriched in PFs (*TAGLN*, *CNN1*, *COL19A1*, *RIMS1*, *CAPN3*, *SCN4B*, *PTGR1*, *TES*, *DKK3*, *ACTA2*, *MYL7*, *CSRP1* and *S100A16*); *THSD4* is not differentially expressed; and *GYS1* and *TUBA1A* are enriched in the cardiomyocyte rather than the Purkinje direction. Data are mean +/-SD; statistics as in **Figure 2** (n = 4 per group).

### Protein-level validation of lead candidates

We next confirmed expression of representative top candidates at the protein level. Immunohistochemistry localized *MYL4*, *CNN1*, *TAGLN* and *DKK3* to the PFs, with limited staining of the adjacent working myocardium (**Figure 7**), extending validation of these markers from transcript to protein and corroborating the immunofluorescence findings for the conduction-system markers above.

**Figure 7.**
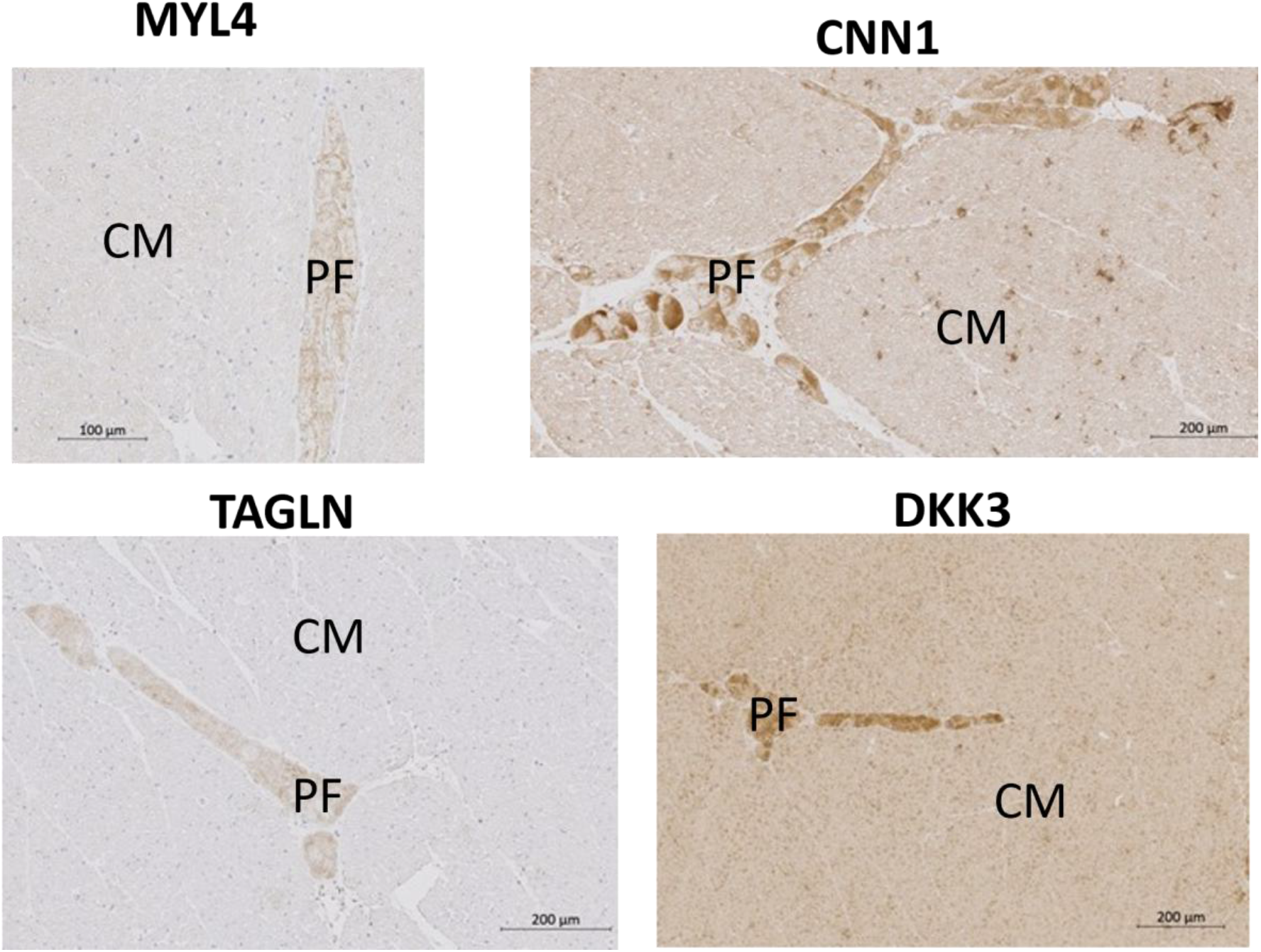
Protein-level validation of lead candidates by immunohistochemistry. Immunohistochemistry (3,3’-diaminobenzidine with a haematoxylin counterstain) for *MYL4*, *CNN1*, *TAGLN* and *DKK3*, each localized to PFs (asterisks) with limited staining of the working myocardium (arrows); scale bars 100 and 200 µm.

### Conservation of candidate markers in the human PF

To assess translational relevance at the transcriptional level, twelve candidates were examined in human ventricular myocardium by RT-qPCR, comparing free-running PFs with left-ventricular cardiomyocytes from four donors (Welch’s t test; **Figure 8**). The canonical conduction markers confirmed the identity and quality of the human Purkinje samples: *MYL4* was strongly enriched in human PFs (approximately eighteen-fold; P < 0.001) and connexin-40 (*GJA5*) was enriched approximately twenty-five-fold (P = 0.04), concordant with the ovine data and with human single-cell atlases of the conduction system ^26^. Among the prioritized markers, a defined subset translated to the human PF: the novel marker *CNTN5* was significantly enriched (approximately fifteen-fold; P = 0.02) and *TAGLN* was significantly enriched (approximately four-fold; P = 0.04), with *CNTN2* showing a concordant Purkinje-direction trend (P = 0.07). By contrast, *CNN1*, *PTGR1*, *SCN4B*, *CNTN4*, *RIMS1* and *TES* were not enriched in human PFs, and *CAPN3* was instead enriched in cardiomyocytes (P = 0.01), reversing its ovine direction. Thus a defined subset of the ovine Purkinje signature - led by *MYL4*, connexin-40, *CNTN5* and *TAGLN* - is conserved at the transcriptional level in the human conduction system, with *CNTN5* emerging as a transcript-and protein-validated Purkinje marker common to both species, while several other markers proved species-restricted.

**Figure 8.**
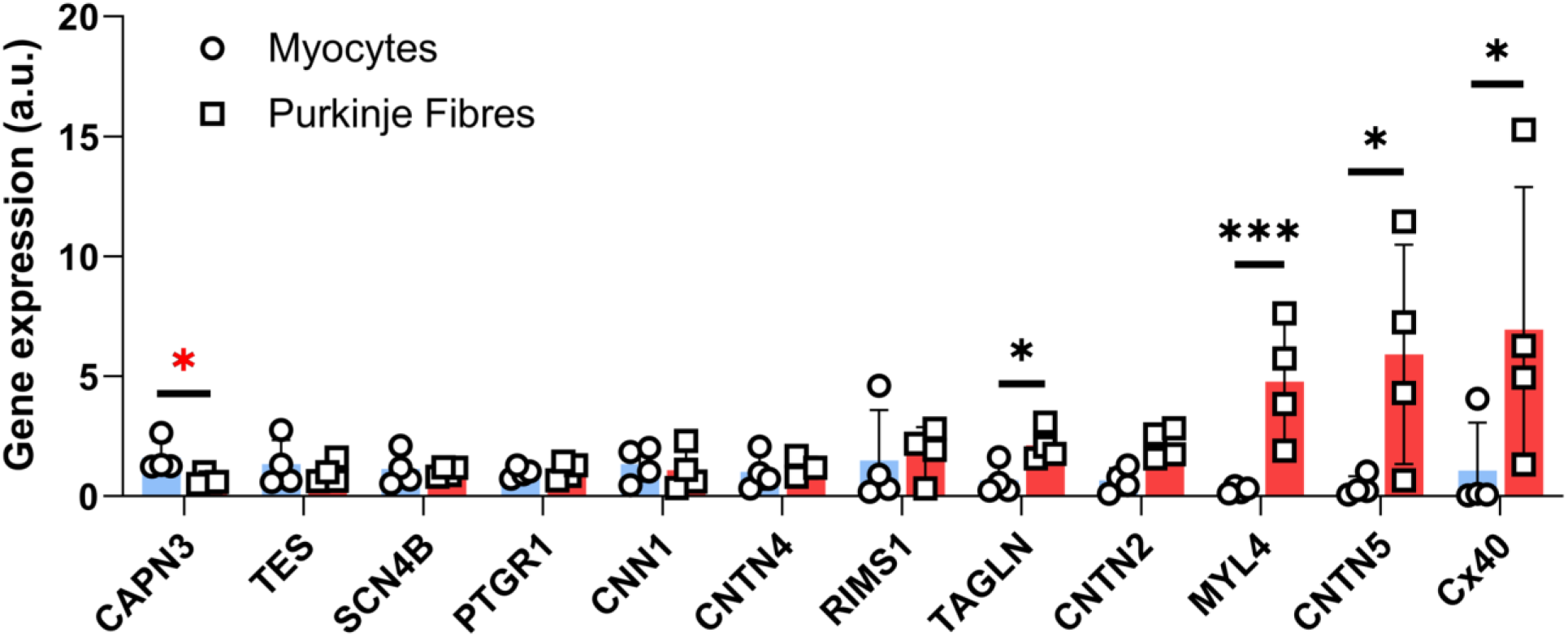
Conservation of candidate markers in the human PF. RT-qPCR of twelve markers in human free-running PFs versus left-ventricular cardiomyocytes from four donors. *MYL4* and connexin-40 (*GJA5*) confirm conserved canonical identity; among the prioritized markers, *CNTN5* and *TAGLN* are significantly enriched in human PFs and *CNTN2* shows a Purkinje-direction trend, and *CAPN3* is enriched in cardiomyocytes. Data are mean +/-SD; statistics as in **Figure 2** (n = 4 donors per group). Full per-gene statistics are provided in **Table S8**.

## Discussion

Using histology-guided LCM and cell-type-resolved transcriptomics, we resolved the molecular identity of ovine PFs against their two most relevant neighbors - working cardiomyocytes and the PF sheath-residing stroma - and prioritized candidate markers with a framework that weights expression specificity together with cross-individual stability. From a PF transcriptome distinct from both comparators, 331 dual-enriched genes were defined, ranked, and validated at the transcript and protein levels, with a defined subset conserved in human myocardium. Beyond cataloguing a Purkinje signature, the study advances the field on three fronts: the first genome-wide, cell-type-resolved portrait of the PF in a large-animal model; an explicit, multi-source paradigm that prioritizes markers likely to remain faithful across physiological and pathological states; and, by resolving candidates by subcellular compartment, a direct link from marker discovery to Purkinje-targeted diagnostic and therapeutic opportunities. These results meet the study objectives and support the central hypothesis, with the single informative exception of *CNTN2*, considered below.

The conceptual impetus for this work is the recognition that a useful tissue marker must be not only specific but faithful: its identification of the target cell must hold across the physiological and pathological states in which it will be applied. The cardiac conduction system is remodeled by ageing, ischemia, heart failure and fibrosis, and many otherwise attractive markers are precisely the genes that respond to these stresses; a marker whose expression rises or falls with disease is, by construction, an unreliable label for the tissue it is meant to denote. We therefore made stability a first-class criterion alongside specificity, defining it as the combination of low inter-individual variability (**Figure 4E**) and a low predicted burden of conditional regulation, and we deliberately excluded diseased and aged comparisons from the marker-defining analysis so that the signature would reflect the resting molecular identity of the PF rather than a state-dependent response. This dual requirement - specificity together with stability - distinguishes the present strategy from conventional fold-change-based marker nomination and is, we propose, the appropriate standard for markers intended for use in the ageing and diseased heart.

To operationalize stability without recourse to longitudinal or multi-condition sampling, the composite score combined five direct experimental sub-scores with five annotation-derived stability pillars that estimate a gene product’s latent propensity to fluctuate (**Figure S2**). The pillars were chosen to capture orthogonal, mechanistically grounded determinants of expression lability. Pillar A (upstream-signal convergence) penalizes genes whose transcription is driven by many converging signaling inputs - adrenergic, calcium, growth-factor, Wnt/Notch, Rho-GTPase, mitogen-activated protein kinase, phosphoinositide 3-kinase and renin-angiotensin among them - because such genes are readily modulated by the same pathways that are activated in disease. Pillar B (transcriptional fragility) penalizes genes under heavy conditional control by transcriptional repression, ubiquitin-proteasomal turnover, post-translational modification, stress-response, autophagy and microRNA regulation, which renders their abundance context-dependent. Pillar C (proteostatic vulnerability) penalizes proteins whose abundance or activity is highly state-sensitive - receptor-and ligand-dependent, conditionally assembled, secreted, kinase-regulated or trafficking-dependent products. Pillar D (multi-pathway breadth and disease responsiveness) penalizes genes embedded in many metabolic, immune, developmental and disease pathways, on the principle that broad pathway participation increases the probability of perturbation. Pillar E (structural embedding), conversely, rewards proteins physically integrated into stable cellular architecture - the cytoskeleton, sarcomere, focal adhesions, conduction hardware such as channels and pumps, lineage-locking transcription factors and the extracellular matrix - because such proteins are buffered against short-term fluctuation and turn over slowly. Pillars A to D therefore act as liability axes and are inverted, whereas pillar E acts as a stabilizing axis and is applied directly; together they encode the explicit hypothesis that a faithful marker is one whose product is structurally anchored and subject to few conditional inputs.

The integrity of the scheme rests on three features. First, each pillar draws on a distinct combination of curated resources - KEGG, Reactome, WikiPathways, the Gene Ontology, UniProt and the matrisome - so that no single annotation source dominates, and pairwise correlation confirmed that the resulting components are largely non-redundant (**Figure 5A**). Second, the framework is transparent, fully auditable and weight-tunable, and integrates direct experimental evidence with meta-analytic annotation within one reproducible pipeline; in this respect it constitutes a state-of-the-art, multi-source approach to marker prioritization rather than reliance on any single statistic. Third, and most importantly, its top-ranked outputs were prospectively validated at the transcript and protein levels (**Figures 6** and **7**), supplying an empirical test of the prioritization itself; the emergence of *CSRP1*, the lowest-variance gene in the panel, as the leading candidate (**Figure 5C**) exemplifies the framework’s intended bias toward robustness as well as specificity.

The signature aligns with established conduction biology while extending it. Canonical fast-conduction, pacemaker and neuronal markers - connexin-40 (*GJA5*), *HCN4*, *NEFM* and *MYL4* - were recapitulated at the transcript and, for connexin-40 and *MYL4*, protein levels (**Figure 2**) ^13,27^. *MYL4* was the most enriched canonical marker (**Figure 2B**) and has been independently nominated as a Purkinje-cell marker in the human cardiac-niche atlas ^26^, while *CNTN5*, which we validated in ovine PFs at both the transcript and protein levels (**Figure 3**), corresponds to the human atrioventricular-bundle and His signature ^26^, consistent with our sampling of proximal conduction tissue; its conservation in human is considered below. Against this concordant background, the failure of *CNTN2* - the most enriched Purkinje transcript in mouse and dog ^13^ - to mark adult ovine ventricular PFs (**Figure 2B**) most plausibly reflects species or developmental-stage differences, or regional specialization of the mature free-running network, and cautions against the uncritical transfer of single rodent markers across species.

Extending the signature across species, human RT-qPCR established that a defined subset of the ovine markers is conserved in the human PF at the transcriptional level (**Figure 8**). *MYL4* and connexin-40 were strongly enriched in human PFs, simultaneously confirming the conservation of the canonical conduction program and the identity of the human samples and matching the human cardiac-niche atlas^26^. Most importantly, the novel marker *CNTN5* - already validated at the transcript and protein levels in sheep - was also significantly enriched in human PFs, establishing it as a transcript-and protein-corroborated cross-species Purkinje marker and the strongest of the newly identified candidates; together with *TAGLN*, these define a compact conserved panel (*MYL4*, connexin-40, *CNTN5* and *TAGLN*) suited to human-relevant identification of the PF. Not every marker translated, however: *CNN1*, *PTGR1*, *SCN4B*, *RIMS1* and the contactin *CNTN4* were not enriched in human PFs, and *CAPN3* reversed direction to the cardiomyocyte compartment; concordant with part of any single-species signature being species-restricted and that empirical cross-species validation is indispensable before clinical extrapolation. This is further exemplified by *CNTN2*, the most enriched Purkinje transcript in rodent and a weak Purkinje-direction trend in human, yet not enriched in adult ovine PFs: a divergence that runs in both directions across species and underscores why a validated, multi-marker panel, rather than any single transferred marker, is the appropriate basis for recognizing the PF across mammals.

The prioritized markers are notable not only for their specificity and stability but for their established cardiovascular phenotypes, which corroborate their relevance to conduction biology and inform their interpretation. *MYL4* encodes the atrial and conduction-system essential myosin light chain, the loss of which causes heritable atrial cardiomyopathy with early-onset atrial fibrillation and atrial standstill ^28^, and its near-absence from working ventricular myocytes makes it an almost binary ventricular Purkinje discriminator. *SCN4B* encodes the sodium-channel auxiliary subunit Na-v-beta-4, mutations of which cause long-QT syndrome type 10 and familial atrial fibrillation and ventricular tachycardia ^29^, and whose resurgent sodium current is mechanistically suited to the prolonged Purkinje action potential. Connexin-40 (*GJA5*) and *HCN4*, recapitulated here, are themselves disease genes, for familial atrial fibrillation ^30^ and for sinus-node dysfunction ^31^ respectively. Among the cytoskeletal candidates, *CSRP1* is associated with dilated cardiomyopathy and its sarcomeric paralogue *CSRP3* (encoding muscle LIM protein) is an established cardiomyopathy gene, while *ACTA2* mutations cause thoracic aortic aneurysm and dissection^32^ - associations that underscore both the contractile-lineage character of these genes and the need to interpret them with spatial context. *DKK3*, a secreted Wnt modulator, is a recognized cardioprotective factor and a cardiac-enriched heart-failure biomarker ^33,34^; its strong Purkinje enrichment is biologically attractive, but because myocardial *DKK3* changes markedly with remodeling it also exemplifies the stability caveat that motivated our framework and is best interpreted in non-diseased tissue. The breadth of these associations - spanning channelopathy, atrial and dilated cardiomyopathy and aortopathy - indicates that the Purkinje molecular signature is populated by genes of genuine clinical consequence and positions the validated markers as entry points to conduction-system disease as well as identity.

A distinctive output of the compartment-aware scoring is a set of markers stratified by subcellular localization, which carries direct translational consequences (**Figure 5C**). Abundant intracellular and cytoskeletal markers - *MYL4*, *MYL7*, *TAGLN*, *CNN1* and *CSRP1* - are optimal for the histological and immunolabelling identification of the conduction system, where signal strength and reproducibility matter most, and indeed several were localized to PFs by immunohistochemistry (**Figure 7**). Their prominence is biologically coherent, since Purkinje cells are classically distinguished by sparse, disorganized myofibrils and an expanded cytoskeletal and glycogen compartment, and TAGLN (SM22-alpha) is recognized to be expressed in myogenic lineages of the developing heart ^35^; because these proteins are also expressed by vascular smooth muscle, however, they are best applied in combination and with spatial confirmation. By contrast, the cell-surface and membrane candidates - connexin-40, the sodium-channel subunit Na-v-beta-4 (*SCN4B*), the contactins and the membrane-associated copine CPNE5 - present opportunities that intracellular markers cannot: surface accessibility enables antibody-and ligand-based detection, flow-cytometric isolation of viable Purkinje cells, and, in principle, in-vivo molecular imaging or cell-type-selective delivery for the mapping or targeted modification of the Purkinje network, an emerging objective in the management of Purkinje-related ventricular arrhythmia^2^. Secreted candidates such as *DKK3* extend this further toward interstitial and circulating biomarker applications. Framing marker discovery in terms of protein compartment thus converts a descriptive catalogue into a menu of application-matched tools, and represents, in our view, an important and generalizable advance for the field.

### Limitations

Several considerations bound the interpretation of these data. First, the stromal comparator comprised the peri-Purkinje collagen sheath rather than purified vascular smooth muscle; this is, however, the contaminant most relevant to in-situ Purkinje microdissection and is directly controlled by the dual-comparison design, and the contractile markers shared with smooth muscle are accordingly recommended for use in combination and with spatial confirmation (**Figure 7**). Second, Purkinje and stromal RNA were necessarily obtained at low input from a cell-sparse starting material; this was addressed by integrity-guided, low-input library chemistry and unique molecular identifier-based quantification, and the close concordance between the sequencing data and independent transcript-and protein-level validation (**Figures 6** and **7**) indicates that the core findings are robust. Third, the RT-qPCR validation used focused biological cohorts (sheep, n = 4 to 5; human, n = 4); the consistency of large, reproducible fold-changes with complete group separation across many independent markers and two protein modalities reinforces confidence, and larger cohorts and digital PCR will further refine effect-size estimates. Fourth, the prioritization weights are deliberately transparent and hypothesis-driven rather than empirically trained, and their prospective validation (**Figures 6** and **7**) provides the relevant performance test; the framework remains fully tunable for sensitivity analysis. Finally, the subcellular-compartment assignments are computational, so the surface and secreted candidates proposed for targeting applications warrant explicit subcellular confirmation; functional and electrophysiological characterization of the candidate markers, and their extension to disease settings, are natural next steps that lie beyond the descriptive scope of the present study.

## Conclusions

In summary, we provide a cell-type-resolved molecular portrait of the ovine PF and a transparent, multi-source framework that prioritizes candidate markers on the dual criteria of specificity and cross-individual stability - the latter chosen so that the resulting markers are likely to remain faithful in the ageing and diseased heart. The approach recapitulates and extends the canonical conduction signature, nominates protein-validated and disease-relevant candidates, demonstrates through the divergence of *CNTN2* the necessity of cross-species validation, and, by resolving markers according to subcellular compartment, defines application-matched tools ranging from robust intracellular labels for conduction-system histology to surface targets for the isolation, imaging and selective modification of Purkinje cells. Human RT-qPCR confirms that a defined subset of the signature-including *MYL4*, connexin-40 and the novel marker *CNTN5* - is conserved at the transcriptional level in the human conduction system. By bridging rodent mechanistic studies and human single-cell atlases in a translationally relevant large-animal model, this work establishes both a validated, prioritized Purkinje-marker resource and a generalizable, state-of-the-art strategy for the discovery of stable, faithful cell-type markers.

## Acknowledgements

Laser-capture microdissection and RNA-sequencing were performed at the PUMA core facility (Neurocentre Magendie, INSERM U1215, University of Bordeaux).

## Nonstandard Abbreviations and Acronyms

cDNA: Complementary DNA
cTnT: cardiac troponin T
GEO: Gene Expression Omnibus
LCM: laser-capture microdissection
PEN: polyethylene naphthalate
PF: Purkinje fiber
RIN: RNA integrity number
RT-qPCR: reverse transcription quantitative polymerase chain reaction
UMI: unique molecular identifier

## Novelty and Significance

### What is known?

- The His-Purkinje network is central to rapid ventricular activation and is a key substrate for life-threatening ventricular arrhythmias and a target for catheter ablation.
- The molecular definition of the cardiac conduction system derives largely from rodent models, and few markers are validated at the protein level or shown to transfer to large mammals or human tissue.
- Established Purkinje markers are limited by sharing with other tissues, inconsistent cross-species conservation, or intracellular localization that precludes surface-based targeting.

### What new information does this article contribute?

- Using RNA-sequencing, this study provides the first genome-wide, cell-type-resolved transcriptome of the cardiac Purkinje fiber in a large-animal (ovine) model, resolved against cardiomyocytes and stroma.
- It introduces a transparent, multi-source framework that prioritizes markers by specificity, cross-individual stability and predicted subcellular accessibility, with leading candidates validated at the transcript level by RT-qPCR and at the protein level by immunolabelling.
- It identifies contactin-5 as a novel, protein-validated Purkinje marker conserved in human myocardium and shows that the rodent gold-standard contactin-2 is not enriched in ovine Purkinje fibers, demonstrating the necessity of cross-species validation.

## Summary paragraph

The cardiac Purkinje network governs rapid, synchronous ventricular activation and is a principal source of malignant ventricular arrhythmias, yet it remains one of the least molecularly defined regions of the heart, and the markers used to recognize it derive largely from rodents and transfer unreliably to large mammals and humans. Using histology-guided laser-capture microdissection and cell-type-resolved RNA-sequencing in the sheep - a translational model with human-like Purkinje anatomy and electrophysiology - we resolve the Purkinje transcriptome against its immediate neighbors and prioritize candidate markers not by enrichment alone but by specificity, cross-individual stability and predicted subcellular accessibility, the properties that determine real-world utility. Leading candidates are validated at transcript and protein levels and tested for conservation in human myocardium, yielding an application-matched toolkit that spans robust intracellular labels and surface-accessible targets and identifying contactin-5 as a novel cross-species Purkinje marker. By showing that the canonical rodent marker contactin-2 is not conserved in the sheep, the work underscores the need for cross-species validation and provides both a validated marker resource and a generalizable strategy for discovering stable, faithful cell-type markers.

