## Supplemental materials for "Cell-Type-Resolved Transcriptomics Defines Stable and Accessible Markers of the Cardiac Purkinje Fiber in Sheep and Human Translation"

##### RT-qPCR validation

RNA for RT-qPCR was extracted from Purkinje fibers using RLT buffer with proteinase K and the QIAGEN miRNeasy Micro kit, and from left-ventricular myocardium using QIAzol and the miRNeasy kit; quantity and purity were assessed by spectrophotometry (NanoDrop). Primers were designed from Ensembl sequences with the NCBI Primer designing tool, synthesised by Sigma-Aldrich and efficiency-validated, with the full list provided in the Data Supplement (**Table S2**). Complementary DNA (cDNA) was synthesised from 200 ng of RNA (OZYME) and amplified with SYBR chemistry (OZYME) on a Bio-Rad CFX96 instrument (95°C for 3 min; 40 cycles of 95°C for 5 s and 60°C for 30 s; followed by melt-curve analysis) in technical triplicate. Relative quantification used the 2^(-delta-delta-Ct) method with *HPRT1* and *B2M* as reference genes in sheep and B2M and GUSB in human. Purkinje fibers were compared with cardiomyocytes (n = 4 biological replicates per group) as described under Statistical analysis. For *CNTN5*, amplification in several ovine cardiomyocyte samples fell at or below the reliably measurable cycle window, so cardiomyocyte expression for that gene is reported as at or below the limit of detection.

##### Free-running PF histology

For histopathological characterization of the free-running PF, false tendons were dissected and formaldehyde-fixed. PFA was rinsed in a phosphate buffer saline (PBS) solution under agitation at 4 °C, then samples were dehydrated and embedded in paraffin using a HistoCore Pearl automat (Leica Biosystems) with successive baths of ethanol at increasing concentration (95% three times, 100% three times), a toluene rinse (three times), and paraffin baths (twice); paraffin blocks were stored at -20°C until use. Embedded tissue samples were sectioned at 6 µm on a RM2255 microtome (Leica) were mounted on to glass slides. Sections were treated with toluene to remove paraffin then rehydrated by serial dilutions of ethanol at 100%, 95%, 80%, followed by immersion in tap water.
Sections randomly assigned to Masson's trichrome staining were treated with: xylidine ponceau, 5 minutes; rinsing with distilled water, 5 minutes; Biebrich Scarlet-acid fuchsin solution, 1 minute; 3% phosphomolybdic, 1 minute; 1% glacial acetic acid, 1 minute; 1% light green SF yellowish, 2 minutes and then 1% acetic acid, 1 minute. Sections were again dehydrated by ethanol 95%, 6 minutes and 100% ethanol, 6 minutes followed by toluene, 3 minutes and stored at room temperature in air.
Sections randomly assigned to periodic acid–Schiff staining were immersed in: 0.5% periodic acid, 5 minutes; rinsing in distilled water, two changes; Schiff's reagent, 15 minutes, protected from light; running tap water, 10 minutes, to develop the reaction; Mayer's haematoxylin, 3 minutes; and running tap water, 5 minutes, to blue the nuclei.
Slides were subsequently scanned in brightfield using an automated slide scanner, Axio Scan Z1 (Zeiss), and observed with Zen Lite Software (Zeiss).

##### Laser-capture microdissection

Laser-capture microdissection (LCM) and RNA-sequencing were performed at the PUMA core facility (Neurocentre Magendie, INSERM U1215, University of Bordeaux). Purkinje cells, cardiomyocytes and collagen sheath-residing stromal cells were targeted specifically and extracted by LCM for downstream RNA-sequencing, guided by cresyl-violet histological staining of PF-rich transeptal sections of the ventricles. Samples were embedded in OCT compound, snap-frozen in liquid nitrogen and stored at -80°C until processing. Cryosections were cut at 20 µm (Purkinje fibers, stroma) or 30 µm (cardiomyocytes) on a Leica CM3050S cryostat and mounted onto RNase-free 1-mm glass slides coated with a polyethylene naphthalate (PEN) membrane (P.A.L.M. Microlaser Technologies AG, Bernried, Germany). To remove the OCT compound, sections were fixed in 95% ethanol for 30 s, then 75% ethanol for 30 s and 50% ethanol for 30 s. Sections were then stained with 1% cresyl violet prepared in 50% ethanol for 30 s and dehydrated sequentially in 50%, 75% and 95% ethanol for 30 s each, followed by two washes in 100% ethanol for 30 s each. Laser microdissection was performed on a Zeiss P.A.L.M. MicroBeam system (355 nm ultraviolet laser coupled to an inverted microscope equipped with PALM software; Carl Zeiss). Target cells were isolated by laser cutting followed by laser pressure catapulting into 0.5-mL adhesive cap tubes (Carl Zeiss Microscopy, Jena, Germany). Three populations were captured from each animal - Purkinje fibers, left-ventricular cardiomyocytes and stroma derived from the peri-Purkinje collagen sheath. Approximately 5-8 mm² of Purkinje fibers and peri-Purkinje collagen sheath were captured per animal, and approximately 30 mm² for cardiomyocytes.

**RNA isolation, library preparation and sequencing**

Cells were lysed immediately after microdissection in 250 µL of guanidine isothiocyanate-containing BL buffer (ReliaPrep RNA Cell Miniprep System, Promega, Madison, WI, USA) supplemented with 10 µL of 1-thioglycerol and stored at -80°C until RNA extraction. Total RNA was extracted using the ReliaPrep RNA Cell Miniprep System (Promega), including on-column DNase treatment, and eluted in 15 µL following the manufacturer's instructions. RNA quality was assessed on an Agilent 2100 Bioanalyzer (Agilent Technologies, Santa Clara, CA, USA) with the Eukaryote Total RNA Pico assay (Agilent, 5067-1513). RNA integrity numbers (RIN) were 7.9-8.2 for cardiomyocytes and 5.3-7.0 for Purkinje fibers. For stroma, the RNA quantity was at the limit of detection, so a reliable RIN could not be determined and was not used; because the Purkinje and stromal material was obtained from the same sections, the stromal (collagen-derived) RNA was taken to be of quality equivalent to that of the Purkinje fibers and of sufficient quality to be processed together with the cardiomyocyte RNA using a poly(A) capture strategy.

Sequencing libraries were prepared with the Takara SMART-Seq mRNA LP (with UMIs) kit (Takara Bio, 634762; protocol version 100422), a low-input-tolerant chemistry that generates full-length, template-switching cDNA by oligo(dT) priming and incorporates a unique molecular identifier (UMI) into the template-switching oligonucleotide for 5-prime UMI counting. Input RNA was 0.5 ng for Purkinje fibers and stroma and 10 ng for cardiomyocytes; cDNA was amplified by 12 cycles (Purkinje fibers, stroma) or 8 cycles (cardiomyocytes) of long-distance PCR, and libraries were dual-indexed by 16 or 14 cycles of indexing PCR with a Unique Dual Index kit (8-base-pair i7 and i5 indices; Takara Bio, 634753; Table S4). cDNA and final libraries were quality-controlled on a LabChip GX Touch HT Nucleic Acid Analyzer (Revvity) with the HT DNA NGS 3K Reagent Kit (Revvity, CLS960013) on an HT DNA X-Mark Chip (Revvity, CLS144006), and were then quantified by qPCR on a LightCycler 480 II (Roche) with the NEBNext Library Quant Kit for Illumina (NEB, E7630L). Libraries were pooled in equivalent amounts for Purkinje fibers and stroma and at twice that depth for cardiomyocytes, and sequenced on a NextSeq 2000 System (Illumina, 20038897) with a NextSeq 1000/2000 P2 XLEAP-SBS Reagent Kit (300 cycles; Illumina, 20100985) and a PhiX spike-in. Paired-end sequencing (2 x 150 base pairs) yielded a mean of approximately 24.5 million read pairs per sample (range 15.2-39.2 million across libraries).

**Read processing, alignment, quality control and differential expression**

Sequencing data were processed with the Cogent NGS Analysis Pipeline v2.0.1 (Takara Bio), which comprises a demultiplexer that extracted the sample barcodes and generated paired Read 1 and Read 2 FASTQ files, and an analyzer that performed read trimming (Cutadapt v3.4), genome indexing and alignment (STAR v2.7.8a) [1], and UMI parsing and gene-level counting (SAMtools v1.9, bedtools v2.29.2 and Subread v2.0.0) against the *Ovis aries* reference genome (assembly ARS-UI_Ramb_v2.0; Ensembl release 113; **Table S5**). Mean mapping rates were 92.3% for cardiomyocytes, 90.2% for Purkinje fibers and 81.9% for stroma. Reads sharing a unique molecular identifier were collapsed to per-gene molecule counts, and the resulting UMI-based gene-matrix count file was imported into RStudio for differential-expression analysis. Quality control - comprising the variance-stabilised mean-standard-deviation relationship, gene-detection saturation, library complexity and multidimensional scaling - was performed with RNAseqQC v0.2.1 in RStudio, and genes with zero counts across all samples were removed. Differential expression was assessed with DESeq2 using the Wald test, with control of the false-discovery rate. Two contrasts were carried forward - Purkinje fibers versus cardiomyocytes and Purkinje fibers versus stroma - and genes were considered Purkinje-enriched at an absolute log2 fold-change greater than 1.0 and an adjusted P value below 0.01.

**Functional annotation and prioritization framework**

Genes were annotated for pathway and ontology membership from KEGG [2], Reactome [3], WikiPathways [4], and Gene Ontology [5], through g:Profiler [6], data version e113_eg59_p19 (Ensembl release 113), and MyGene.info [7], accessed December 2025; UniProt subcellular localization was obtained from UniProtKB release 2025_04 [8], and core matrisome membership from the NABA gene sets (MSigDB v2025.1.Hs) [9,10].

Candidates were prioritized by a transparent, single-level composite score that combined five experimental sub-scores with five annotation-based stability pillars, each expressed as a percentile rank or a normalized value across the 286 scored genes. The experimental sub-scores were fold-change effect size, fold-change consistency across the two contrasts, statistical confidence (the geometric mean of the negative log10 adjusted P values), expression abundance, and inter-animal consistency (one minus the coefficient of variation). The five stability pillars quantified, from binary annotation-term membership, the latent propensity of each gene product to fluctuate: (A) upstream-signal convergence, (B) transcriptional fragility, (C) proteostatic vulnerability, (D) multi-pathway breadth and disease responsiveness, and (E) structural embedding; pillars A to D were treated as instability (liability) axes and inverted, whereas pillar E was applied directly. The 41-term pillar dictionary, with database identifiers and scoring direction, is provided in the Data Supplement (Table S7). A composite stability score (with a default weighting of experimental to annotation components of 2:1) defined stability tiers (high at or above the 0.67 percentile; lower at or below 0.33; borderline otherwise). A specificity score was computed as the percentile rank of log1p of the product of mean Purkinje expression and mean log2 fold-change, and a compartment bonus reflected predicted accessibility (cell-surface 1.0; intracellular 0.7; unknown 0.5) derived from Gene Ontology cellular-component and UniProt localization. The final priority score was 0.55 times the stability percentile plus 0.35 times specificity plus 0.10 times the compartment bonus, and ordered all 286 scored genes (Figure 5C). The framework is implemented using an in-house Python-based analytical suite.

**Table S1. Human cardiac tissue donors**

| Donor | Sex | Age (y) | Infarction status / sampling |
|---|---|---|---|
| hFP1 | Female | 65 | No infarction |
| hFP2 | Male | 55 | Prior myocardial infarction; PF sampled remote from infarct |
| hFP3 | Male | 74 | Prior myocardial infarction; PF sampled remote from infarct |
| hFP4 | Female | 26 | No infarction |

*Table S1. Human heart donors (N = 4) used for cross-species RT-qPCR validation. Hearts were obtained from brain-dead donors unsuitable for transplantation through a program approved by the French Biomedicine Agency (PFS14-004), in collaboration with the University Hospital of Bordeaux, with family consent and in conformity with the Declaration of Helsinki.*

**Table S2. RT-qPCR primer sequences**

| Target gene | Species | Forward primer (5'->3') | Reverse primer (5'->3') |
|---|---|---|---|
| *HPRT1* | Sheep | TGCTGAGGATTTGGAGAAGGTG | CCAACAGGTCGGCAAAGAAC |
| *B2M* | Sheep | CTGTCGCTGTCTGGACTGG | CCCCGTTCTTCAGCAAATCG |
| *ACTA2* | Sheep | TAGCCCTGGACTTTGAGAACG | GAAACGCTCATTCCCGATGG |
| *CAPN3* | Sheep | TCAACATGCGCGAGGTGTC | AAGACCCGGAGGATGAACTC |
| *CNN1* | Sheep | GAACACCAACCACACGCAAG | CTCCGCGTATTTCACTCCCA |
| *CNTN2* | Sheep | QIAGEN QuantiTect Primer Assay, cat. 249900 | |
| *CNTN4* | Sheep | ATCAGGATAATCGCCGCTTCG | AATATCAGTGGTGTGGGTGGTC |
| *CNTN5* | Sheep | QIAGEN QuantiTect Primer Assay, cat. 249900 | |
| *COL19A1* | Sheep | GCCGAGGGAGATGTGTTGAA | TCGGGCAGCAATAAGAGTCC |
| *CSRP1* | Sheep | GCTACGGCAAGAAGTACGGG | TCCTCGTGTCTGATGCCGA |
| *DKK3* | Sheep | GTGCTGTGCCTTCCAGAGAG | GTGATGAGGTCCAGAAGCCG |
| *GJA5 (Cx40)* | Sheep | AGTGAATGGAAGGATTGCCCT | GGACACATAGCAGTTGACGGG |
| *GYS1* | Sheep | GAAGGCAGCGCATTATCCAG | TCGTGATGCTCGTAGGTGAAG |
| *HCN4* | Sheep | CAACTCCTGGGGAAAGCAGT | ATGAACATGGCGTAGCAGGT |
| *MYL4* | Sheep | TGGACTTTGAGACCTTCCTGC | CTTGTCGAACACACGCAGC |
| *MYL7* | Sheep | CTTCCGCCTTTTTGACCCCAG | AAACATCTGCTCCACCTCAGC |

| Target gene | Species | Forward primer (5'->3') | Reverse primer (5'->3') |
|---|---|---|---|
| *NEFM* | Sheep | QIAGEN QuantiTect Primer Assay, cat. 249900 | |
| *PTGR1* | Sheep | ACTATTGTGGTGGCTCCTTCTG | AGCCAGAGACAGTGGTAATGTG |
| *RIMS1* | Sheep | CAAGCATGAGCGGAGAGATGT | CACTATGGCAACCACTTTGGC |
| *S100A16* | Sheep | TAGAGAAGGCGGTGGTCGT | GCTGCTCTTGCTGATCTTGT |
| *SCN4B* | Sheep | TGAGAAATCGGACCCCAAGG | TATTTGCCCGTGTCGCTGAA |
| *TAGLN* | Sheep | - | - |
| *TBX5* | Sheep | ACAAAGTGAAGGTGACCGGC | GAGCTGCCGCATCCAATGAG |
| *TES* | Sheep | GATGTCAAACTCCCCCGTGA | TGTGTTCCGCTGATTTGTCCT |
| *THSD4* | Sheep | GGTTATGGGAAAGCCCCGT | TGAGCTGAAGGGTGGTAGTC |
| *TUBA1A* | Sheep | ACCTAAGGATGTCAACGCCG | GCTCAGCATACACACAGCAC |
| *B2M* | Human | GGGCATTCCTGAAGCTGACAGCA | TGTCGGATGGATGAAACCCAGACAC |
| *GUSB* | Human | GGAGAGCTCATTTGGAATTTTGCCG | TGGCTACTGAGTGGGGATACCTGG |
| *CAPN3* | Human | GCCAGGAGCAAAACCTACATC | GGTGGAGGGCACGATGAC |
| *CNN1* | Human | ACTGCGACACGCTCAATGT | GTTGTGTGCGTGGTGGTTG |
| *CNTN2* | Human | Bio-Rad PrimePCR SYBR Green Assay, cat. 10025636 | |
| *CNTN4* | Human | Bio-Rad PrimePCR SYBR Green Assay, cat. 10025636 | |
| *CNTN5* | Human | Bio-Rad PrimePCR SYBR Green Assay, cat. 10025636 | |
| *GJA5 (Cx40)* | Human | AACACAGACAGGCAGAGGATT | AGCCAGACCTTGCCTACCAC |
| *MYL4* | Human | GAGACGTTCTTGCCCATCCT | GTGCCATTGCTCTCCTTGTC |
| *PTGR1* | Human | GGACAACGCACTCCATTTCTG | GTCCCCAGAGCCAAAGACAG |
| *RIMS1* | Human | CAGACGGATGGGGACTTCAG | GCACTAAGGCTGGATCTCCG |
| *SCN4B* | Human | Bio-Rad PrimePCR SYBR Green Assay, cat. 10025636 | |
| *TAGLN* | Human | TGTACCCTGATGGCTCCAAG | ATAGTCCTCAGCCGCCTTCA |
| *TES* | Human | CCGAAAGGGCTGGCTATGAT | CCAGCACATCGGGGTTTCT |

*Table S2. Primers used for RT-qPCR (sheep and human). Sequences were obtained from Ensembl reference sequences, designed with the NCBI Primer designing tool, efficiencies were validated before use. Reference genes: HPRT1 and B2M (sheep); B2M and GUSB (human).*

#### Table S3. Antibodies and detection reagents

| Target / reagent | Host / type | Dilution | Supplier (cat. no.) | Use |
|---|---|---|---|---|
| Cardiac troponin T (cTnT) | Mouse monoclonal | 1:500 | Invitrogen (MA5-12960) | IF |
| GJA5 / Cx40 | Biotin-conjugated | 1:200 | Bioss (bs-1050R-biotin) | IF |
| MYL4 | Biotin-conjugated | 1:300 | MyBioSource (MBS2095280) | IF, IHC |
| CNTN5 | Rabbit polyclonal | 1:500 | Sigma-Aldrich (HPA039492) | IF |
| TAGLN | Biotin-conjugated | 1:500 | Bio-Techne (NBP3-20101B) | IHC |
| CNN1 | Rabbit polyclonal | 1:200 | Proteintech (13938-1-AP) | IHC |
| DKK3 | Rabbit polyclonal | 1:200 | Proteintech (10365-1-AP) | IHC |

**Secondary / detection reagents**

| Target / reagent | Host / type | Dilution | Supplier (cat. no.) | Use |
|---|---|---|---|---|
| Donkey anti-mouse Alexa Fluor 488 | - | 1:1000 | Invitrogen (A21202) | IF |
| Donkey anti-rabbit Alexa Fluor 555 | - | 1:1000 | Invitrogen (A31572) | IF |
| Streptavidin Alexa Fluor 647 | - | 1:1000 | Invitrogen (S21374) | IF |
| Goat anti-rabbit IgG (H+L), biotinylated | - | 1:500 | Vector Laboratories (BA-1000-1.5) | IHC |

*Table S3. Primary antibodies, secondary antibodies and detection reagents for immunofluorescence (IF) and immunohistochemistry (IHC). Biotin-conjugated primaries were revealed with streptavidin-Alexa Fluor 647 (IF) or an avidin-biotin-peroxidase complex (IHC). IF imaging used a Leica THUNDER Imager with a Leica DFC9000 GTC camera.*

#### Table S4. Sequencing library dual indices

| Sample | Population | Index (UDI) | i7 (8 bp) | i5 (8 bp) |
|---|---|---|---|---|
| PF130923 | Purkinje fiber | U177 | ATTATCGA | CATCCTAA |
| PF190923 | Purkinje fiber | U179 | CGCAGAAG | GTAATCGT |
| PF241023 | Purkinje fiber | U181 | ATCAGAGA | GGCTGCTA |
| PF301023 | Purkinje fiber | U183 | GTCTCGCG | AGGTATGG |
| PF220124 | Purkinje fiber | U185 | CAACCTCT | GATGGCCA |
| PF240124 | Purkinje fiber | U187 | CTTCGTTA | TACTGGCC |
| Stroma130923 | Stroma | U178 | GTCGCTAG | TGCTTGAT |
| Stroma190923 | Stroma | U180 | TCAGGACC | AATGGTAA |
| Stroma241023 | Stroma | U182 | CTGGATAA | CCGGCTAC |
| Stroma301023 | Stroma | U184 | TCGAATAA | GAACTCAG |
| Stroma220124 | Stroma | U186 | AGTTGAAC | TGCGCGTT |
| Stroma240124 | Stroma | U188 | GCTGCGGC | ACTAGCTC |
| CM130923 | Cardiomyocyte | U189 | AGTAACCT | GTACCGTA |
| CM190923 | Cardiomyocyte | U190 | TAAGATAT | AGGTAGTT |
| CM241023 | Cardiomyocyte | U191 | TGGTCCTG | GCCTATTC |
| CM301023 | Cardiomyocyte | U192 | GTCCTAAC | ATGACCGA |
| CM220124 | Cardiomyocyte | U171 | TCTCCGTC | ACTTCGTT |
| CM240124 | Cardiomyocyte | U172 | CCGATCCT | GGCATATA |

*Table S4. Unique dual indices (UDI) assigned to each library (Takara Unique Dual Index Kit, plate 97-192). i7 and i5 are the 8-bp index sequences as read on the Illumina NextSeq 2000.*

#### Table S5. Per-sample RNA-sequencing quality control (18 libraries)

| Sample | Population | RIN | Input (ng) | Barcoded reads (M) | Mapping (%) | Genes |
|---|---|---|---|---|---|---|
| PF#1 | Purkinje fiber | 6 | 0.5 | 15.6 | 88 | 13,260 |
| PF#2 | Purkinje fiber | 5.3 | 0.5 | 15.8 | 88.8 | 13,434 |
| PF#3 | Purkinje fiber | 6.8 | 0.5 | 17.2 | 86.5 | 13,891 |
| PF#4 | Purkinje fiber | 6.7 | 0.5 | 15.2 | 91.7 | 13,826 |
| PF#5 | Purkinje fiber | 6.7 | 0.5 | 15.9 | 92.8 | 15,213 |
| PF#6 | Purkinje fiber | 7 | 0.5 | 18.2 | 92.4 | 15,501 |
| Stroma#1 | Stroma | - | 0.5 | 17.3 | 83.2 | 15,442 |
| Stroma#2 | Stroma | - | 0.5 | 21.2 | 83.7 | 15,689 |
| Stroma#3 | Stroma | - | 0.5 | 21.9 | 76.2 | 15,026 |
| Stroma#4 | Stroma | - | 0.5 | 20.1 | 80.4 | 13,989 |
| Stroma#5 | Stroma | - | 0.5 | 20.8 | 85.1 | 16,954 |
| Stroma#6 | Stroma | - | 0.5 | 27.5 | 80.6 | 17,871 |
| CM#1 | Cardiomyocyte | 8.2 | 10 | 39.2 | 91.6 | 17,797 |
| CM#2 | Cardiomyocyte | 8 | 10 | 37.7 | 92.1 | 17,742 |
| CM#3 | Cardiomyocyte | 8.2 | 10 | 34.1 | 92.7 | 17,024 |
| CM#4 | Cardiomyocyte | 8 | 10 | 32.7 | 92.5 | 17,090 |
| CM#5 | Cardiomyocyte | 8 | 10 | 37.5 | 92 | 17,249 |
| CM#6 | Cardiomyocyte | 7.9 | 10 | 33 | 93.1 | 17,012 |

*Table S5. Per-sample RNA-sequencing metrics for the 18 sequenced libraries (6 Purkinje fiber, 6 cardiomyocyte, 6 stroma). RIN, RNA integrity number; input, RNA mass into library preparation; mapping (%), reads mapped/trimmed reads.*

#### Table S6. Purkinje-fiber-enriched genes without a mapped human ortholog (45 loci; differential expression only)

| Locus | Ensembl ID | Mean log2FC | Min adj. P | Status |
|---|---|---|---|---|
| U01 | ENSOARG00020026390 | 6.08 | 2.59e-115 | No stable gene symbol or human ortholog; excluded from scoring |
| U02 | ENSOARG00020001206 | 3.49 | 2.52e-82 | No stable gene symbol or human ortholog; excluded from scoring |
| U03 | ENSOARG00020036760 | 3.67 | 1.22e-63 | No stable gene symbol or human ortholog; excluded from scoring |
| U04 | ENSOARG00020039990 | 1.90 | 3.77e-59 | No stable gene symbol or human ortholog; excluded from scoring |
| U05 | ENSOARG00020018953 | 5.96 | 2.60e-53 | No stable gene symbol or human ortholog; excluded from scoring |
| U06 | ENSOARG00020026311 | 3.81 | 2.53e-33 | No stable gene symbol or human ortholog; excluded from scoring |
| U07 | ENSOARG00020039894 | 4.10 | 1.43e-32 | No stable gene symbol or human ortholog; excluded from scoring |
| U08 | ENSOARG00020034652 | 1.85 | 4.96e-31 | No stable gene symbol or human ortholog; excluded from scoring |
| U09 | ENSOARG00020000917 | 2.86 | 7.02e-31 | No stable gene symbol or human ortholog; excluded from scoring |
| U10 | ENSOARG00020016172 | 1.44 | 4.89e-28 | No stable gene symbol or human ortholog; excluded from scoring |
| U11 | ENSOARG00020021008 | 3.99 | 3.35e-26 | No stable gene symbol or human ortholog; excluded from scoring |
| U12 | ENSOARG00020032004 | 1.63 | 5.41e-25 | No stable gene symbol or human ortholog; excluded from scoring |
| U13 | ENSOARG00020018455 | 2.34 | 3.78e-24 | No stable gene symbol or human ortholog; excluded from scoring |
| U14 | ENSOARG00020011746 | 6.31 | 5.51e-21 | No stable gene symbol or human ortholog; excluded from scoring |
| U15 | ENSOARG00020027339 | 2.20 | 1.81e-20 | No stable gene symbol or human ortholog; excluded from scoring |
| U16 | ENSOARG00020036459 | 5.32 | 3.90e-20 | No stable gene symbol or human ortholog; excluded from scoring |
| U17 | ENSOARG00020033895 | 1.53 | 9.08e-20 | No stable gene symbol or human ortholog; excluded from scoring |
| U18 | ENSOARG00020014228 | 1.76 | 2.36e-19 | No stable gene symbol or human ortholog; excluded from scoring |
| U19 | ENSOARG00020040379 | 3.16 | 2.63e-19 | No stable gene symbol or human ortholog; excluded from scoring |
| U20 | ENSOARG00020011597 | 2.15 | 8.45e-19 | No stable gene symbol or human ortholog; excluded from scoring |
| U21 | ENSOARG00020010580 | 2.05 | 1.06e-18 | No stable gene symbol or human ortholog; excluded from scoring |
| U22 | ENSOARG00020024082 | 1.36 | 3.26e-18 | No stable gene symbol or human ortholog; excluded from scoring |
| U23 | ENSOARG00020023033 | 2.70 | 5.73e-18 | No stable gene symbol or human ortholog; excluded from scoring |
| U24 | ENSOARG00020036096 | 2.09 | 6.18e-18 | No stable gene symbol or human ortholog; excluded from scoring |
| U25 | ENSOARG00020034316 | 2.89 | 1.28e-17 | No stable gene symbol or human ortholog; excluded from scoring |
| U26 | ENSOARG00020026408 | 2.69 | 3.34e-17 | No stable gene symbol or human ortholog; excluded from scoring |
| U27 | ENSOARG00020013021 | 1.61 | 8.27e-15 | No stable gene symbol or human ortholog; excluded from scoring |
| U28 | ENSOARG00020028418 | 1.15 | 3.31e-14 | No stable gene symbol or human ortholog; excluded from scoring |
| U29 | ENSOARG00020024897 | 1.20 | 3.50e-14 | No stable gene symbol or human ortholog; excluded from scoring |
| U30 | ENSOARG00020010016 | 1.67 | 5.62e-13 | No stable gene symbol or human ortholog; excluded from scoring |
| U31 | ENSOARG00020031342 | 2.01 | 2.19e-12 | No stable gene symbol or human ortholog; excluded from scoring |
| U32 | ENSOARG00020007097 | 1.37 | 2.23e-12 | No stable gene symbol or human ortholog; excluded from scoring |
| U33 | ENSOARG00020034858 | 1.13 | 9.98e-12 | No stable gene symbol or human ortholog; excluded from scoring |
| U34 | ENSOARG00020006224 | 1.28 | 1.66e-11 | No stable gene symbol or human ortholog; excluded from scoring |
| U35 | ENSOARG00020031009 | 1.67 | 3.38e-11 | No stable gene symbol or human ortholog; excluded from scoring |
| U36 | ENSOARG00020033126 | 1.83 | 1.45e-10 | No stable gene symbol or human ortholog; excluded from scoring |
| U37 | ENSOARG00020038425 | 1.87 | 2.00e-10 | No stable gene symbol or human ortholog; excluded from scoring |
| U38 | ENSOARG00020034154 | 2.03 | 7.86e-10 | No stable gene symbol or human ortholog; excluded from scoring |
| U39 | ENSOARG00020036341 | 1.59 | 2.44e-09 | No stable gene symbol or human ortholog; excluded from scoring |
| U40 | ENSOARG00020002769 | 1.60 | 2.49e-09 | No stable gene symbol or human ortholog; excluded from scoring |
| U41 | ENSOARG00020009504 | 1.83 | 3.07e-08 | No stable gene symbol or human ortholog; excluded from scoring |
| U42 | ENSOARG00020028366 | 2.44 | 1.50e-06 | No stable gene symbol or human ortholog; excluded from scoring |
| U43 | ENSOARG00020026541 | 1.11 | 2.16e-06 | No stable gene symbol or human ortholog; excluded from scoring |
| U44 | ENSOARG00020035313 | 1.55 | 7.40e-05 | No stable gene symbol or human ortholog; excluded from scoring |
| U45 | ENSOARG00020020457 | 1.96 | 9.23e-04 | No stable gene symbol or human ortholog; excluded from scoring |

*Table S6. The 45 Purkinje-fiber-enriched loci that lacked a stable gene symbol and a mappable human ortholog in the annotation and were therefore excluded from scoring; they are reported with their differential-expression statistics only (mean |log2 fold-change| across the two comparisons and the smallest adjusted P). Loci are numbered U01-U45.*

#### Table S7. Stability-pillar dictionary: 41 annotation terms (Pillars A-E)

| Pillar | # | Annotation term | Database identifiers / enrichment codes | Scoring direction |
|---|---|---|---|---|
| A | 1 | MAPK/ERK signaling cascade | KEGG hsa04010; Reactome MAPK | inverted |
| A | 2 | PI3K-Akt-mTOR cascade | KEGG hsa04151/hsa04150; Reactome | inverted |
| A | 3 | Wnt/Notch developmental signaling | KEGG hsa04310/hsa04330; Reactome; GO:BP | inverted |
| A | 4 | JAK-STAT / cytokine cross-activation | KEGG hsa04630; Reactome cytokine signaling; WP IL-18 | inverted |
| A | 5 | FoxO / apoptotic stress axis | KEGG hsa04068; Reactome | inverted |
| A | 6 | TGF-beta / BMP morphogen axis | KEGG hsa04350; Reactome; WP | inverted |
| A | 7 | Growth factor / RTK signaling | KEGG hsa04012; Reactome; WP BDNF/VEGF | inverted |
| A | 8 | RAAS / angiotensin axis | KEGG; Reactome; WP RAAS | inverted |
| A | 9 | cAMP / PKA adrenergic signaling | KEGG hsa04024/hsa04261; WP | inverted |
| A | 10 | Ca2+ / calmodulin-CaMK axis | KEGG hsa04020; WP Calcium Regulation in Cardiac Cell; GO:MF calmodulin binding | inverted |
| A | 11 | cGMP-PKG / natriuretic signaling | KEGG hsa04022 | inverted |
| A | 12 | GPCR / G-protein second messenger | Reactome GPCR downstream; KEGG; GO:MF GPCR activity | inverted |
| A | 13 | Rho GTPase / cytoskeletal remodeling | Reactome Rho GTPase cycle; KEGG hsa04810 | inverted |
| B | 1 | miRNA-mediated gene suppression | WP MicroRNAs in Cardiomyocyte Hypertrophy | inverted |
| B | 2 | RNA pol II transcriptional regulation | Reactome RNA Pol II / Generic Transcription; GO:BP transcription by RNA Pol II | inverted |
| B | 3 | Negative transcriptional repression | GO:BP negative regulation of transcription by RNA Pol II | inverted |
| B | 4 | Chromatin / epigenetic remodeling | Reactome chromatin organisation; GO:MF chromatin binding | inverted |
| B | 5 | Post-translational protein modification | Reactome post-translational protein modification; GO:BP protein phosphorylation | inverted |
| B | 6 | Ubiquitin / proteasomal degradation | WP Parkin-Ubiquitin proteasomal degradation; Reactome metabolism of proteins | inverted |
| B | 7 | Autophagy / lysosomal turnover | KEGG hsa04136/hsa04145; Reactome macroautophagy | inverted |
| B | 8 | Cellular stress response | Reactome cellular responses to stress; GO:BP unfolded protein response | inverted |
| C | 1 | Receptor / ligand-dependent activity | GO:MF receptor activity / receptor binding | inverted |
| C | 2 | Kinase phosphorylation substrate | GO:MF protein kinase activity | inverted |
| C | 3 | GTPase-regulated activity | GO:MF GTPase activity; Reactome Rho GTPase cycle | inverted |
| C | 4 | Ca2+-gated / calmodulin-dependent activity | GO:MF calmodulin binding / calcium channel regulator activity | inverted |
| C | 5 | Secreted / extracellularly released | UniProt secreted; Reactome vesicle-mediated transport | inverted |
| C | 6 | Membrane trafficking / endocytosis | KEGG hsa04144; Reactome membrane trafficking | inverted |
| C | 7 | Conditional complex assembly | GO:BP protein-containing complex assembly; Reactome | inverted |
| D | 1 | Cardiac-specific pathway member | KEGG hsa04260/hsa04261; Reactome striated muscle contraction; WP Cardiac Conduction | inverted |
| D | 2 | Neuronal system co-annotation | Reactome neuronal system; KEGG synaptic pathways | inverted |
| D | 3 | Multi-system metabolic membership | Reactome metabolism; KEGG metabolic pathways | inverted |
| D | 4 | Immune / inflammatory co-activation | Reactome immune system; WP IL-18 signaling | inverted |
| D | 5 | Developmental pathway participation | Reactome developmental biology; WP Cardiac Progenitor Differentiation | inverted |
| D | 6 | Hemostasis / vascular cross-pathway | Reactome hemostasis | inverted |
| D | 7 | Multi-disease pathway involvement | KEGG disease maps; Reactome disease | inverted |
| E | 1 | Sarcomere / myofilament terminal effector | Reactome (striated) muscle contraction; KEGG hsa04260; GO:BP sarcomere organisation | direct |
| E | 2 | Constitutive cytoskeletal scaffold | GO:CC cytoskeleton/intermediate filament/actin/microtubule; GO:MF structural constituent of cytoskeleton | direct |

| Pillar | # | Annotation term | Database identifiers / enrichment codes | Scoring direction |
|---|---|---|---|---|
| E | 3 | ECM structural deposit | Reactome ECM organisation; GO:MF ECM structural constituent; MatrisomeDB core matrisome | direct |
| E | 4 | Conduction hardware (channel/pump) | Reactome potassium channels / cardiac conduction; KEGG hsa04540 gap junction | direct |
| E | 5 | Lineage-locking transcription factor | WP Cardiac Progenitor Differentiation; GO:MF DNA-binding TF activity, Pol II-specific | direct |
| E | 6 | Focal adhesion / integrin structural | WP focal adhesion; KEGG hsa04512 ECM-receptor interaction; Reactome hemostasis | direct |

*Table S7. The 41 binary annotation terms underlying the five meta-analytic stability pillars (A-E), with their source database identifiers and scoring direction. Within each pillar the hit count is divided by the per-pillar maximum (A=6, B=5, C=4, D=5, E=3); pillars A-D are inverted (fewer regulatory/turnover annotations indicate greater predicted stability) and pillar E is used directly (greater structural embedding indicates greater stability). Sources: KEGG, Reactome, WikiPathways (WP), Gene Ontology (GO:BP/CC/MF), UniProt and MatrisomeDB.*

#### Table S8. RT-qPCR results (sheep; Purkinje fibers vs cardiomyocytes, n = 4)

| Gene | n (PF/CM) | Mean ΔΔCt | Fold-change (95% CI) | Direction | Welch t (df) | Welch P | MWU P | Call |
|---|---|---|---|---|---|---|---|---|
| **Figure 2 - canonical conduction-system markers** | | | | | | | | |
| - | 4 / 4 | -8.01 | 258.6 (89.4-747.9) | PF | 14.00 (4.4) | <0.0001 | 0.0286 | PF-enriched |
| - | 4 / 4 | -3.28 | 9.7 (3.0-31.7) | PF | 5.95 (3.1) | 0.0083 | 0.0286 | PF-enriched |
| - | 4 / 4 | -5.22 | 37.2 (4.5-303.8) | PF | 5.33 (3.2) | 0.0114 | 0.0286 | PF-enriched |
| - | 4 / 4 | -4.92 | 30.3 (10.3-88.6) | PF | 8.61 (4.3) | 0.0008 | 0.0286 | PF-enriched |
| - | 4 / 4 | -1.01 | 2.0 (1.2-3.5) | PF | 3.10 (6.0) | 0.0214 | 0.0571 | PF-enriched |
| - | 4 / 4 | +0.78 | 1.7 (0.5-5.4) | CM | 1.34 (3.8) | 0.2532 | 0.3429 | Not enriched |
| **Figure 3 - additional contactins** | | | | | | | | |
| *CNTN4* | 4 / 4 | +0.89 | 1.8 (0.7-4.9) | CM | 1.78 (3.8) | 0.1518 | 0.1143 | Not enriched |
| *CNTN5* | 4 / 2 | -8.42 | 343 (146-805) | PF | 19.09 (3.9) | <0.0001 | 0.1333 | PF-enriched |
| **Figure 6 - prioritized new markers** | | | | | | | | |
| *TAGLN* | 4 / 4 | -7.98 | 252.2 (90.0-706.6) | PF | 14.22 (4.5) | <0.0001 | 0.0286 | PF-enriched |
| *CNN1* | 4 / 4 | -6.02 | 64.8 (24.9-168.4) | PF | 10.69 (6.0) | <0.0001 | 0.0286 | PF-enriched |
| *COL19A1* | 4 / 4 | -6.74 | 106.9 (50.5-226.5) | PF | 15.34 (5.8) | <0.0001 | 0.0286 | PF-enriched |
| *RIMS1* | 4 / 4 | -6.83 | 113.7 (25.7-502.3) | PF | 8.05 (5.3) | 0.0004 | 0.0286 | PF-enriched |
| *CAPN3* | 4 / 4 | -5.87 | 58.5 (28.5-119.9) | PF | 14.80 (4.7) | <0.0001 | 0.0286 | PF-enriched |
| *SCN4B* | 4 / 4 | -5.47 | 44.5 (19.5-101.4) | PF | 11.27 (6.0) | <0.0001 | 0.0286 | PF-enriched |
| *PTGR1* | 4 / 4 | -5.23 | 37.6 (17.6-80.2) | PF | 13.45 (3.9) | 0.0002 | 0.0286 | PF-enriched |
| *TES* | 4 / 4 | -4.32 | 19.9 (8.7-45.4) | PF | 10.18 (3.9) | 0.0006 | 0.0286 | PF-enriched |
| *DKK3* | 4 / 4 | -4.50 | 22.7 (5.8-88.4) | PF | 6.55 (3.7) | 0.0036 | 0.0286 | PF-enriched |
| *ACTA2* | 4 / 4 | -3.87 | 14.6 (6.2-34.7) | PF | 7.63 (6.0) | 0.0003 | 0.0286 | PF-enriched |
| *MYL7* | 4 / 4 | -3.73 | 13.2 (3.5-50.0) | PF | 5.18 (4.5) | 0.0048 | 0.0286 | PF-enriched |
| *CSRP1* | 4 / 4 | -2.88 | 7.3 (3.6-14.9) | PF | 6.92 (6.0) | 0.0005 | 0.0286 | PF-enriched |
| *S100A16* | 4 / 4 | -1.35 | 2.6 (1.6-4.0) | PF | 5.02 (5.8) | 0.0027 | 0.0286 | PF-enriched |
| *GYS1* | 4 / 4 | +1.07 | 2.1 (1.0-4.3) | CM | 2.59 (6.0) | 0.0415 | 0.0571 | CM-enriched |
| *THSD4* | 4 / 4 | -0.80 | 1.7 (0.04-68) | PF | 0.38 (5.7) | 0.7211 | 0.4857 | Not enriched |
| *TUBA1A* | 4 / 4 | +2.48 | 5.6 (2.1-14.5) | CM | 4.48 (5.4) | 0.0053 | 0.0286 | CM-enriched |

*Table S8. Full RT-qPCR statistics for Purkinje-fiber marker validation in sheep (Purkinje fibers, PF, versus left-ventricular cardiomyocytes, CM). Relative expression was normalized to the geometric mean of two reference genes (HPRT1 and b2m). Primary test: Welch's unequal-variance t test on ΔCt (two-sided P; Welch P); sensitivity test: two-tailed exact Mann-Whitney U test (two-sided P; MWU P). Mean ΔΔCt = mean ΔCt(PF) - mean ΔCt(CM) in cycles (a negative value indicates enrichment in PF); Fold-change is 2^-ΔΔCt expressed in the direction indicated, with the 95% confidence interval; Welch t (df) gives the test statistic and Welch-Satterthwaite degrees of freedom. n = 4 biological replicates per group (CNTN5: n = 4 PF, n = 2 CM, note that cardiomyocyte expression for 2 / 4 samples were at or below the limit of detection). Significance threshold P < 0.05; no correction for multiplicity.*

**Figure S1. Functional, gene ontology, and disease-association landscape**

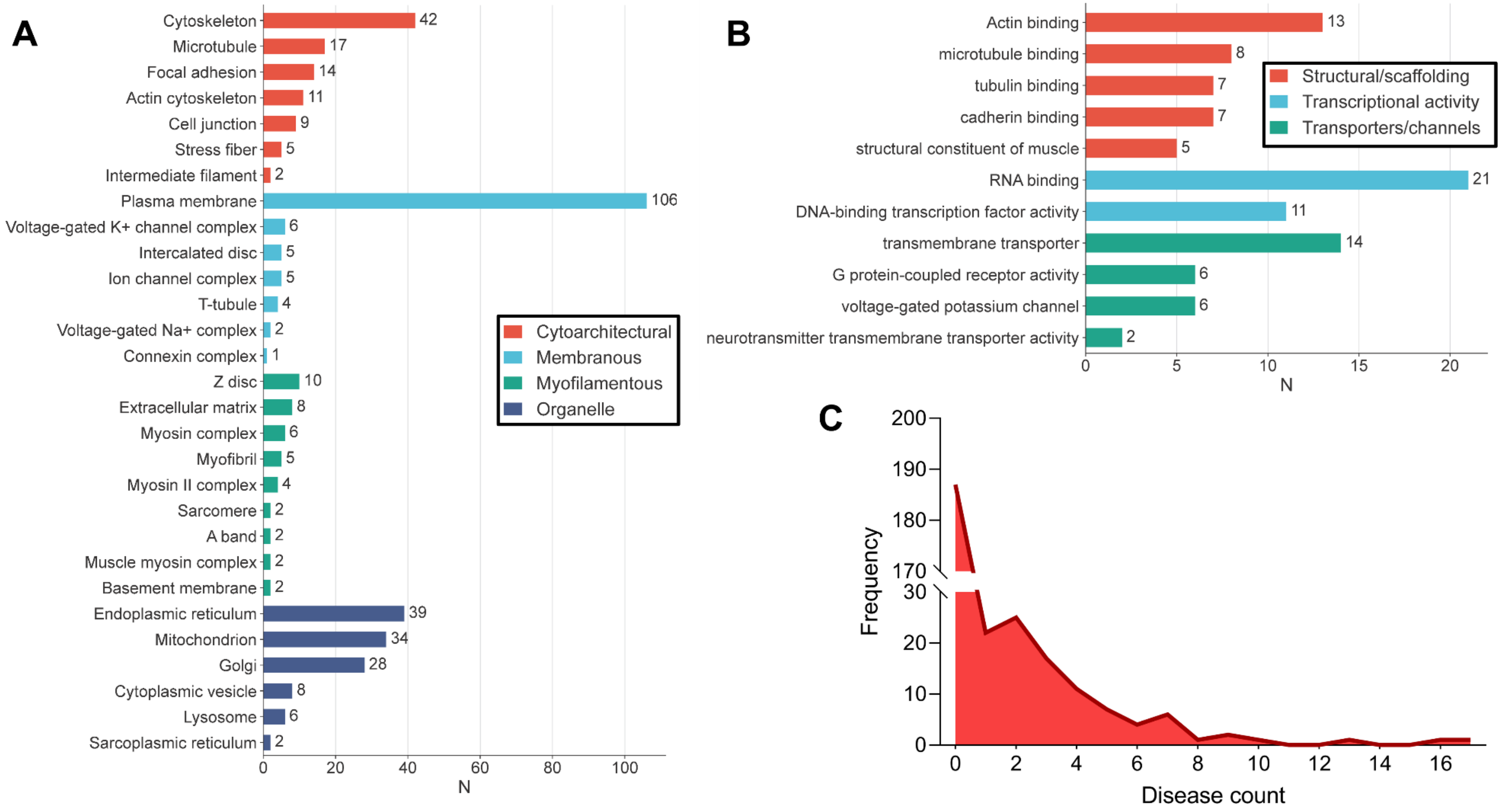


*Figure S1. Functional and disease-association landscape of the Purkinje-enriched gene panel. (A) Number of panel genes annotated to each Gene Ontology cellular-component (subcellular localization) term, grouped as cytoarchitectural, membranous, myofilamentous and organellar categories. (B) Number of panel genes annotated to each Gene Ontology molecular-function term, grouped as structural/scaffolding, transcriptional and transporter/channel activities. (C) Frequency distribution of the number of associated diseases per gene across the panel. Annotation counts are non-exclusive, as a gene may carry several terms.*

#### Figure S2. Conceptual basis of the pathway-based stability scoring

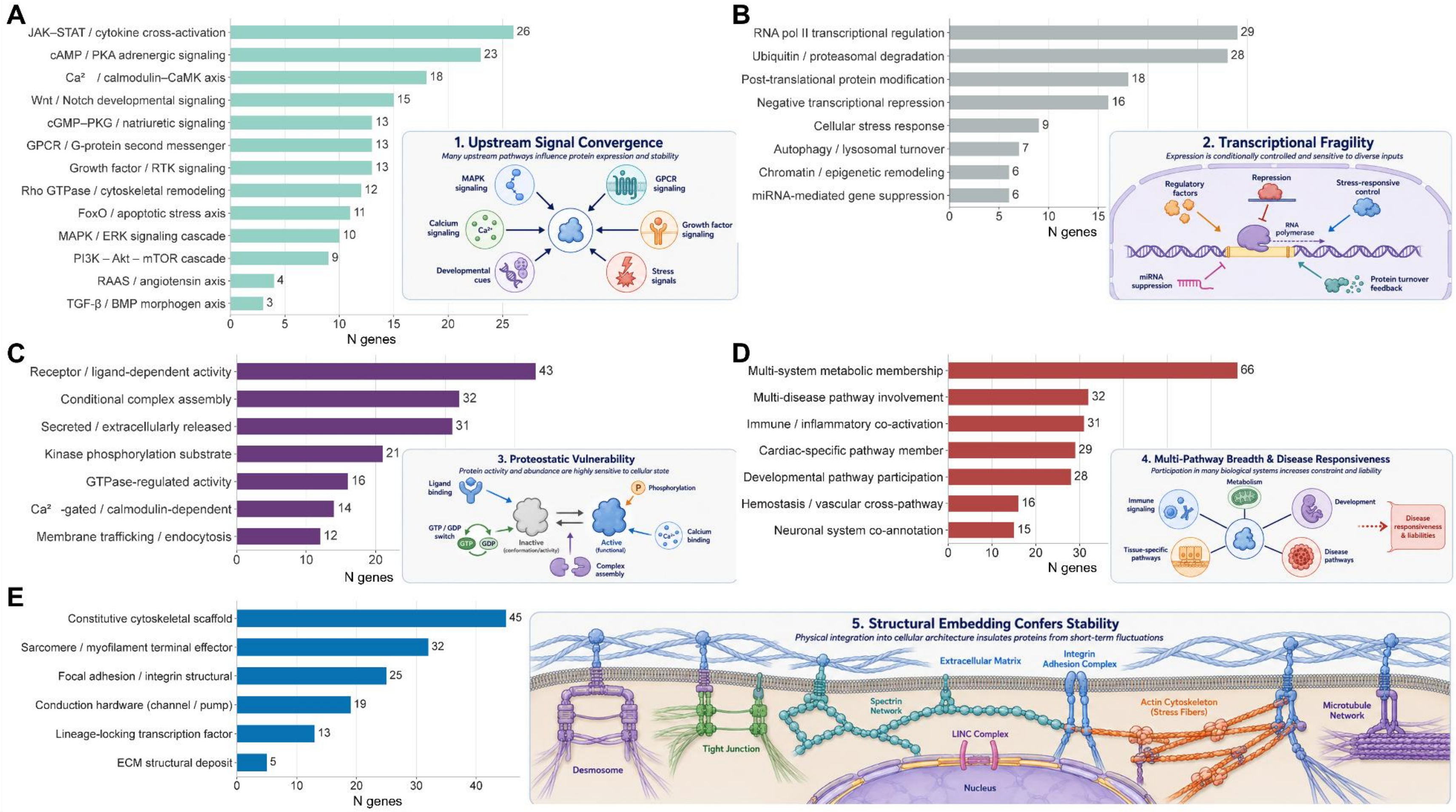


*Figure S2. Conceptual basis of the pathway-based stability scoring. Each of the five pillars (A, upstream signal convergence; B, transcriptional fragility; C, proteostatic vulnerability; D, multi-pathway breadth and disease responsiveness; E, structural grounding) aggregates a defined set of binary annotation terms. For pillars A-D, fewer annotations indicate a transcript whose expression is less likely to be conditionally regulated and therefore more stable; for pillar E, more structural annotations indicate greater stability. The figure also summarizes, per pillar, the number of genes annotated to each term across the marker set. Schematic insets illustrate each stability pillar definitions.*

### Comprehensive marker gene list and pathway membership

#### Table S9. Prioritized Purkinje-fiber marker genes with pathway membership (286 scored genes)

| Rank | Sheep symbol | Human ortholog | Mean log2FC | Min adj. P | Compartment | Class | Stability | Priority | Stability-pillar terms fired (pathway membership) |
|---|---|---|---|---|---|---|---|---|---|
| 1 | CSRP1 | CSRP1 | 3.30 | 5.55e-127 | cell surface | structural | High | 0.9767 | A: Growth factor / RTK signaling ; B: Cellular stress response ; E: Sarcomere / myofilament terminal effector |
| 2 | PTGR1 | PTGR1 | 3.82 | 1.96e-130 | cell surface | metabolic | High | 0.9662 | D: Multi-system metabolic membership |
| 3 | TAGLN | TAGLN | 3.91 | 6.59e-92 | intracellular | structural | High | 0.9580 | |
| 4 | CNN1 | CNN1 | 4.86 | 9.41e-86 | intracellular | structural | High | 0.9514 | A: cAMP / PKA adrenergic signaling ; A: Ca2+ / calmodulin-CaMK axis ; C: Ca2+-gated / calmodulin-dependent activity ; D: Developmental pathway participation ; E: Sarcomere / myofilament terminal effector ; E: Constitutive cytoskeletal scaffold |
| 5 | MYL4 | MYL4 | 6.69 | 7.50e-179 | intracellular | structural | High | 0.9486 | A: cAMP / PKA adrenergic signaling ; D: Cardiac-specific pathway member ; E: Sarcomere / myofilament terminal effector |
| 6 | S100A16 | S100A16 | 1.81 | 3.82e-104 | cell surface | other | High | 0.9305 | C: Ca2+-gated / calmodulin-dependent activity |
| 7 | PRSS23 | PRSS23 | 4.61 | 2.51e-121 | cell surface | metabolic | High | 0.9291 | B: Post-translational protein modification ; B: Ubiquitin / proteasomal degradation ; D: Multi-system metabolic membership |
| 8 | DKK3 | DKK3 | 3.53 | 7.67e-57 | cell surface | regulatory | High | 0.9170 | A: Wnt/Notch developmental signaling ; C: Receptor / ligand-dependent activity ; C: Secreted / extracellularly released |
| 9 | DDR1 | DDR1 | 2.93 | 1.43e-77 | cell surface | regulatory | High | 0.9152 | C: Receptor / ligand-dependent activity ; C: Kinase phosphorylation substrate ; C: Conditional complex assembly ; E: ECM structural deposit |
| 10 | TES | TES | 3.74 | 6.15e-76 | cell surface | structural | High | 0.9141 | |
| 11 | MYL7 | MYL7 | 3.43 | 9.91e-34 | intracellular | structural | High | 0.9041 | A: Rho GTPase / cytoskeletal remodeling ; E: Sarcomere / myofilament terminal effector ; E: Focal adhesion / integrin structural |
| 12 | NES | NES | 3.12 | 5.64e-47 | intracellular | structural | High | 0.9036 | C: Receptor / ligand-dependent activity ; E: Constitutive cytoskeletal scaffold |
| 13 | PON2 | PON2 | 2.05 | 2.45e-99 | cell surface | metabolic | High | 0.8928 | D: Multi-system metabolic membership |
| 14 | TUBA1A | TUBA1A | 2.19 | 9.95e-67 | cell surface | structural | High | 0.8868 | A: JAK-STAT / cytokine cross-activation ; A: FoxO / apoptotic stress axis ; B: Ubiquitin / proteasomal degradation ; D: Neuronal system co-annotation ; D: Immune / inflammatory co-activation ; D: Developmental pathway participation ; D: Hemostasis / vascular cross-pathway ; D: Multi-disease pathway involvement ; E: Constitutive cytoskeletal scaffold ; E: Conduction hardware (channel/pump) ; E: Focal adhesion / integrin structural |
| 15 | DES | DES | 2.39 | 9.89e-27 | cell surface | structural | High | 0.8813 | A: JAK-STAT / cytokine cross-activation ; D: Cardiac-specific pathway member ; D: Immune / inflammatory co-activation ; D: Multi-disease pathway involvement ; E: Sarcomere / myofilament terminal effector ; E: Constitutive cytoskeletal scaffold |
| 16 | BMP5 | BMP5 | 3.34 | 1.83e-53 | cell surface | regulatory | High | 0.8760 | A: TGF-beta / BMP morphogen axis ; C: Receptor / ligand-dependent activity ; C: Secreted / extracellularly released |
| 17 | WFDC10A | WFDC10A | 6.62 | 5.74e-28 | cell surface | regulatory | High | 0.8718 | C: Secreted / extracellularly released |
| 18 | PGM5 | PGM5 | 2.23 | 2.66e-28 | cell surface | structural | High | 0.8707 | C: Conditional complex assembly ; E: Constitutive cytoskeletal scaffold |
| 19 | GYG1 | GYG1 | 2.64 | 6.94e-35 | cell surface | regulatory | High | 0.8627 | A: JAK-STAT / cytokine cross-activation ; D: Multi-system metabolic membership ; D: Immune / inflammatory co-activation ; D: Multi-disease pathway involvement |
| 20 | PPIA | PPIA | 1.77 | 1.19e-62 | cell surface | structural | High | 0.8596 | B: Post-translational protein modification ; C: Kinase phosphorylation substrate ; C: Secreted / extracellularly released |
| 21 | LRP4 | LRP4 | 2.46 | 4.70e-59 | cell surface | regulatory | High | 0.8591 | A: Wnt/Notch developmental signaling ; C: Receptor / ligand-dependent activity ; E: ECM structural deposit |
| 22 | ANXA6 | ANXA6 | 1.60 | 5.33e-42 | cell surface | structural | High | 0.8508 | A: Ca2+ / calmodulin-CaMK axis ; C: Ca2+-gated / calmodulin-dependent activity ; D: Cardiac-specific pathway member ; E: Sarcomere / myofilament terminal effector |
| 23 | MYL9 | MYL9 | 2.84 | 4.23e-24 | intracellular | structural | High | 0.8446 | A: cAMP / PKA adrenergic signaling ; A: cGMP-PKG / natriuretic signaling ; A: Rho GTPase / cytoskeletal remodeling ; C: Conditional complex assembly ; D: Cardiac-specific pathway member ; E: Sarcomere / myofilament terminal effector ; E: Constitutive cytoskeletal scaffold ; E: Focal adhesion / integrin structural |
| 24 | TAPBPL | TAPBPL | 1.93 | 9.80e-53 | cell surface | other | High | 0.8438 | C: Conditional complex assembly |

| Rank | Sheep symbol | Human ortholog | Mean log2FC | Min adj. P | Compartment | Class | Stability | Priority | Stability-pillar terms fired (pathway membership) |
|---|---|---|---|---|---|---|---|---|---|
| 25 | SLC39A8 | SLC39A8 | 2.93 | 1.08e-26 | cell surface | metabolic | High | 0.8406 | |
| 26 | DNTTIP1 | DNTTIP1 | 1.96 | 5.42e-51 | intracellular | regulatory | High | 0.8390 | |
| 27 | PDLIM4 | PDLIM4 | 2.24 | 2.19e-20 | cell surface | structural | High | 0.8364 | E: Constitutive cytoskeletal scaffold |
| 28 | TLE1 | TLE1 | 2.51 | 1.11e-70 | intracellular | regulatory | High | 0.8361 | A: Wnt/Notch developmental signaling |
| 29 | ACTA2 | ACTA2 | 3.41 | 8.22e-45 | cell surface | structural | High | 0.8346 | A: Wnt/Notch developmental signaling ; A: JAK-STAT / cytokine cross-activation ; A: cAMP / PKA adrenergic signaling ; D: Cardiac-specific pathway member ; D: Immune / inflammatory co-activation ; D: Developmental pathway participation ; E: Sarcomere / myofilament terminal effector ; E: Constitutive cytoskeletal scaffold ; E: ECM structural deposit |
| 30 | REC114 | REC114 | 4.71 | 8.20e-55 | intracellular | other | High | 0.8332 | |
| 31 | MYL6 | MYL6 | 1.40 | 1.10e-33 | cell surface | structural | High | 0.8302 | E: Sarcomere / myofilament terminal effector |
| 32 | IGF2R | IGF2R | 1.85 | 1.64e-93 | cell surface | regulatory | High | 0.8279 | A: JAK-STAT / cytokine cross-activation ; B: Autophagy / lysosomal turnover ; C: Receptor / ligand-dependent activity ; C: Secreted / extracellularly released ; C: Membrane trafficking / endocytosis ; D: Immune / inflammatory co-activation |
| 33 | TSPO | TSPO | 1.70 | 6.72e-31 | cell surface | metabolic | High | 0.8245 | C: Receptor / ligand-dependent activity ; D: Multi-system metabolic membership |
| 34 | MAP1A | MAP1A | 2.48 | 2.02e-35 | intracellular | structural | High | 0.8221 | B: Ubiquitin / proteasomal degradation ; E: Constitutive cytoskeletal scaffold |
| 35 | SLC25A23 | SLC25A23 | 2.24 | 1.04e-22 | intracellular | metabolic | High | 0.8205 | |
| 36 | RIMS1 | RIMS1 | 4.21 | 1.63e-30 | cell surface | structural | High | 0.8203 | D: Neuronal system co-annotation |
| 37 | GABARAPL1 | GABARAPL1 | 1.78 | 2.77e-92 | cell surface | structural | High | 0.8195 | A: FoxO / apoptotic stress axis ; B: Autophagy / lysosomal turnover ; C: Receptor / ligand-dependent activity ; C: Conditional complex assembly ; D: Multi-disease pathway involvement ; E: Constitutive cytoskeletal scaffold |
| 38 | OAZ2 | OAZ2 | 1.69 | 7.19e-38 | intracellular | regulatory | High | 0.8173 | D: Multi-system metabolic membership |
| 39 | CALM3 | CALM3 | 1.60 | 3.77e-59 | cell surface | regulatory | High | 0.8162 | A: MAPK/ERK signaling cascade ; A: RAAS / angiotensin axis ; A: cAMP / PKA adrenergic signaling ; A: Ca2+ / calmodulin-CaMK axis ; A: cGMP-PKG / natriuretic signaling ; C: Receptor / ligand-dependent activity ; C: Ca2+-gated / calmodulin-dependent activity ; D: Cardiac-specific pathway member ; D: Hemostasis / vascular cross-pathway ; E: Sarcomere / myofilament terminal effector ; E: Constitutive cytoskeletal scaffold ; E: Focal adhesion / integrin structural |
| 40 | SCN4B | SCN4B | 4.83 | 1.64e-40 | cell surface | metabolic | High | 0.8143 | D: Cardiac-specific pathway member ; D: Developmental pathway participation ; E: Sarcomere / myofilament terminal effector ; E: Conduction hardware (channel/pump) |
| 41 | RAI2 | RAI2 | 2.44 | 6.58e-30 | intracellular | other | High | 0.8092 | |
| 42 | TBX5 | TBX5 | 2.21 | 6.08e-23 | intracellular | regulatory | High | 0.8085 | B: RNA pol II transcriptional regulation ; D: Developmental pathway participation ; E: Lineage-locking transcription factor |
| 43 | UBAC2 | UBAC2 | 1.90 | 1.91e-65 | intracellular | regulatory | High | 0.8071 | A: Wnt/Notch developmental signaling |
| 44 | KIF5B | KIF5B | 2.04 | 1.45e-36 | intracellular | structural | High | 0.8065 | A: Rho GTPase / cytoskeletal remodeling ; C: Membrane trafficking / endocytosis ; C: Conditional complex assembly ; D: Immune / inflammatory co-activation ; D: Hemostasis / vascular cross-pathway ; D: Multi-disease pathway involvement ; E: Constitutive cytoskeletal scaffold ; E: Focal adhesion / integrin structural |
| 45 | CPNE5 | CPNE5 | 4.62 | 6.07e-49 | cell surface | other | Borderline | 0.8005 | |
| 46 | ATP1A2 | ATP1A2 | 3.14 | 3.23e-42 | cell surface | regulatory | High | 0.7974 | A: cAMP / PKA adrenergic signaling ; A: cGMP-PKG / natriuretic signaling ; D: Cardiac-specific pathway member ; D: Multi-disease pathway involvement ; E: Sarcomere / myofilament terminal effector ; E: Conduction hardware (channel/pump) |
| 47 | SLIT2 | SLIT2 | 3.30 | 1.19e-56 | cell surface | structural | High | 0.7956 | B: Post-translational protein modification ; C: Secreted / extracellularly released ; C: Conditional complex assembly ; D: Developmental pathway participation |
| 48 | HAGH | HAGH | 1.94 | 2.85e-23 | intracellular | metabolic | High | 0.7937 | D: Multi-system metabolic membership |
| 49 | VAMP1 | VAMP1 | 3.55 | 4.63e-28 | cell surface | regulatory | High | 0.7935 | C: Receptor / ligand-dependent activity ; C: Conditional complex assembly |
| 50 | DHCR24 | DHCR24 | 2.72 | 4.23e-23 | intracellular | metabolic | High | 0.7927 | D: Multi-system metabolic membership |
| 51 | RRAS2 | RRAS2 | 2.15 | 2.40e-28 | cell surface | regulatory | High | 0.7926 | A: MAPK/ERK signaling cascade ; A: cAMP / PKA adrenergic signaling ; A: Rho GTPase / cytoskeletal remodeling ; B: Autophagy / lysosomal turnover ; C: GTPase-regulated activity |
| 52 | LRATD1 | LRATD1 | 3.72 | 4.64e-39 | intracellular | other | High | 0.7915 | |

| Rank | Sheep symbol | Human ortholog | Mean log2FC | Min adj. P | Compartment | Class | Stability | Priority | Stability-pillar terms fired (pathway membership) |
|---|---|---|---|---|---|---|---|---|---|
| 53 | EPHA4 | EPHA4 | 3.60 | 1.39e-55 | cell surface | regulatory | Borderline | 0.7863 | C: Receptor / ligand-dependent activity ; C: Kinase phosphorylation substrate ; D: Developmental pathway participation |
| 54 | SMPDL3A | SMPDL3A | 3.12 | 4.56e-18 | cell surface | regulatory | High | 0.7861 | C: Secreted / extracellularly released |
| 55 | TSC22D1 | TSC22D1 | 2.41 | 4.16e-44 | cell surface | regulatory | High | 0.7799 | B: RNA pol II transcriptional regulation |
| 56 | CAPN3 | CAPN3 | 4.73 | 1.48e-47 | cell surface | structural | Borderline | 0.7722 | C: Conditional complex assembly |
| 57 | MSRB1 | MSRB1 | 1.78 | 7.12e-39 | intracellular | regulatory | High | 0.7676 | B: Ubiquitin / proteasomal degradation ; D: Multi-system metabolic membership ; E: Constitutive cytoskeletal scaffold |
| 58 | DUSP14 | DUSP14 | 2.48 | 1.97e-57 | unknown | regulatory | High | 0.7674 | |
| 59 | KCNA3 | KCNA3 | 2.50 | 5.40e-32 | cell surface | metabolic | High | 0.7516 | D: Neuronal system co-annotation ; E: Conduction hardware (channel/pump) |
| 60 | ATP2B1 | ATP2B1 | 1.64 | 5.60e-65 | cell surface | regulatory | High | 0.7509 | A: cAMP / PKA adrenergic signaling ; A: Ca2+ / calmodulin-CaMK axis ; A: cGMP-PKG / natriuretic signaling ; C: Ca2+-gated / calmodulin-dependent activity ; D: Cardiac-specific pathway member ; D: Hemostasis / vascular cross-pathway ; D: Multi-disease pathway involvement ; E: Sarcomere / myofilament terminal effector ; E: Conduction hardware (channel/pump) ; E: Focal adhesion / integrin structural |
| 61 | CCDC85C | CCDC85C | 3.30 | 1.36e-35 | unknown | regulatory | High | 0.7506 | A: Wnt/Notch developmental signaling |
| 62 | C22orf39 | C22orf39 | 1.21 | 1.43e-32 | cell surface | regulatory | High | 0.7496 | |
| 63 | KCNJ3 | KCNJ3 | 2.60 | 1.42e-14 | cell surface | metabolic | High | 0.7493 | A: Ca2+ / calmodulin-CaMK axis ; A: GPCR / G-protein second messenger ; D: Cardiac-specific pathway member ; D: Neuronal system co-annotation ; E: Conduction hardware (channel/pump) |
| 64 | FGF12 | FGF12 | 3.25 | 1.38e-42 | cell surface | regulatory | Borderline | 0.7489 | A: MAPK/ERK signaling cascade ; A: PI3K-Akt-mTOR cascade ; C: Receptor / ligand-dependent activity ; D: Cardiac-specific pathway member ; D: Developmental pathway participation ; E: Sarcomere / myofilament terminal effector ; E: Conduction hardware (channel/pump) ; E: Focal adhesion / integrin structural |
| 65 | ALCAM | ALCAM | 2.41 | 3.38e-16 | cell surface | regulatory | High | 0.7430 | C: Receptor / ligand-dependent activity ; C: Secreted / extracellularly released ; D: Developmental pathway participation |
| 66 | OAZ1 | OAZ1 | 1.40 | 2.65e-38 | intracellular | regulatory | High | 0.7427 | D: Multi-system metabolic membership |
| 67 | COX8A | COX8A | 1.66 | 1.07e-23 | intracellular | metabolic | High | 0.7423 | B: RNA pol II transcriptional regulation ; B: Cellular stress response ; D: Cardiac-specific pathway member ; D: Multi-system metabolic membership ; D: Multi-disease pathway involvement ; E: Sarcomere / myofilament terminal effector |
| 68 | SYNDIG1 | SYNDIG1 | 7.01 | 1.41e-29 | cell surface | regulatory | Borderline | 0.7361 | C: Receptor / ligand-dependent activity ; C: Conditional complex assembly |
| 69 | ZNF219 | ZNF219 | 1.99 | 3.16e-23 | intracellular | regulatory | High | 0.7322 | A: JAK-STAT / cytokine cross-activation ; B: RNA pol II transcriptional regulation ; B: Negative transcriptional repression ; C: Receptor / ligand-dependent activity ; D: Immune / inflammatory co-activation ; E: Lineage-locking transcription factor |
| 70 | LGR4 | LGR4 | 1.97 | 2.27e-54 | cell surface | regulatory | Borderline | 0.7316 | A: Wnt/Notch developmental signaling ; A: GPCR / G-protein second messenger ; B: Ubiquitin / proteasomal degradation ; C: Receptor / ligand-dependent activity |
| 71 | ANAPC1 | ANAPC1 | 2.45 | 8.49e-48 | intracellular | regulatory | High | 0.7315 | A: FoxO / apoptotic stress axis ; B: Ubiquitin / proteasomal degradation ; D: Immune / inflammatory co-activation ; D: Multi-disease pathway involvement |
| 72 | GPR4 | GPR4 | 2.87 | 6.87e-22 | cell surface | regulatory | High | 0.7313 | A: GPCR / G-protein second messenger ; C: Receptor / ligand-dependent activity |
| 73 | SNX21 | SNX21 | 1.75 | 2.21e-29 | intracellular | other | High | 0.7266 | |
| 74 | ASS1 | ASS1 | 3.92 | 4.62e-33 | cell surface | metabolic | Borderline | 0.7239 | D: Multi-system metabolic membership |
| 75 | RASL12 | RASL12 | 1.95 | 2.12e-23 | unknown | metabolic | High | 0.7212 | C: GTPase-regulated activity |
| 76 | IGFBP2 | IGFBP2 | 5.55 | 1.48e-48 | cell surface | regulatory | Borderline | 0.7182 | A: cAMP / PKA adrenergic signaling ; B: Ubiquitin / proteasomal degradation ; C: Receptor / ligand-dependent activity ; D: Multi-system metabolic membership |
| 77 | UBE2H | UBE2H | 1.37 | 3.10e-37 | intracellular | metabolic | High | 0.7157 | B: Post-translational protein modification ; B: Ubiquitin / proteasomal degradation ; D: Multi-system metabolic membership ; D: Immune / inflammatory co-activation |
| 78 | IL20RA | IL20RA | 3.32 | 1.79e-21 | cell surface | regulatory | Borderline | 0.6981 | A: JAK-STAT / cytokine cross-activation ; C: Receptor / ligand-dependent activity ; C: Conditional complex assembly ; D: Immune / inflammatory co-activation |
| 79 | DNAJB6 | DNAJB6 | 1.33 | 8.65e-24 | intracellular | structural | High | 0.6955 | C: Conditional complex assembly |
| 80 | HEBP2 | HEBP2 | 1.38 | 3.53e-37 | cell surface | other | High | 0.6946 | A: JAK-STAT / cytokine cross-activation ; D: Immune / inflammatory co-activation |
| 81 | MSX2 | MSX2 | 3.97 | 6.46e-33 | intracellular | regulatory | Borderline | 0.6911 | B: RNA pol II transcriptional regulation ; B: Negative transcriptional repression ; E: Lineage-locking transcription factor |

| Rank | Sheep symbol | Human ortholog | Mean log2FC | Min adj. P | Compartment | Class | Stability | Priority | Stability-pillar terms fired (pathway membership) |
|---|---|---|---|---|---|---|---|---|---|
| 82 | UCHL1 | UCHL1 | 5.01 | 4.83e-55 | cell surface | regulatory | Borderline | 0.6888 | B: Post-translational protein modification ; B: Ubiquitin / proteasomal degradation ; C: Receptor / ligand-dependent activity ; D: Multi-system metabolic membership ; D: Multi-disease pathway involvement |
| 83 | FADS3 | FADS3 | 1.88 | 4.42e-16 | intracellular | metabolic | High | 0.6887 | |
| 84 | CCN2 | CCN2 | 3.02 | 4.72e-65 | cell surface | regulatory | Borderline | 0.6843 | A: Growth factor / RTK signaling ; B: RNA pol II transcriptional regulation ; C: Secreted / extracellularly released |
| 85 | MBOAT2 | MBOAT2 | 3.49 | 2.73e-46 | intracellular | metabolic | Borderline | 0.6728 | D: Multi-system metabolic membership |
| 86 | KLHL32 | KLHL32 | 3.36 | 1.26e-41 | unknown | other | Borderline | 0.6700 | |
| 87 | NPFFR2 | NPFFR2 | 5.28 | 2.76e-17 | cell surface | regulatory | Borderline | 0.6637 | A: GPCR / G-protein second messenger ; C: Receptor / ligand-dependent activity ; E: Constitutive cytoskeletal scaffold |
| 88 | ZNF787 | ZNF787 | 1.50 | 4.62e-33 | intracellular | regulatory | High | 0.6629 | E: Lineage-locking transcription factor |
| 89 | ENO1 | ENO1 | 1.40 | 1.41e-24 | cell surface | regulatory | High | 0.6611 | A: JAK-STAT / cytokine cross-activation ; B: RNA pol II transcriptional regulation ; B: Negative transcriptional repression ; D: Multi-system metabolic membership ; D: Immune / inflammatory co-activation ; D: Multi-disease pathway involvement ; E: Sarcomere / myofilament terminal effector |
| 90 | PREPL | PREPL | 1.45 | 9.05e-30 | intracellular | structural | High | 0.6584 | E: Constitutive cytoskeletal scaffold |
| 91 | THSD4 | THSD4 | 5.41 | 1.20e-54 | cell surface | structural | Borderline | 0.6555 | B: Post-translational protein modification ; B: Ubiquitin / proteasomal degradation ; C: Conditional complex assembly ; D: Multi-system metabolic membership ; D: Multi-disease pathway involvement |
| 92 | SCD | SCD | 1.97 | 1.20e-17 | intracellular | metabolic | High | 0.6531 | B: Chromatin / epigenetic remodeling ; D: Multi-system metabolic membership |
| 93 | CAB39 | CAB39 | 1.81 | 9.23e-18 | cell surface | regulatory | High | 0.6502 | A: PI3K-Akt-mTOR cascade ; A: JAK-STAT / cytokine cross-activation ; C: Kinase phosphorylation substrate ; D: Immune / inflammatory co-activation ; E: Focal adhesion / integrin structural |
| 94 | PAK4 | PAK4 | 1.63 | 1.38e-18 | intracellular | structural | High | 0.6466 | A: MAPK/ERK signaling cascade ; A: Growth factor / RTK signaling ; A: Rho GTPase / cytoskeletal remodeling ; C: Kinase phosphorylation substrate ; C: Conditional complex assembly ; E: Focal adhesion / integrin structural |
| 95 | EFHD2 | EFHD2 | 2.16 | 1.18e-24 | unknown | structural | Borderline | 0.6410 | A: Rho GTPase / cytoskeletal remodeling ; C: GTPase-regulated activity |
| 96 | KCNA5 | KCNA5 | 3.67 | 1.25e-26 | cell surface | regulatory | Borderline | 0.6368 | A: Wnt/Notch developmental signaling ; C: Receptor / ligand-dependent activity ; C: Conditional complex assembly ; D: Cardiac-specific pathway member ; D: Neuronal system co-annotation ; E: Sarcomere / myofilament terminal effector ; E: Conduction hardware (channel/pump) |
| 97 | SHB | SHB | 2.07 | 1.26e-17 | cell surface | regulatory | Borderline | 0.6364 | A: Growth factor / RTK signaling |
| 98 | ODC1 | ODC1 | 1.58 | 1.22e-24 | intracellular | metabolic | High | 0.6331 | D: Multi-system metabolic membership |
| 99 | DSTN | DSTN | 1.37 | 8.18e-28 | cell surface | structural | High | 0.6313 | E: Constitutive cytoskeletal scaffold |
| 100 | RAMP1 | RAMP1 | 2.27 | 4.69e-16 | cell surface | regulatory | Borderline | 0.6306 | A: cAMP / PKA adrenergic signaling ; A: GPCR / G-protein second messenger ; C: Receptor / ligand-dependent activity |
| 101 | MID1IP1 | MID1IP1 | 1.79 | 3.81e-30 | intracellular | metabolic | Borderline | 0.6275 | D: Multi-system metabolic membership ; E: Constitutive cytoskeletal scaffold |
| 102 | MGLL | MGLL | 1.86 | 3.89e-09 | cell surface | metabolic | High | 0.6249 | B: Chromatin / epigenetic remodeling ; D: Multi-system metabolic membership ; D: Hemostasis / vascular cross-pathway ; E: Focal adhesion / integrin structural |
| 103 | ADM | ADM | 2.47 | 1.01e-10 | cell surface | regulatory | High | 0.6229 | A: cAMP / PKA adrenergic signaling ; A: GPCR / G-protein second messenger ; B: Cellular stress response ; C: Receptor / ligand-dependent activity ; C: Secreted / extracellularly released ; E: Sarcomere / myofilament terminal effector |
| 104 | IRX1 | IRX1 | 3.51 | 2.85e-13 | intracellular | regulatory | Borderline | 0.6219 | B: RNA pol II transcriptional regulation ; B: Negative transcriptional repression ; D: Developmental pathway participation ; E: Lineage-locking transcription factor |
| 105 | SCN5A | SCN5A | 1.86 | 9.24e-07 | cell surface | regulatory | High | 0.6200 | A: Ca2+ / calmodulin-CaMK axis ; C: Ca2+-gated / calmodulin-dependent activity ; D: Cardiac-specific pathway member ; D: Developmental pathway participation ; E: Sarcomere / myofilament terminal effector ; E: Conduction hardware (channel/pump) ; E: Lineage-locking transcription factor |
| 106 | MAGED2 | MAGED2 | 1.85 | 1.53e-34 | cell surface | other | Borderline | 0.6191 | B: RNA pol II transcriptional regulation ; B: Negative transcriptional repression ; D: Hemostasis / vascular cross-pathway ; E: Focal adhesion / integrin structural |
| 107 | SLC9A2 | SLC9A2 | 6.30 | 1.76e-27 | cell surface | other | Borderline | 0.6169 | |
| 108 | STAP2 | STAP2 | 1.98 | 2.88e-14 | cell surface | regulatory | High | 0.6141 | B: Cellular stress response |

| Rank | Sheep symbol | Human ortholog | Mean log2FC | Min adj. P | Compartment | Class | Stability | Priority | Stability-pillar terms fired (pathway membership) |
|---|---|---|---|---|---|---|---|---|---|
| 109 | SCD5 | SCD5 | 3.29 | 1.72e-37 | intracellular | metabolic | Borderline | 0.6078 | B: Chromatin / epigenetic remodeling ; D: Multi-system metabolic membership ; D: Developmental pathway participation |
| 110 | AVPI1 | AVPI1 | 1.28 | 3.39e-20 | intracellular | other | High | 0.6051 | |
| 111 | PDPN | PDPN | 4.21 | 7.49e-35 | cell surface | regulatory | Borderline | 0.6017 | C: Receptor / ligand-dependent activity ; D: Hemostasis / vascular cross-pathway ; E: Focal adhesion / integrin structural |
| 112 | S100A10 | S100A10 | 2.22 | 1.50e-31 | cell surface | structural | Borderline | 0.6004 | B: RNA pol II transcriptional regulation ; C: Ca2+-gated / calmodulin-dependent activity ; C: Conditional complex assembly ; D: Hemostasis / vascular cross-pathway ; E: Focal adhesion / integrin structural |
| 113 | HTATIP2 | HTATIP2 | 2.84 | 1.03e-27 | intracellular | regulatory | Borderline | 0.5987 | B: RNA pol II transcriptional regulation ; C: Kinase phosphorylation substrate |
| 114 | ZPBP2 | ZPBP2 | 4.27 | 4.23e-24 | cell surface | regulatory | Borderline | 0.5986 | C: Secreted / extracellularly released ; C: Conditional complex assembly |
| 115 | MASP1 | MASP1 | 1.92 | 5.26e-20 | cell surface | metabolic | Borderline | 0.5981 | C: Ca2+-gated / calmodulin-dependent activity ; C: Secreted / extracellularly released ; D: Multi-disease pathway involvement |
| 116 | NKX2-5 | NKX2-5 | 1.72 | 9.48e-18 | intracellular | regulatory | High | 0.5952 | B: RNA pol II transcriptional regulation ; B: Negative transcriptional repression ; D: Cardiac-specific pathway member ; D: Developmental pathway participation ; E: Sarcomere / myofilament terminal effector ; E: Conduction hardware (channel/pump) ; E: Lineage-locking transcription factor |
| 117 | RBPMS2 | RBPMS2 | 1.96 | 4.09e-06 | intracellular | regulatory | High | 0.5921 | |
| 118 | ADAM19 | ADAM19 | 2.37 | 2.16e-17 | cell surface | structural | Borderline | 0.5909 | E: ECM structural deposit |
| 119 | PCDH19 | PCDH19 | 4.23 | 3.81e-32 | cell surface | structural | Borderline | 0.5880 | D: Developmental pathway participation |
| 120 | ACVR1 | ACVR1 | 1.92 | 4.42e-36 | cell surface | regulatory | Borderline | 0.5877 | A: TGF-beta / BMP morphogen axis ; C: Receptor / ligand-dependent activity ; C: Kinase phosphorylation substrate ; D: Developmental pathway participation |
| 121 | KCNH2 | KCNH2 | 1.46 | 1.72e-17 | cell surface | metabolic | Borderline | 0.5865 | A: JAK-STAT / cytokine cross-activation ; D: Cardiac-specific pathway member ; D: Neuronal system co-annotation ; D: Immune / inflammatory co-activation ; E: Sarcomere / myofilament terminal effector ; E: Conduction hardware (channel/pump) |
| 122 | SPTSSB | SPTSSB | 3.88 | 8.46e-28 | intracellular | metabolic | Borderline | 0.5860 | D: Multi-system metabolic membership |
| 123 | KCNG2 | KCNG2 | 2.17 | 6.18e-24 | cell surface | regulatory | Borderline | 0.5839 | D: Neuronal system co-annotation ; D: Multi-system metabolic membership ; E: Conduction hardware (channel/pump) |
| 124 | SMIM29 | SMIM29 | 1.54 | 5.99e-15 | intracellular | other | High | 0.5829 | |
| 125 | BCAT1 | BCAT1 | 6.08 | 1.27e-27 | intracellular | regulatory | Borderline | 0.5817 | D: Multi-system metabolic membership |
| 126 | RASL10B | RASL10B | 2.90 | 2.46e-28 | cell surface | regulatory | Borderline | 0.5810 | C: GTPase-regulated activity |
| 127 | MYO1G | MYO1G | 2.50 | 1.74e-14 | cell surface | structural | Borderline | 0.5776 | A: Ca2+ / calmodulin-CaMK axis ; C: Ca2+-gated / calmodulin-dependent activity ; E: Constitutive cytoskeletal scaffold |
| 128 | NTF3 | NTF3 | 3.22 | 5.43e-28 | cell surface | regulatory | Borderline | 0.5748 | A: MAPK/ERK signaling cascade ; A: PI3K-Akt-mTOR cascade ; A: Growth factor / RTK signaling ; C: Receptor / ligand-dependent activity ; C: Kinase phosphorylation substrate ; C: Secreted / extracellularly released ; D: Multi-disease pathway involvement |
| 129 | CCDC112 | CCDC112 | 1.68 | 6.87e-11 | intracellular | structural | High | 0.5729 | E: Constitutive cytoskeletal scaffold |
| 130 | CERS4 | CERS4 | 1.44 | 2.21e-27 | intracellular | metabolic | Borderline | 0.5713 | D: Multi-system metabolic membership |
| 131 | TRMT112 | TRMT112 | 1.03 | 3.20e-17 | intracellular | regulatory | High | 0.5705 | B: Ubiquitin / proteasomal degradation ; D: Multi-system metabolic membership |
| 132 | PDE4D | PDE4D | 1.75 | 2.74e-18 | cell surface | regulatory | Borderline | 0.5678 | A: cAMP / PKA adrenergic signaling ; A: GPCR / G-protein second messenger ; C: Ca2+-gated / calmodulin-dependent activity ; D: Multi-system metabolic membership ; E: Constitutive cytoskeletal scaffold |
| 133 | NACC2 | NACC2 | 1.82 | 2.63e-19 | intracellular | regulatory | Borderline | 0.5672 | B: RNA pol II transcriptional regulation ; B: Negative transcriptional repression ; E: Lineage-locking transcription factor |
| 134 | RASSF3 | RASSF3 | 2.03 | 2.18e-10 | cell surface | structural | Borderline | 0.5591 | E: Constitutive cytoskeletal scaffold |
| 135 | PARM1 | PARM1 | 1.86 | 1.50e-13 | cell surface | other | High | 0.5528 | |
| 136 | SVBP | SVBP | 1.40 | 1.31e-32 | cell surface | structural | Borderline | 0.5514 | B: Post-translational protein modification ; B: Ubiquitin / proteasomal degradation ; C: Secreted / extracellularly released ; D: Multi-system metabolic membership ; E: Constitutive cytoskeletal scaffold |
| 137 | TNFSF12 | TNFSF12 | 1.42 | 9.07e-36 | cell surface | regulatory | Borderline | 0.5489 | A: JAK-STAT / cytokine cross-activation ; C: Receptor / ligand-dependent activity ; C: Secreted / extracellularly released ; D: Immune / inflammatory co-activation |

| Rank | Sheep symbol | Human ortholog | Mean log2FC | Min adj. P | Compartment | Class | Stability | Priority | Stability-pillar terms fired (pathway membership) |
|---|---|---|---|---|---|---|---|---|---|
| 138 | SLC17A7 | SLC17A7 | 5.67 | 6.54e-22 | cell surface | metabolic | Lower | 0.5470 | D: Neuronal system co-annotation |
| 139 | RTN1 | RTN1 | 5.88 | 4.01e-23 | intracellular | regulatory | Lower | 0.5450 | |
| 140 | STYXL2 | STYXL2 | 1.81 | 2.18e-28 | intracellular | structural | High | 0.5437 | |
| 141 | HCN4 | HCN4 | 3.29 | 1.38e-34 | cell surface | metabolic | Lower | 0.5424 | A: cAMP / PKA adrenergic signaling ; D: Neuronal system co-annotation ; E: Sarcomere / myofilament terminal effector ; E: Conduction hardware (channel/pump) |
| 142 | MYOT | MYOT | 2.79 | 3.88e-12 | cell surface | structural | Borderline | 0.5422 | C: Receptor / ligand-dependent activity ; E: Sarcomere / myofilament terminal effector ; E: Constitutive cytoskeletal scaffold |
| 143 | COL19A1 | COL19A1 | 5.99 | 5.06e-26 | cell surface | structural | Lower | 0.5413 | C: Secreted / extracellularly released ; E: ECM structural deposit |
| 144 | FUT9 | FUT9 | 5.63 | 5.65e-16 | intracellular | metabolic | Borderline | 0.5404 | D: Multi-system metabolic membership |
| 145 | PLCB2 | PLCB2 | 3.38 | 3.49e-37 | intracellular | regulatory | Lower | 0.5402 | A: RAAS / angiotensin axis ; A: cAMP / PKA adrenergic signaling ; A: Ca2+ / calmodulin-CaMK axis ; A: cGMP-PKG / natriuretic signaling ; B: miRNA-mediated gene suppression ; D: Cardiac-specific pathway member ; D: Neuronal system co-annotation ; D: Multi-system metabolic membership ; E: Sarcomere / myofilament terminal effector |
| 146 | IRX2 | IRX2 | 2.83 | 7.45e-08 | intracellular | regulatory | Borderline | 0.5378 | B: RNA pol II transcriptional regulation ; B: Negative transcriptional repression ; D: Developmental pathway participation ; E: Lineage-locking transcription factor |
| 147 | CD99L2 | CD99L2 | 1.42 | 4.47e-16 | cell surface | other | Borderline | 0.5373 | C: Secreted / extracellularly released ; D: Hemostasis / vascular cross-pathway ; E: Focal adhesion / integrin structural |
| 148 | IDS | IDS | 1.30 | 7.74e-21 | intracellular | metabolic | Borderline | 0.5307 | B: Autophagy / lysosomal turnover ; D: Multi-system metabolic membership ; D: Multi-disease pathway involvement |
| 149 | TLE4 | TLE4 | 2.58 | 2.03e-44 | intracellular | regulatory | Lower | 0.5271 | A: Wnt/Notch developmental signaling ; B: RNA pol II transcriptional regulation ; B: Negative transcriptional repression ; B: Chromatin / epigenetic remodeling |
| 150 | ME3 | ME3 | 1.80 | 9.22e-11 | intracellular | metabolic | High | 0.5245 | D: Multi-system metabolic membership |
| 151 | TMEM130 | TMEM130 | 3.46 | 2.80e-10 | cell surface | other | Borderline | 0.5204 | |
| 152 | PKIB | PKIB | 2.96 | 5.91e-35 | intracellular | regulatory | Lower | 0.5195 | A: cAMP / PKA adrenergic signaling ; A: Ca2+ / calmodulin-CaMK axis ; D: Cardiac-specific pathway member |
| 153 | ADISSP | ADISSP | 1.33 | 4.48e-18 | cell surface | regulatory | Borderline | 0.5184 | C: Secreted / extracellularly released |
| 154 | BMERB1 | BMERB1 | 3.59 | 2.09e-30 | intracellular | structural | Lower | 0.5150 | E: Constitutive cytoskeletal scaffold |
| 155 | SFRP5 | SFRP5 | 3.80 | 2.44e-23 | cell surface | regulatory | Lower | 0.5124 | A: Wnt/Notch developmental signaling ; B: RNA pol II transcriptional regulation ; C: Secreted / extracellularly released |
| 156 | UBE2QL1 | UBE2QL1 | 1.85 | 4.22e-09 | cell surface | metabolic | Borderline | 0.5121 | B: Ubiquitin / proteasomal degradation |
| 157 | VGLL2 | VGLL2 | 4.69 | 3.64e-19 | intracellular | regulatory | Lower | 0.5071 | B: RNA pol II transcriptional regulation |
| 158 | EML2 | EML2 | 1.87 | 4.68e-13 | intracellular | structural | Borderline | 0.5055 | B: Post-translational protein modification ; C: Kinase phosphorylation substrate ; E: Constitutive cytoskeletal scaffold |
| 159 | HK1 | HK1 | 1.44 | 7.68e-23 | intracellular | regulatory | Borderline | 0.5039 | B: Ubiquitin / proteasomal degradation ; D: Multi-system metabolic membership ; D: Multi-disease pathway involvement ; E: Conduction hardware (channel/pump) |
| 160 | SCUBE2 | SCUBE2 | 2.64 | 2.48e-32 | cell surface | other | Lower | 0.5029 | C: Secreted / extracellularly released |
| 161 | CHRM1 | CHRM1 | 3.57 | 1.18e-21 | cell surface | regulatory | Lower | 0.5009 | A: PI3K-Akt-mTOR cascade ; A: cAMP / PKA adrenergic signaling ; A: Ca2+ / calmodulin-CaMK axis ; A: GPCR / G-protein second messenger ; A: Rho GTPase / cytoskeletal remodeling ; C: Receptor / ligand-dependent activity ; D: Cardiac-specific pathway member ; D: Multi-disease pathway involvement ; E: Focal adhesion / integrin structural |
| 162 | SERINC2 | SERINC2 | 1.67 | 1.95e-12 | cell surface | metabolic | High | 0.4995 | D: Multi-system metabolic membership |
| 163 | TUBB2B | TUBB2B | 1.71 | 7.75e-23 | intracellular | structural | Borderline | 0.4985 | A: JAK-STAT / cytokine cross-activation ; B: Ubiquitin / proteasomal degradation ; C: GTPase-regulated activity ; D: Neuronal system co-annotation ; D: Immune / inflammatory co-activation ; D: Developmental pathway participation ; D: Hemostasis / vascular cross-pathway ; D: Multi-disease pathway involvement ; E: Constitutive cytoskeletal scaffold ; E: Conduction hardware (channel/pump) ; E: Focal adhesion / integrin structural |
| 164 | SDF2L1 | SDF2L1 | 1.70 | 5.20e-25 | intracellular | regulatory | Borderline | 0.4974 | |
| 165 | GJA5 | GJA5 | 2.30 | 3.02e-12 | cell surface | structural | Borderline | 0.4968 | A: Ca2+ / calmodulin-CaMK axis ; C: Secreted / extracellularly released ; C: Membrane trafficking / endocytosis ; C: Conditional complex assembly ; D: Cardiac-specific pathway member ; E: Conduction hardware (channel/pump) |

| Rank | Sheep symbol | Human ortholog | Mean log2FC | Min adj. P | Compartment | Class | Stability | Priority | Stability-pillar terms fired (pathway membership) |
|---|---|---|---|---|---|---|---|---|---|
| 166 | BLVRA | BLVRA | 1.96 | 2.96e-17 | cell surface | metabolic | Borderline | 0.4960 | B: Cellular stress response ; D: Multi-system metabolic membership |
| 167 | EIF1B | EIF1B | 1.44 | 6.44e-17 | unknown | metabolic | High | 0.4926 | |
| 168 | CDS2 | CDS2 | 1.40 | 1.94e-17 | intracellular | metabolic | Borderline | 0.4919 | D: Multi-system metabolic membership |
| 169 | SHF | SHF | 1.96 | 6.43e-23 | intracellular | regulatory | Borderline | 0.4901 | |
| 170 | STEAP3 | STEAP3 | 2.02 | 6.64e-13 | cell surface | metabolic | Borderline | 0.4900 | A: Rho GTPase / cytoskeletal remodeling ; B: RNA pol II transcriptional regulation ; C: GTPase-regulated activity |
| 171 | ACOX2 | ACOX2 | 2.02 | 1.83e-13 | intracellular | metabolic | Borderline | 0.4860 | D: Multi-system metabolic membership |
| 172 | KHDRBS3 | KHDRBS3 | 3.29 | 4.91e-26 | intracellular | regulatory | Lower | 0.4809 | |
| 173 | SLC4A3 | SLC4A3 | 1.63 | 3.77e-25 | cell surface | metabolic | Borderline | 0.4808 | |
| 174 | MAPT | MAPT | 1.93 | 1.77e-13 | cell surface | structural | Borderline | 0.4799 | A: MAPK/ERK signaling cascade ; A: Growth factor / RTK signaling ; B: Negative transcriptional repression ; C: Conditional complex assembly ; D: Multi-disease pathway involvement ; E: Constitutive cytoskeletal scaffold |
| 175 | GNAZ | GNAZ | 1.75 | 4.88e-07 | cell surface | regulatory | Borderline | 0.4767 | A: Ca2+ / calmodulin-CaMK axis ; A: GPCR / G-protein second messenger ; C: Receptor / ligand-dependent activity ; C: GTPase-regulated activity ; D: Cardiac-specific pathway member ; D: Multi-disease pathway involvement |
| 176 | NKAIN1 | NKAIN1 | 2.68 | 6.20e-07 | cell surface | structural | Borderline | 0.4743 | |
| 177 | GORASP2 | GORASP2 | 1.16 | 3.12e-16 | intracellular | other | Borderline | 0.4688 | A: FoxO / apoptotic stress axis ; C: Conditional complex assembly |
| 178 | RD3L | RD3L | 3.67 | 1.22e-15 | unknown | other | Lower | 0.4633 | |
| 179 | WDR97 | WDR97 | 1.24 | 8.26e-17 | unknown | metabolic | Borderline | 0.4622 | B: Post-translational protein modification ; C: Kinase phosphorylation substrate |
| 180 | PRKAG2 | PRKAG2 | 1.65 | 1.45e-15 | cell surface | regulatory | Borderline | 0.4612 | A: JAK-STAT / cytokine cross-activation ; A: FoxO / apoptotic stress axis ; C: Kinase phosphorylation substrate |
| 181 | CITED2 | CITED2 | 1.41 | 1.55e-11 | intracellular | regulatory | Borderline | 0.4605 | B: RNA pol II transcriptional regulation ; B: Negative transcriptional repression ; B: Chromatin / epigenetic remodeling ; B: Cellular stress response |
| 182 | IRAG1 | IRAG1 | 1.81 | 9.45e-17 | intracellular | regulatory | Borderline | 0.4587 | A: cGMP-PKG / natriuretic signaling ; E: Sarcomere / myofilament terminal effector |
| 183 | RTN4 | RTN4 | 1.26 | 1.96e-10 | cell surface | structural | Borderline | 0.4553 | |
| 184 | GFRA4 | GFRA4 | 2.50 | 7.16e-27 | cell surface | regulatory | Lower | 0.4546 | C: Receptor / ligand-dependent activity ; C: Secreted / extracellularly released ; D: Developmental pathway participation |
| 185 | NUDT16 | NUDT16 | 1.72 | 9.91e-13 | intracellular | metabolic | Borderline | 0.4481 | D: Multi-system metabolic membership |
| 186 | CCDC158 | CCDC158 | 2.98 | 1.90e-11 | intracellular | other | Borderline | 0.4447 | |
| 187 | ACTR1B | ACTR1B | 1.41 | 1.31e-17 | cell surface | structural | Borderline | 0.4369 | A: JAK-STAT / cytokine cross-activation ; D: Immune / inflammatory co-activation ; D: Multi-disease pathway involvement ; E: Constitutive cytoskeletal scaffold |
| 188 | SPG21 | SPG21 | 1.57 | 8.42e-17 | intracellular | regulatory | Borderline | 0.4359 | C: Receptor / ligand-dependent activity |
| 189 | MYH6 | MYH6 | 2.56 | 8.25e-13 | intracellular | structural | Borderline | 0.4334 | A: JAK-STAT / cytokine cross-activation ; A: cAMP / PKA adrenergic signaling ; A: Ca2+ / calmodulin-CaMK axis ; A: cGMP-PKG / natriuretic signaling ; C: Ca2+-gated / calmodulin-dependent activity ; D: Cardiac-specific pathway member ; D: Immune / inflammatory co-activation ; D: Developmental pathway participation ; D: Multi-disease pathway involvement ; E: Sarcomere / myofilament terminal effector ; E: Lineage-locking transcription factor |
| 190 | MSRA | MSRA | 1.66 | 1.71e-11 | cell surface | metabolic | Borderline | 0.4323 | B: Ubiquitin / proteasomal degradation ; D: Multi-system metabolic membership |
| 191 | SYPL2 | SYPL2 | 2.29 | 9.05e-06 | unknown | other | Borderline | 0.4304 | |
| 192 | TNFRSF13B | TNFRSF13B | 2.79 | 8.50e-18 | cell surface | regulatory | Lower | 0.4279 | A: JAK-STAT / cytokine cross-activation ; C: Receptor / ligand-dependent activity ; D: Immune / inflammatory co-activation |
| 193 | VASP | VASP | 1.08 | 1.43e-10 | cell surface | structural | Borderline | 0.4228 | A: cGMP-PKG / natriuretic signaling ; D: Immune / inflammatory co-activation ; D: Developmental pathway participation ; E: Constitutive cytoskeletal scaffold ; E: Focal adhesion / integrin structural |
| 194 | FAAH | FAAH | 1.86 | 1.12e-13 | intracellular | structural | Borderline | 0.4221 | D: Multi-system metabolic membership ; E: Constitutive cytoskeletal scaffold |

| Rank | Sheep symbol | Human ortholog | Mean log2FC | Min adj. P | Compartment | Class | Stability | Priority | Stability-pillar terms fired (pathway membership) |
|---|---|---|---|---|---|---|---|---|---|
| 195 | VXN | VXN | 3.32 | 1.94e-14 | cell surface | other | Lower | 0.4180 | |
| 196 | AGTR1 | AGTR1 | 2.17 | 1.16e-17 | cell surface | regulatory | Lower | 0.4151 | A: RAAS / angiotensin axis ; A: cAMP / PKA adrenergic signaling ; A: Ca2+ / calmodulin-CaMK axis ; A: cGMP-PKG / natriuretic signaling ; A: GPCR / G-protein second messenger ; C: Receptor / ligand-dependent activity ; C: Secreted / extracellularly released ; C: Membrane trafficking / endocytosis ; D: Cardiac-specific pathway member ; E: Sarcomere / myofilament terminal effector |
| 197 | THSD7B | THSD7B | 4.58 | 8.20e-21 | cell surface | other | Lower | 0.4122 | B: Post-translational protein modification ; B: Ubiquitin / proteasomal degradation ; D: Multi-system metabolic membership ; D: Multi-disease pathway involvement |
| 198 | SLIT3 | SLIT3 | 1.75 | 1.17e-13 | cell surface | ecm | Lower | 0.4108 | B: Negative transcriptional repression ; D: Developmental pathway participation |
| 199 | KIF21A | KIF21A | 1.42 | 1.99e-17 | intracellular | structural | Borderline | 0.4043 | C: Secreted / extracellularly released ; C: Membrane trafficking / endocytosis ; D: Hemostasis / vascular cross-pathway ; E: Constitutive cytoskeletal scaffold ; E: Focal adhesion / integrin structural |
| 200 | LTA4H | LTA4H | 1.49 | 3.44e-15 | cell surface | metabolic | Lower | 0.4042 | A: JAK-STAT / cytokine cross-activation ; D: Multi-system metabolic membership ; D: Immune / inflammatory co-activation |
| 201 | MDGA2 | MDGA2 | 2.92 | 7.20e-16 | cell surface | other | Lower | 0.4038 | B: Post-translational protein modification ; B: Ubiquitin / proteasomal degradation ; D: Multi-system metabolic membership |
| 202 | LGI4 | LGI4 | 1.26 | 3.00e-08 | cell surface | ecm | Borderline | 0.4032 | D: Developmental pathway participation |
| 203 | JAK1 | JAK1 | 1.31 | 8.55e-30 | cell surface | regulatory | Lower | 0.4012 | A: PI3K-Akt-mTOR cascade ; A: JAK-STAT / cytokine cross-activation ; B: RNA pol II transcriptional regulation ; B: Post-translational protein modification ; C: Receptor / ligand-dependent activity ; C: Kinase phosphorylation substrate ; D: Immune / inflammatory co-activation |
| 204 | FNDC4 | FNDC4 | 1.73 | 1.11e-16 | cell surface | regulatory | Lower | 0.4007 | C: Secreted / extracellularly released |
| 205 | IRX5 | IRX5 | 1.78 | 1.96e-07 | intracellular | regulatory | Borderline | 0.3957 | B: RNA pol II transcriptional regulation ; E: Lineage-locking transcription factor |
| 206 | RNF208 | RNF208 | 2.18 | 2.63e-06 | intracellular | metabolic | Borderline | 0.3956 | B: Ubiquitin / proteasomal degradation |
| 207 | CAMK1 | CAMK1 | 2.75 | 1.95e-32 | intracellular | regulatory | Lower | 0.3888 | A: Growth factor / RTK signaling ; A: RAAS / angiotensin axis ; A: Ca2+ / calmodulin-CaMK axis ; B: Post-translational protein modification ; C: Kinase phosphorylation substrate ; C: Ca2+-gated / calmodulin-dependent activity ; D: Cardiac-specific pathway member ; D: Neuronal system co-annotation |
| 208 | ATP6V0E1 | ATP6V0E1 | 1.12 | 3.99e-17 | intracellular | metabolic | Borderline | 0.3878 | A: JAK-STAT / cytokine cross-activation ; B: Cellular stress response ; D: Neuronal system co-annotation ; D: Immune / inflammatory co-activation ; D: Developmental pathway participation |
| 209 | SLC7A2 | SLC7A2 | 2.83 | 1.42e-23 | cell surface | metabolic | Lower | 0.3866 | |
| 210 | ENTREP2 | ENTREP2 | 1.90 | 2.24e-12 | unknown | other | Borderline | 0.3862 | |
| 211 | MRPS6 | MRPS6 | 1.54 | 8.97e-11 | intracellular | structural | Borderline | 0.3810 | B: Ubiquitin / proteasomal degradation ; D: Multi-system metabolic membership |
| 212 | PAWR | PAWR | 1.47 | 1.24e-11 | cell surface | regulatory | Lower | 0.3741 | B: RNA pol II transcriptional regulation ; B: Negative transcriptional repression ; C: Conditional complex assembly ; E: Constitutive cytoskeletal scaffold |
| 213 | BEX5 | BEX5 | 3.06 | 2.66e-11 | intracellular | regulatory | Lower | 0.3685 | C: Receptor / ligand-dependent activity |
| 214 | RCAN1 | RCAN1 | 1.67 | 3.49e-09 | intracellular | regulatory | Borderline | 0.3620 | A: Growth factor / RTK signaling ; B: miRNA-mediated gene suppression |
| 215 | TMTC1 | TMTC1 | 2.42 | 2.91e-08 | intracellular | structural | Lower | 0.3560 | |
| 216 | ASAP1 | ASAP1 | 2.06 | 2.82e-15 | cell surface | structural | Lower | 0.3555 | C: GTPase-regulated activity ; C: Membrane trafficking / endocytosis ; C: Conditional complex assembly |
| 217 | SNRPN | SNRPN | 1.17 | 3.90e-11 | intracellular | other | Borderline | 0.3533 | D: Multi-system metabolic membership |
| 218 | NPPA | NPPA | 2.33 | 0.00 | cell surface | regulatory | Borderline | 0.3532 | A: JAK-STAT / cytokine cross-activation ; A: cAMP / PKA adrenergic signaling ; A: cGMP-PKG / natriuretic signaling ; B: miRNA-mediated gene suppression ; B: RNA pol II transcriptional regulation ; B: Ubiquitin / proteasomal degradation ; C: Receptor / ligand-dependent activity ; C: Secreted / extracellularly released ; D: Cardiac-specific pathway member ; D: Multi-system metabolic membership ; D: Immune / inflammatory co-activation ; E: Sarcomere / myofilament terminal effector ; E: Conduction hardware (channel/pump) |
| 219 | CDC42EP3 | CDC42EP3 | 1.32 | 1.56e-12 | cell surface | structural | Lower | 0.3527 | A: Rho GTPase / cytoskeletal remodeling ; C: GTPase-regulated activity ; C: Conditional complex assembly ; E: Constitutive cytoskeletal scaffold |
| 220 | CASS4 | CASS4 | 1.89 | 1.11e-09 | cell surface | regulatory | Lower | 0.3457 | C: Kinase phosphorylation substrate ; E: Constitutive cytoskeletal scaffold |

| Rank | Sheep symbol | Human ortholog | Mean log2FC | Min adj. P | Compartment | Class | Stability | Priority | Stability-pillar terms fired (pathway membership) |
|---|---|---|---|---|---|---|---|---|---|
| 221 | GNAO1 | GNAO1 | 1.61 | 6.91e-17 | cell surface | regulatory | Lower | 0.3432 | A: Wnt/Notch developmental signaling ; A: Ca2+ / calmodulin-CaMK axis ; A: GPCR / G-protein second messenger ; C: Receptor / ligand-dependent activity ; C: GTPase-regulated activity ; D: Cardiac-specific pathway member ; E: Sarcomere / myofilament terminal effector |
| 222 | WFS1 | WFS1 | 2.25 | 5.88e-17 | intracellular | regulatory | Lower | 0.3352 | A: Ca2+ / calmodulin-CaMK axis ; B: RNA pol II transcriptional regulation ; B: Negative transcriptional repression ; B: Post-translational protein modification ; B: Ubiquitin / proteasomal degradation ; B: Cellular stress response ; C: Ca2+-gated / calmodulin-dependent activity ; D: Multi-system metabolic membership |
| 223 | CASQ1 | CASQ1 | 2.60 | 1.62e-08 | intracellular | metabolic | Lower | 0.3347 | D: Cardiac-specific pathway member ; E: Sarcomere / myofilament terminal effector ; E: Conduction hardware (channel/pump) |
| 224 | SSX2IP | SSX2IP | 1.85 | 3.54e-13 | intracellular | structural | Lower | 0.3337 | C: Conditional complex assembly ; E: Constitutive cytoskeletal scaffold |
| 225 | NUDCD3 | NUDCD3 | 1.12 | 6.59e-10 | intracellular | other | Borderline | 0.3304 | C: Conditional complex assembly |
| 226 | WIPI1 | WIPI1 | 1.38 | 6.46e-10 | intracellular | structural | Lower | 0.3249 | B: Autophagy / lysosomal turnover ; B: Cellular stress response ; C: Receptor / ligand-dependent activity ; C: Conditional complex assembly ; D: Multi-disease pathway involvement ; E: Constitutive cytoskeletal scaffold |
| 227 | COLQ | COLQ | 1.87 | 1.49e-12 | cell surface | structural | Lower | 0.3225 | C: Conditional complex assembly |
| 228 | RITA1 | RITA1 | 1.39 | 1.47e-13 | intracellular | structural | Lower | 0.3211 | A: Wnt/Notch developmental signaling ; B: RNA pol II transcriptional regulation ; B: Negative transcriptional repression ; E: Constitutive cytoskeletal scaffold |
| 229 | PGBD5 | PGBD5 | 2.05 | 1.48e-10 | intracellular | metabolic | Lower | 0.3186 | |
| 230 | H2BC15 | H2BC15 | 2.23 | 2.37e-16 | cell surface | structural | Lower | 0.3180 | B: Chromatin / epigenetic remodeling ; D: Immune / inflammatory co-activation ; D: Developmental pathway participation |
| 231 | DZANK1 | DZANK1 | 1.57 | 1.41e-08 | intracellular | structural | Borderline | 0.3179 | E: Constitutive cytoskeletal scaffold |
| 232 | ADGRA3 | ADGRA3 | 1.27 | 3.75e-11 | cell surface | regulatory | Lower | 0.3178 | A: GPCR / G-protein second messenger ; C: Receptor / ligand-dependent activity |
| 233 | FHIT | FHIT | 1.86 | 4.83e-09 | cell surface | metabolic | Lower | 0.3156 | B: Ubiquitin / proteasomal degradation ; D: Multi-system metabolic membership |
| 234 | RPP38 | RPP38 | 1.79 | 7.45e-12 | intracellular | other | Lower | 0.3140 | D: Multi-system metabolic membership |
| 235 | PTPRF | PTPRF | 1.76 | 3.33e-07 | cell surface | regulatory | Lower | 0.3110 | A: Growth factor / RTK signaling ; D: Neuronal system co-annotation |
| 235 | PCDH7 | PCDH7 | 1.45 | 5.13e-09 | cell surface | structural | Lower | 0.3110 | A: Rho GTPase / cytoskeletal remodeling ; C: GTPase-regulated activity ; D: Hemostasis / vascular cross-pathway ; E: Focal adhesion / integrin structural |
| 237 | PHOSPHO1 | PHOSPHO1 | 1.79 | 4.17e-09 | cell surface | structural | Lower | 0.3028 | D: Multi-system metabolic membership ; D: Multi-disease pathway involvement |
| 238 | TRAM2 | TRAM2 | 2.16 | 4.33e-10 | intracellular | structural | Lower | 0.3023 | |
| 239 | PIK3R3 | PIK3R3 | 1.75 | 2.03e-19 | intracellular | regulatory | Lower | 0.3005 | A: MAPK/ERK signaling cascade ; A: PI3K-Akt-mTOR cascade ; A: JAK-STAT / cytokine cross-activation ; A: FoxO / apoptotic stress axis ; A: Growth factor / RTK signaling ; A: cAMP / PKA adrenergic signaling ; B: miRNA-mediated gene suppression ; D: Multi-system metabolic membership ; D: Immune / inflammatory co-activation ; D: Developmental pathway participation ; D: Hemostasis / vascular cross-pathway ; E: Focal adhesion / integrin structural |
| 240 | STARD3NL | STARD3NL | 1.26 | 3.73e-12 | intracellular | regulatory | Lower | 0.2940 | D: Multi-system metabolic membership |
| 241 | ALDH18A1 | ALDH18A1 | 1.14 | 1.39e-10 | intracellular | metabolic | Lower | 0.2902 | D: Multi-system metabolic membership |
| 242 | DIPK1B | DIPK1B | 1.29 | 2.63e-07 | intracellular | regulatory | Borderline | 0.2877 | |
| 243 | RAB38 | RAB38 | 2.48 | 5.21e-12 | cell surface | regulatory | Lower | 0.2717 | B: Post-translational protein modification ; B: Ubiquitin / proteasomal degradation ; C: GTPase-regulated activity ; C: Secreted / extracellularly released ; C: Membrane trafficking / endocytosis ; C: Conditional complex assembly ; D: Multi-system metabolic membership |
| 244 | TIE1 | TIE1 | 1.96 | 1.23e-08 | cell surface | regulatory | Lower | 0.2648 | C: Kinase phosphorylation substrate |
| 245 | POMGNT2 | POMGNT2 | 1.51 | 7.27e-16 | intracellular | metabolic | Lower | 0.2575 | B: Post-translational protein modification ; B: Ubiquitin / proteasomal degradation ; D: Multi-system metabolic membership |
| 246 | LGMN | LGMN | 1.55 | 1.32e-08 | cell surface | regulatory | Lower | 0.2512 | A: JAK-STAT / cytokine cross-activation ; B: Autophagy / lysosomal turnover ; D: Multi-system metabolic membership ; D: Immune / inflammatory co-activation |
| 247 | GALM | GALM | 1.35 | 3.51e-11 | cell surface | metabolic | Lower | 0.2483 | D: Multi-system metabolic membership ; D: Multi-disease pathway involvement |
| 248 | MSI2 | MSI2 | 1.38 | 1.89e-09 | intracellular | regulatory | Lower | 0.2475 | A: Rho GTPase / cytoskeletal remodeling ; C: GTPase-regulated activity |
| 249 | NAP1L5 | NAP1L5 | 1.61 | 9.50e-10 | intracellular | other | Lower | 0.2463 | |

| Rank | Sheep symbol | Human ortholog | Mean log2FC | Min adj. P | Compartment | Class | Stability | Priority | Stability-pillar terms fired (pathway membership) |
|---|---|---|---|---|---|---|---|---|---|
| 250 | KCNMB4 | KCNMB4 | 2.12 | 5.81e-04 | cell surface | regulatory | Lower | 0.2428 | A: cGMP-PKG / natriuretic signaling ; D: Neuronal system co-annotation ; D: Hemostasis / vascular cross-pathway ; E: Sarcomere / myofilament terminal effector ; E: Conduction hardware (channel/pump) ; E: Focal adhesion / integrin structural |
| 251 | MAP2K1 | MAP2K1 | 1.49 | 1.01e-10 | intracellular | regulatory | Lower | 0.2365 | A: MAPK/ERK signaling cascade ; A: JAK-STAT / cytokine cross-activation ; A: FoxO / apoptotic stress axis ; A: Growth factor / RTK signaling ; A: cAMP / PKA adrenergic signaling ; A: cGMP-PKG / natriuretic signaling ; B: miRNA-mediated gene suppression ; B: Negative transcriptional repression ; C: Kinase phosphorylation substrate ; D: Immune / inflammatory co-activation ; D: Developmental pathway participation ; D: Multi-disease pathway involvement ; E: Sarcomere / myofilament terminal effector ; E: Focal adhesion / integrin structural |
| 252 | KIF22 | KIF22 | 1.53 | 6.78e-08 | intracellular | structural | Lower | 0.2246 | C: Secreted / extracellularly released ; C: Membrane trafficking / endocytosis ; D: Immune / inflammatory co-activation ; D: Hemostasis / vascular cross-pathway ; E: Constitutive cytoskeletal scaffold ; E: Focal adhesion / integrin structural |
| 253 | PBX3 | PBX3 | 1.31 | 4.79e-09 | intracellular | regulatory | Lower | 0.2188 | A: TGF-beta / BMP morphogen axis ; B: RNA pol II transcriptional regulation ; D: Developmental pathway participation ; E: Lineage-locking transcription factor |
| 254 | TMEM163 | TMEM163 | 2.06 | 9.74e-08 | cell surface | regulatory | Lower | 0.2102 | |
| 255 | MSANTD3 | MSANTD3 | 1.19 | 7.65e-09 | unknown | other | Lower | 0.2100 | |
| 256 | BICD1 | BICD1 | 1.59 | 7.87e-09 | intracellular | structural | Lower | 0.2078 | C: Receptor / ligand-dependent activity ; C: Secreted / extracellularly released ; C: Membrane trafficking / endocytosis ; C: Conditional complex assembly ; E: Constitutive cytoskeletal scaffold |
| 257 | TMEM40 | TMEM40 | 2.49 | 1.38e-07 | intracellular | other | Lower | 0.2015 | |
| 258 | RAB33A | RAB33A | 2.14 | 6.06e-18 | cell surface | regulatory | Lower | 0.1996 | B: Post-translational protein modification ; B: Ubiquitin / proteasomal degradation ; C: GTPase-regulated activity ; C: Secreted / extracellularly released ; C: Membrane trafficking / endocytosis ; C: Conditional complex assembly ; D: Multi-system metabolic membership |
| 259 | B3GALNT1 | B3GALNT1 | 2.15 | 1.76e-08 | intracellular | metabolic | Lower | 0.1988 | D: Multi-system metabolic membership |
| 260 | ISYNA1 | ISYNA1 | 1.37 | 8.67e-09 | intracellular | metabolic | Lower | 0.1983 | D: Multi-system metabolic membership |
| 261 | MAPK13 | MAPK13 | 2.80 | 7.97e-07 | cell surface | regulatory | Lower | 0.1976 | A: MAPK/ERK signaling cascade ; A: JAK-STAT / cytokine cross-activation ; A: FoxO / apoptotic stress axis ; A: cAMP / PKA adrenergic signaling ; C: Kinase phosphorylation substrate ; D: Cardiac-specific pathway member ; D: Immune / inflammatory co-activation |
| 262 | IGF1R | IGF1R | 1.31 | 7.78e-09 | cell surface | regulatory | Lower | 0.1920 | A: MAPK/ERK signaling cascade ; A: PI3K-Akt-mTOR cascade ; A: FoxO / apoptotic stress axis ; A: Growth factor / RTK signaling ; B: miRNA-mediated gene suppression ; B: Autophagy / lysosomal turnover ; C: Receptor / ligand-dependent activity ; C: Kinase phosphorylation substrate ; C: Membrane trafficking / endocytosis ; C: Conditional complex assembly ; D: Multi-disease pathway involvement ; E: Constitutive cytoskeletal scaffold ; E: Focal adhesion / integrin structural |
| 263 | TOMM34 | TOMM34 | 1.49 | 4.19e-08 | intracellular | regulatory | Lower | 0.1913 | |
| 264 | SNX10 | SNX10 | 1.16 | 7.91e-06 | intracellular | structural | Lower | 0.1850 | E: Constitutive cytoskeletal scaffold |
| 265 | SHISA2 | SHISA2 | 1.97 | 2.30e-06 | intracellular | regulatory | Lower | 0.1837 | A: Wnt/Notch developmental signaling |
| 266 | TMEM108 | TMEM108 | 1.20 | 6.20e-07 | intracellular | regulatory | Lower | 0.1794 | |
| 267 | ACY1 | ACY1 | 1.35 | 1.05e-07 | cell surface | metabolic | Lower | 0.1791 | D: Multi-system metabolic membership ; D: Multi-disease pathway involvement |
| 268 | DTNB | DTNB | 1.19 | 1.10e-06 | cell surface | regulatory | Lower | 0.1735 | |
| 269 | KIF26B | KIF26B | 1.91 | 1.11e-04 | intracellular | structural | Lower | 0.1711 | C: Secreted / extracellularly released ; C: Membrane trafficking / endocytosis ; D: Hemostasis / vascular cross-pathway ; E: Constitutive cytoskeletal scaffold ; E: Focal adhesion / integrin structural |
| 270 | CDKN1A | CDKN1A | 1.38 | 1.40e-07 | intracellular | regulatory | Lower | 0.1711 | A: PI3K-Akt-mTOR cascade ; A: JAK-STAT / cytokine cross-activation ; A: FoxO / apoptotic stress axis ; A: Growth factor / RTK signaling ; B: RNA pol II transcriptional regulation ; C: Kinase phosphorylation substrate ; D: Immune / inflammatory co-activation ; D: Developmental pathway participation ; D: Multi-disease pathway involvement |
| 271 | SLC29A2 | SLC29A2 | 1.22 | 0.00 | cell surface | metabolic | Lower | 0.1588 | |
| 272 | BBS5 | BBS5 | 1.38 | 4.07e-06 | cell surface | structural | Lower | 0.1573 | C: Conditional complex assembly ; E: Constitutive cytoskeletal scaffold |
| 273 | CAB39L | CAB39L | 1.39 | 2.01e-06 | intracellular | regulatory | Lower | 0.1563 | A: PI3K-Akt-mTOR cascade ; C: Kinase phosphorylation substrate |

| Rank | Sheep symbol | Human ortholog | Mean log2FC | Min adj. P | Compartment | Class | Stability | Priority | Stability-pillar terms fired (pathway membership) |
|---|---|---|---|---|---|---|---|---|---|
| 274 | ZDHHC14 | ZDHHC14 | 1.48 | 2.24e-07 | intracellular | metabolic | Lower | 0.1504 | |
| 275 | TJP2 | TJP2 | 1.29 | 7.29e-08 | cell surface | structural | Lower | 0.1472 | C: Kinase phosphorylation substrate |
| 276 | TMEM53 | TMEM53 | 1.21 | 2.93e-07 | intracellular | regulatory | Lower | 0.1446 | |
| 277 | TMC4 | TMC4 | 1.46 | 1.27e-05 | cell surface | metabolic | Lower | 0.1336 | |
| 278 | TTLL7 | TTLL7 | 1.07 | 1.78e-07 | intracellular | structural | Lower | 0.1267 | B: Post-translational protein modification ; B: Ubiquitin / proteasomal degradation ; D: Multi-system metabolic membership ; E: Constitutive cytoskeletal scaffold |
| 279 | C1orf53 | C1orf53 | 1.41 | 7.80e-07 | unknown | other | Lower | 0.1197 | |
| 280 | RGS2 | RGS2 | 1.69 | 9.46e-04 | cell surface | regulatory | Lower | 0.1186 | A: cAMP / PKA adrenergic signaling ; A: Ca2+ / calmodulin-CaMK axis ; A: cGMP-PKG / natriuretic signaling ; A: GPCR / G-protein second messenger ; C: GTPase-regulated activity ; C: Ca2+-gated / calmodulin-dependent activity ; D: Cardiac-specific pathway member |
| 281 | ANKRD6 | ANKRD6 | 1.36 | 3.58e-04 | intracellular | regulatory | Lower | 0.0969 | A: Wnt/Notch developmental signaling |
| 282 | NABP1 | NABP1 | 1.06 | 1.58e-04 | intracellular | other | Lower | 0.0927 | B: RNA pol II transcriptional regulation |
| 283 | E2F3 | E2F3 | 1.32 | 6.19e-06 | intracellular | regulatory | Lower | 0.0906 | A: Wnt/Notch developmental signaling ; A: FoxO / apoptotic stress axis ; B: RNA pol II transcriptional regulation ; D: Multi-disease pathway involvement ; E: Lineage-locking transcription factor |
| 284 | INPP4B | INPP4B | 1.49 | 2.53e-04 | intracellular | regulatory | Lower | 0.0877 | D: Multi-system metabolic membership |
| 285 | FAM169A | FAM169A | 1.56 | 5.92e-04 | intracellular | other | Lower | 0.0814 | A: Rho GTPase / cytoskeletal remodeling ; C: GTPase-regulated activity |
| 286 | ST6GALNAC4 | ST6GALNAC4 | 1.25 | 8.53e-05 | intracellular | metabolic | Lower | 0.0774 | B: Ubiquitin / proteasomal degradation ; D: Multi-system metabolic membership ; D: Multi-disease pathway involvement |

*Table S9. The 286 Purkinje-fiber-enriched genes carrying a mappable human ortholog, ranked by final priority score. Mean log2FC, mean of the |log2 fold-change| versus cardiomyocytes and versus stroma (positive = higher in Purkinje fibers); Min adj. P, smallest Benjamini-Hochberg-adjusted P across the two comparisons; Compartment, predicted subcellular localization (accessibility weight); Class, functional class; Stability, composite stability tier; Priority, final priority score (0.55 x stability percentile + 0.35 x specificity + 0.10 x compartment weight). The final column lists the stability pillars and terms a gene fired (its pathway membership). Sign convention: enrichment in Purkinje fibers corresponds to a positive mean log2FC.*

#### Table S10. RT-qPCR results (human; Purkinje fibers vs cardiomyocytes, n = 4)

| Gene | n (PF/CM) | Mean ΔΔCt | Fold-change (95% CI) | Direction | Welch t (df) | Welch P | MWU P | Call |
|---|---|---|---|---|---|---|---|---|
| *MYL4* | 4 / 4 | -4.16 | 17.9 (6.4-49.8) | PF | 6.91 (6.0) | 0.0005 | 0.0286 | PF-enriched |
| *GJA5 (Cx40)* | 4 / 4 | -4.65 | 25.1 (1.2-545.1) | PF | 2.81 (4.4) | 0.0435 | 0.0571 | PF-enriched |
| *CNTN5* | 4 / 4 | -3.95 | 15.4 (2.0-120.3) | PF | 3.27 (5.9) | 0.0174 | 0.0571 | PF-enriched |
| *TAGLN* | 4 / 4 | -2.08 | 4.2 (1.1-16.5) | PF | 3.05 (3.6) | 0.0434 | 0.0571 | PF-enriched |
| *CNTN2* | 4 / 4 | -2.17 | 4.5 (0.8-24.3) | PF | 2.66 (3.4) | 0.0667 | 0.0286 | Trend (PF) |
| *CAPN3* | 4 / 4 | +1.22 | 2.3 (1.3-4.2) | CM | 3.58 (5.7) | 0.0127 | 0.0286 | CM-enriched |
| *CNN1* | 4 / 4 | +0.44 | 1.4 (0.4-5.0) | CM | 0.58 (5.8) | 0.5851 | 0.8857 | Not enriched |
| *PTGR1* | 4 / 4 | -0.05 | 1.0 (0.6-1.8) | PF | 0.16 (5.2) | 0.8763 | 1.0000 | Not enriched |
| *SCN4B* | 4 / 4 | -0.04 | 1.0 (0.4-2.6) | PF | 0.08 (3.7) | 0.9430 | 0.6857 | Not enriched |
| *CNTN4* | 4 / 4 | -0.50 | 1.4 (0.5-4.4) | PF | 0.86 (3.9) | 0.4382 | 0.4857 | Not enriched |
| *RIMS1* | 4 / 4 | -1.01 | 2.0 (0.2-17.5) | PF | 0.81 (5.3) | 0.4501 | 0.4857 | Not enriched |
| *TES* | 4 / 4 | +0.27 | 1.2 (0.4-3.5) | CM | 0.44 (5.0) | 0.6767 | 1.0000 | Not enriched |

*Table S10. Full RT-qPCR statistics for candidate-marker conservation in human myocardium (free-running Purkinje fibers, PF, versus left-ventricular cardiomyocytes, CM; corresponds to Figure 8). Relative expression was normalized to the geometric mean of two reference genes (b2m and GUSB). Primary test: Welch's unequal-variance t test on ΔCt (two-sided P; Welch P); sensitivity test: two-tailed exact Mann-Whitney U test (two-sided P; MWU P). Mean ΔΔCt, Fold-change (95% CI) and Welch t (df) are defined as in Table S7. n = 4 donors per group. Significance threshold P < 0.05; no correction for multiplicity. All twelve genes were recomputed from the raw Cq export and reproduce the reported values.*

#### Figure S3. Per-pillar heatmap ranking the top fifty candidate genes

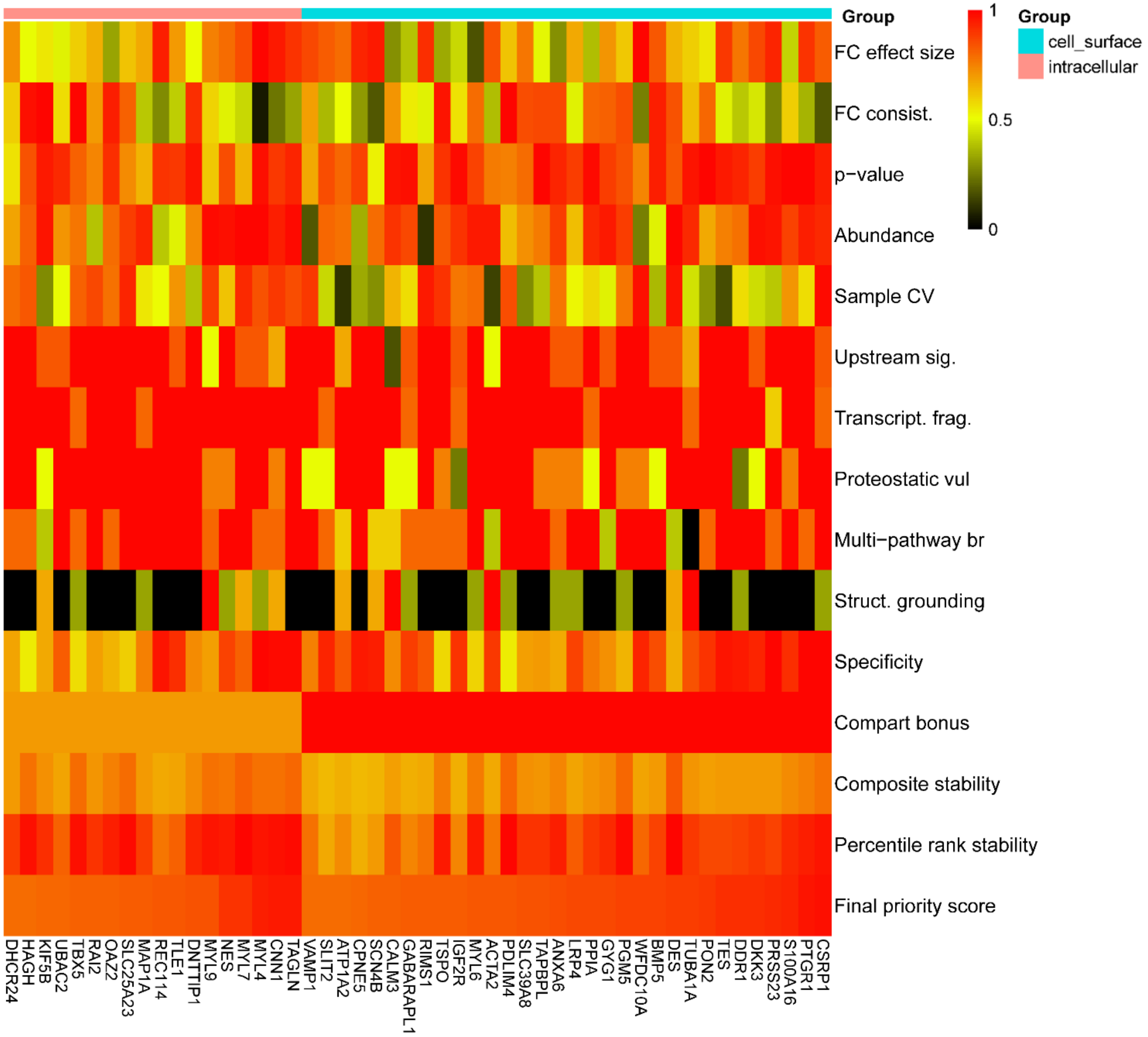


*Figure S3. Pillar-level scoring of the fifty top-ranked candidate Purkinje-fiber markers. Heatmap of the 50 highest-priority genes (rows), separated into intramural/intracellular and cell-surface groups and ordered by composite priority score within each group. Color encodes each gene's score for the individual scoring pillars (columns), with warmer colors denoting higher pillar scores.*
